# Click Chemistry-Based Strategy for Modular Ligand Attachment to siRNAs: Toward Extrahepatic RNAi

**DOI:** 10.64898/2026.05.21.726808

**Authors:** Julia A. Rädler, Eliza Filipiak, Antonin Marquant, Miina Ojansivu, Tomasz Czapik, Alyssa Hill, Nina Ahlskog, Samantha Roudi, Cristiana Barradas, Yanjie Huang, Osama Saher, Matthew Wood, Rula Zain, Malgorzata Honcharenko, Samir EL Andaloussi

**Affiliations:** Biomolecular and Cellular Medicine, Department of Laboratory Medicine, Karolinska Institutet, Huddinge, Stockholm, 141 52, Sweden; Karolinska ATMP Center, Karolinska Institutet, Huddinge, Stockholm, 141 52, Sweden; Department of Cellular Therapy and Allogeneic Stem Cell Transplantation (CAST), Karolinska University Hospital, Huddinge, Stockholm, 141 52, Sweden; Institute of Developmental and Regenerative Medicine, Department of Paediatrics, University of Oxford, Oxford, OX3 7TY, United Kingdom; Center for Rare Diseases, Clinical Genetics and Genomics, Karolinska University Hospital, Solna, Stockholm, 171 76, Sweden

## Abstract

Efficient extrahepatic delivery of siRNAs remains a major limitation for broadening their therapeutic potential. Using a modular, orthogonal click chemistry platform, we generated 28 siRNA conjugates varying in ligand class, valency, and spatial arrangement. Following systemic administration, fatty acid conjugates – particularly palmitic acid (C16) – outperformed sterol- and phospholipid-based designs in promoting extrahepatic gene silencing, with preferential activity observed in heart and skeletal muscle. Increasing ligand valency through 3′,5′-bis-conjugation generally enhanced activity compared to 5’-mono conjugation. Nevertheless, bis-C22 conjugates showed increased hepatic activity, suggesting a shift in tissue distribution linked to hydrophobicity. Architectural parameters further modulated outcomes: Branched 5′ C16 conjugates, bearing two lipids on one terminus, were markedly less active than their bis counterparts and required short PEG spacers to restore activity. Notably, bis-lipid conjugation strategies that enhanced extrahepatic activity for an siRNA did not translate to an ASO gapmer, underscoring modality-specific constraints. Together, these findings delineate structure-activity relationships and establish bis-fatty-acid conjugation as a robust design principle for achieving extrahepatic RNAi.

**GRAPHICAL ABSTRACT:** 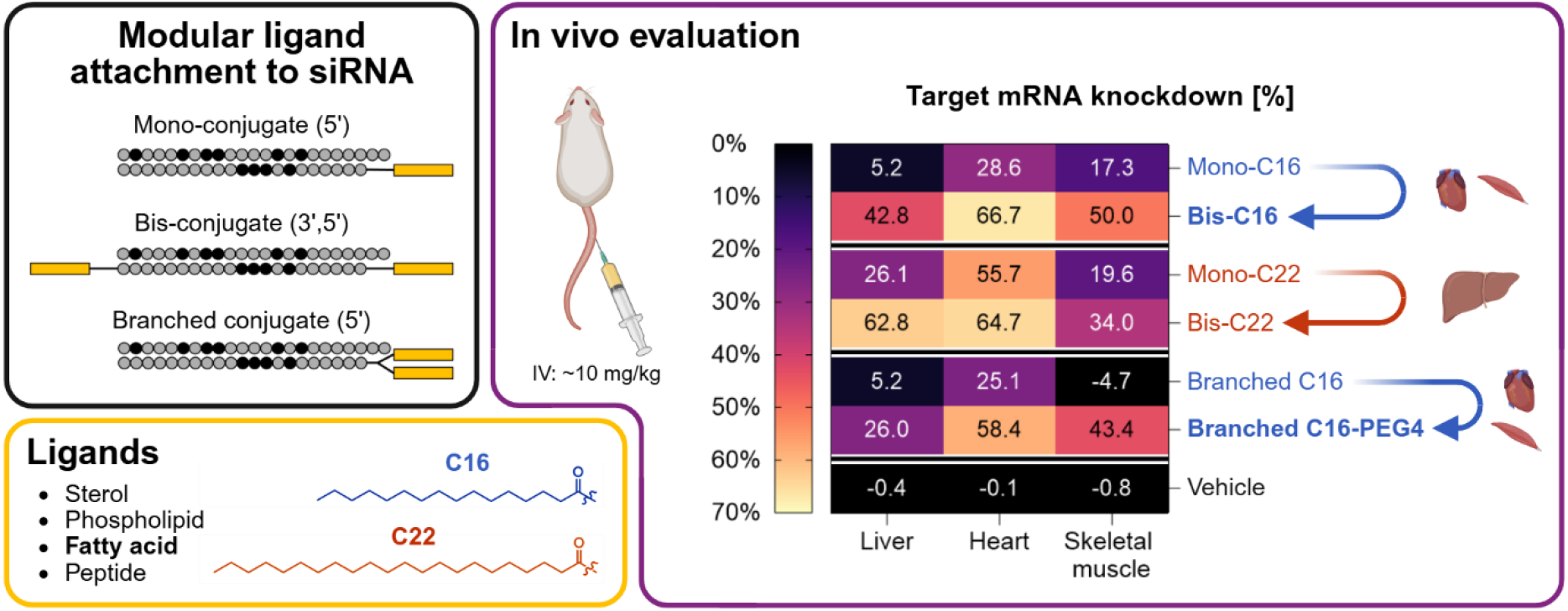

## INTRODUCTION

Oligonucleotides, including small interfering RNAs (siRNAs) and antisense oligonucleotides (ASOs), hold enormous therapeutic potential owing to their ability to modulate the expression of essentially any gene in a highly specific manner. Their rational design, combined with their long duration of action renders oligonucleotides an attractive therapeutic modality. Major advances in chemical stabilization and structural optimization of the siRNA and ASO scaffolds have been instrumental in improving in vivo stability, efficacy, and tolerability, thereby greatly facilitating clinical translation (1, 2). Despite these advances, the vast majority of targets remain inaccessible, as certain physicochemical properties of oligonucleotides, such as their size, anionic charge, and hydrophilicity, hamper productive delivery.

The conjugation of ligands to siRNAs has proven highly effective in improving their pharmacokinetic properties and aiding delivery. The leading example is N-acetylgalactosamine (GalNAc), which enables efficient delivery to hepatocytes via the asialoglycoprotein receptor (ASGPR) (3) and has paved the way for the clinical approval of multiple siRNA and ASO therapeutics (4). However, extrahepatic delivery remains challenging, which significantly restricts the therapeutic scope of systemically administered oligonucleotide drugs.

To date, a broad spectrum of ligands has been explored for conjugate-mediated delivery beyond the liver. Among these, lipid-based ligands are one of the most extensively studied. Their hydrophobicity promotes interactions with plasma proteins, enabling conjugates to “hitchhike” on endogenous carriers such as albumin and lipoproteins to modulate biodistribution, extravasation, and tissue access. Cholesterol-conjugated siRNAs were the first lipid conjugates shown to enable systemic delivery in vivo (5), but their strong association with low-density lipoprotein (LDL) largely restricts them to LDLR-rich tissues such as liver and adrenal gland (6). In contrast, fatty acids engage a wider range of plasma proteins, including albumin, thereby supporting broader (extrahepatic) tissue delivery (7, 8). In this way, potent skeletal and cardiac muscle silencing has been achieved with siRNA conjugated to docosanoic acid, a saturated C22 fatty acid (9, 10), and the albumin-mediated interstitial access provided by C16 for ASOs (11, 12). This principle extends to divalent C18 structures and dendritic amphiphiles (i.e., incorporating four C12 chains), which leverage high-affinity albumin binding to support systemic delivery to tumors and other extrahepatic tissues such as heart, fat and lung (13–17). These findings highlight the potential of lipid conjugation to expand oligonucleotide distribution beyond the liver. However, it has been shown that many design variables such as ligand class, valency, spatial arrangement, as well as head group and linker chemistry influence the pharmacokinetic properties of oligonucleotide conjugates, emphasizing the need for modular platforms that facilitate a systematic exploration of the conjugate space.

To further explore the conjugate design space for siRNA delivery, we used a modular click chemistry platform that supports rapid and controlled assembly of chemically defined siRNA conjugates. This approach enabled a systematic evaluation of multiple ligand classes and architectures, with an emphasis on lipophilic conjugates. We generated and tested diverse configurations incorporating seven lipid-based ligands, two peptides, and one phospholipid, varying both valency and spatial arrangement to probe how these parameters shape in vivo activity. Across this design space, fatty-acid conjugates promoted extrahepatic RNAi, with 3′,5′-bis-conjugates consistently outperforming their 5′-mono counterparts. Notably, C16 emerged as a particularly effective ligand for heart and skeletal muscle delivery, achieving its highest potency as a 3′,5′-bis-conjugate or as a branched 5′ conjugate incorporating PEG4. In addition, bis-C16OOH, bis-C20OOH, and mono-C22 displayed the strongest heart-selective delivery profiles. Importantly, the benefits of bis-fatty-acid conjugation were specific to siRNA, as analogous designs did not translate to gapmer ASOs. Collectively, our findings provide a comparative framework for rational conjugate selection and design, highlighting how ligand identity, valency, and configuration govern productive gene silencing outside the liver.

## MATERIAL AND METHODS

### Materials

The organic solvents acetonitrile (ACN, High Performance Liquid Chromatography (HPLC) grade, Honeywell), dichloromethane (DCM, SigmaAldrich, Analytical Grade), dimethyl sulfoxide (DMSO, Merck) and N-methyl-2-pyrrolidone (NMP, SigmaAldrich) were of commercial grade and used as received. N,N-dimethylformamide (DMF, SigmaAldrich), tetrahydrofuran (THF, Acros Organics) and pyridine (Py, SigmaAldrich) were additionally dried over 4A molecular sieves. Modified oligonucleotides were synthesized and received from AstraZeneca. The following reagents were used for peptide synthesis: Fmoc protected amino acids (SigmaAldrich, Merck), N,N′-Diisopropylcarbodiimide (DIC, Iris Biotech), Oxyma (SigmaAldrich), lutidine (Thermoscientific), acetic anhydrate (SigmaAldrich), trifluoroacetic acid (TFA, SigmaAldrich), triisopropylsilane (TIS, SigmaAldrich) and methyl tert-butyl ether (SigmaAldrich) were used. The following were used as substrates for the reactions: palmitic acid N-Hydroxysuccinimide ester palmitic acid NHS ester, SigmaAldrich), Palmitic acid PEG4-N-hydroxysuccinimide ester (Palmitic acid PEG4-NHS, BroadPharm), azido-PEG3-amine (Lumiprobe), docosanoic acid (SigmaAldrich), hexafluorophosphate benzotriazole tetramethyl uronium (HBTU, SigmaAldrich), hydrochloric acid (HCl), trans-cyclooctene N-Hydroxysuccinimide (TCO-NHS ester, BroadPharm), and 1,2-dioleoyl-sn-glycero-3-phosphoethanolamine - dibenzocyclooctyne group (DOPE-DBCO, BroadPharm).

### HPLC and UPLC/MS

Reversed-phase HPLC was carried out on a Waters HPLC system using a Waters XBridge Prep C18 5 µm OBD 19×100mm column with 10 mL/min flow rate using detection at 265 nm and 280 nm. Two different buffer systems were used for oligonucleotides and peptides: For the purification of oligonucleotides and their conjugates, 50 mM triethylammonium acetate (TEAA) buffer in water with 10% ACN was used as a buffer A2 and ACN as a buffer B2. For the purification of peptides, 0.2% trifluoroacetic acid (TFA) in H_2_O was used as a buffer A1 and 0.2% TFA in ACN was used as a buffer B1. Mass spectrometry (electrospray time-of-flight, ES TOF-MS) experiments were performed on the Waters Xevo G2-S QTOF UPLC/mass system using an ACQUITY Premier BEH C18 1.7 μm 2.1 x 100 mm column. The following buffers were used: 5 mM ammonium acetate in water (A3) and 5 mM ammonium acetate in ACN (B3).

### Synthesis of Lipid and Peptide Ligands and Conjugation to Oligonucleotides

A modular panel of azide-functionalized lipid and peptide ligands was synthesized to enable site-selective bioorthogonal conjugation to modified Sod1 sense oligonucleotides. Lipid–PEG– azide derivatives based on palmitic (C16), docosanoic (C22), and lithocholic acid (LCA) scaffolds were prepared via amide coupling of activated carboxylic acids with azido-PEG amines. Branched lipid architectures and negatively charged variants were generated using analogous coupling and deprotection strategies.

Azide-modified peptides, including the cardiomyocyte-targeting peptide (CTP), Ahx-GWWG, and the cationic linker L4+, were synthesized by Fmoc solid-phase peptide synthesis. Selected peptides were further functionalized with methyltetrazine to enable orthogonal ligation chemistry.

Sod1 oligonucleotides bearing 5′- or 3′,5′-bicyclo[6.1.0]non-4-yne (BCN) modifications were conjugated to azide-functionalized ligands via strain-promoted azide–alkyne cycloaddition (SPAAC). Bis- and hetero-bifunctional 3′,5′-conjugates were obtained using oligonucleotides incorporating orthogonal BCN and trans-cyclooctene (TCO) handles, enabling sequential SPAAC and inverse-electron-demand Diels–Alder (IEDDA) reactions with methyltetrazine-modified ligands.

Detailed synthetic procedures and analytical data for all compounds are provided in the Supporting Information.

### siRNA synthesis

Based on a previously published Sod1 siRNA sequence (18), the sense and antisense strands were synthesized by Axolabs GmbH with the chemical modifications detailed in **Supplementary Table 1**.

### ASO synthesis

Fully phosphorothioate (PS)-modified LNA gapmer oligonucleotides were synthesized by automated solid-phase phosphoramidite chemistry on a K&A H-8 synthesizer (K&A Labs GmbH) at the 10 µmol scale using controlled pore glass (CPG) universal support. Standard β-cyanoethyl phosphoramidite chemistry was applied using commercially available DNA (Sigma-Aldrich) and LNA phosphoramidites (Glen Research). Phosphoramidites (0.1 M in anhydrous ACN) were activated with 0.25 M 5-(Ethylthio)-1H-Tetrazole (ETT); coupling times were 60 s for DNA monomers and 4 min for LNA monomers. All internucleotide linkages were converted to phosphorothioates by sulfurization with 0.1 M DDTT (3-((dimethylaminomethylene)amino)-3H-1,2,4-dithiazole-5-thione) in pyridine/acetonitrile, replacing the standard iodine oxidation step. Capping was performed using acetic anhydride/N-methylimidazole reagents according to standard protocols.

Following chain assembly, oligonucleotides were cleaved from the support and deprotected with concentrated aqueous ammonia at 55°C for 12–16 h. Crude products were purified by reversed-phase HPLC and desalted by size-exclusion chromatography. Identity was confirmed by LCMS-QToF mass spectrometry.

### Dissolution and annealing

Single-stranded siRNAs were dissolved to 1 mM in 90% DPBS (Gibco) and 10% DMSO (D4540, Sigma-Aldrich) (v/v). To ensure proper dissolution they were incubated at 60-70°C for 10 min. For annealing, sense and antisense strands were mixed at an equimolar ratio and incubated for 5 min at 95°C followed by gradual (1°C/min) cooling to RT. For in vivo experiments, the siRNAs were further diluted to 300 µM in DPBS resulting in a final concentration of 3% DMSO (v/v).

ASOs were dissolved in 400 µL of nuclease-free water and 1% DMSO (D4540, Sigma-Aldrich) (v/v). ASO concentrations were measured using a NanoDrop spectrophotometer. For in vivo experiments, ASOs were further diluted to 200 µM in a final formulation of H_2_O:DPBS:DMSO at a 63:36.37:0.63 v/v/v ratio.

### Critical aggregation concentration (CAC)

Serial dilutions of oligonucleotide conjugates, siRNA duplexes and ASOs, were prepared in a 96-well plate ranging from 50 µM to 100 nM in 75 μL of Ca^2+^/Mg^2+^ free DPBS with Nile Red (0.25 µg), and the plate was agitated at 37°C in the dark for 1.5 h. Fluorescence was measured on a CLARIOstar Plus plate fluorimeter at excitation 535 ± 10 nm and emission 612 ± 10 nm. The critical aggregation concentration (CAC) was defined as the intersection point of the two linear regions in the plot of Nile Red fluorescence versus oligonucleotide concentration.

### In vitro transfection

Neuro-2a cells (5 × 10^3^ cells/well) were seeded into 96-well tissue-culture plates and allowed to adhere overnight. The following day, siRNA conjugates (1 pmol/well) were transfected using Lipofectamine™ RNAiMAX (0.3 μL/well; 13778150, Thermo Fisher Scientific) in duplicate wells, following the manufacturer’s protocol. Cells were incubated for an additional 48 h prior to RNA isolation.

For RNA extraction, culture medium was completely removed, and plates were stored at −80°C until further processing. Total RNA was isolated with the Maxwell® RSC simplyRNA Cells Kit (AS1390, Promega) using the Maxwell® RSC Instrument according to the manufacturer’s instructions. RNA concentration was quantified using the QuantiFluor® RNA System (E3310, Promega) on a Quantus™ Fluorometer (Promega).

For cDNA synthesis, 100–200 ng of total RNA were reverse-transcribed using the High-Capacity cDNA Reverse Transcription Kit (43-688-13, Applied Biosystems™). Quantitative PCR was performed on a CFX Opus 96 Real-Time PCR System (Bio-Rad) using TaqMan™ Fast Advanced Master Mix (4444557, Applied Biosystems™) and TaqMan® Gene Expression Assays specific for Sod1 (Mm01344233_g1) and Gapdh (Mm99999915_g1). Each qPCR reaction contained 10 ng cDNA.

Gene expression levels were calculated using the ΔΔCt method, with Sod1 expression normalized to the endogenous control Gapdh, and further normalized to the untreated control group.

### Animal experiments

Animal experiments were conducted in Sweden and the United Kingdom and approved by the respective regulatory authorities. Experiments performed in Sweden (single-dose studies with siRNAs and ASOs) were approved by the Swedish Animal Ethics Committee in Linköping (permit number 13849-2020; 14772-2023; 2173-2021) under supervision of the Swedish Board of Agriculture and conducted in accordance with national legislation and EU Directive 2010/63/EU. Repeat-dose siRNA experiments were approved by the UK Home Office under the Animals (Scientific Procedures) Act 1986, Project License (PPL) no: PP6777529. All procedures were designed to minimize animal suffering and the number of animals used.

Female NMRI mice (4-5 weeks old or 20-25 g) were obtained from commercial vendors and maintained under specific pathogen-free conditions with ad libitum access to food and water under controlled temperature, humidity, and a 12 h light/dark cycle. Animals were acclimatized prior to experimentation, randomly assigned to treatment groups (n = 5), and monitored daily by trained animal care staff with veterinary supervision available when required.

For single-dose studies conducted in Sweden, mice were administered siRNAs (600 nmol/kg) or ASOs (1 µmol/kg) via intravenous tail-vein injection. For multiple-dose studies in the UK, mice received siRNA once weekly for three consecutive weeks at 400 nmol/kg via the same route of injection. Animals were euthanized seven days after the final dose, and tissues including liver, heart, skeletal muscle (*tibialis anterior*), kidneys and brain were harvested and stored at -80C until further processing. For ASO-injected mice, only liver, heart and skeletal muscle (quadriceps) were collected.

### RNA isolation

In Sweden, all tissues except liver were homogenized on TissueLyser II (QIAGEN) for 10 min at 30 Hz/s with a 5 mm stainless steel bead (69989, QIAGEN) in 1 mL of TRI Reagent® (T9424, Sigma-Aldrich). Liver samples were lysed in 5 mL of TRI Reagent® in gentleMACS™ M tubes (130-093-236, Miltenyi Biotec) on gentleMACS™ Dissociator (Miltenyi Biotec) using program RNA_02_01. Depending on the tissue weight, the homogenates were diluted further in TRI Reagent® to proceed with approximately 30 mg of homogenized tissue for RNA extraction according to the manufacturer’s instructions for TRI Reagent®. RNA concentrations were measured on a NanoDrop spectrophotometer.

In the UK, total RNA was extracted from 20–30 mg of tissue using the Maxwell® RSC simplyRNA Tissue Kit (AS1340, Promega) and the Maxwell® RSC Instrument according to the manufacturer’s instructions. RNA concentration was determined using a NanoDrop 2000 Spectrophotometer (Thermo Fisher Scientific).

### RT-qPCR

RNA (1-2 µg, depending on tissue) was reverse-transcribed into cDNA with High Capacity cDNA Reverse Transcription Kit (43-688-13, Applied Biosystems™) according to the manufacturer’s instructions.

#### Sod1/Gapdh analysis

Quantitative PCR was performed on a CFX Opus 96 Real-Time PCR System (Bio-Rad) with TaqMan™ Fast Advanced Master Mix (4444557, Applied Biosystems™) and TaqMan® Gene Expression Assays for *Sod1* (Mm01344233_g1) and *Gapdh* (Mm99999915_g1). Each reaction contained 10 ng cDNA. Following the ΔΔCt method, *Sod1* expression was analyzed relative to *Gapdh*, and normalized to the vehicle-treated control group.

#### Malat1/Rplp0 analysis

Quantitative PCR was performed using the same instrument and master mix with a TaqMan® Gene Expression Assay for *Malat1* (Mm01227912_s1) and custom primers (forward: GAGGAATCAGATGAGGATATGGGA; reverse: AAGCAGGCTGACTTGGTTGC) and probe (TCGGTCTCTTCGACTAATCCCGCCAA) for *Rplp0* (36B4). Each reaction contained 12 ng cDNA. Relative expression levels were calculated using the ΔΔCt method and normalized to the vehicle-treated control group.

### Statistical analysis

For groupwise comparisons, ordinary one-way ANOVA was applied, followed by Dunnett’s multiple comparisons test to evaluate differences between groups. Statistical tests were performed in GraphPad Prism 10, and data are presented as mean ± SD.

## RESULTS AND DISCUSSION

To enable flexible and scalable generation of oligonucleotide-ligand conjugates, we employed a modular platform based on orthogonal click-chemistry reactions. Homo-bifunctional conjugates were generated using strain-promoted azide-alkyne cycloaddition (SPAAC), while hetero-bifunctional conjugates were produced by combining SPAAC with inverse electron demand Diels-Alder (IEDDA) reactions (**Supplementary Figure 1**). Using this strategy, we generated 28 distinct siRNA conjugates that varied in ligand type, valency, and linker architecture (**Supplementary Table 2**). All ligands were attached to the sense strand of a previously validated siRNA targeting mouse Sod1 mRNA (18) to allow for direct comparison of tissue-specific silencing across constructs.

### Bis-fatty-acid conjugation promotes extrahepatic siRNA activity

To investigate how ligand identity and valency influence systemic siRNA delivery across different lipid classes, we generated (5′) mono- and (3′,5′) bis-conjugates of one sterol, lithocholic-acid (LCA); one phospholipid, dioleoylphosphatidylethanolamine (DOPE); and four fatty-acids (C16, C16OOH, C20OOH and C22) (**Figure 1A** and **1B**). Given that ligand hydrophobicity has been shown to contribute to conjugate-mediated oligonucleotide biodistribution (7, 8), we first assessed the relative hydrophobicity of each single-stranded siRNA conjugate using HPLC retention time as a proxy (**Figure 1C**). Comparative retention times were consistent with the expected physicochemical properties of the attached ligands. LCA conjugates were relatively hydrophilic, in line with previous reports (7, 8). As expected, fatty-acid conjugates exhibited increasing hydrophobicity with greater aliphatic chain length and ligand valency, while the presence of terminal carboxyl groups substantially reduced hydrophobicity (e.g., C16 vs. C16OOH). Conjugation of DOPE, which bears two unsaturated C18 acyl chains, resulted in comparatively long HPLC retention times, ranking both mono- and bis-DOPE among the most hydrophobic conjugates tested here (**Figure 1C** and **Supplementary Table 1**). Moreover, we confirmed that all siRNA conjugates remained functional by validating target mRNA knockdown in vitro (**Supplementary Figure 2**).

**Figure 1:**
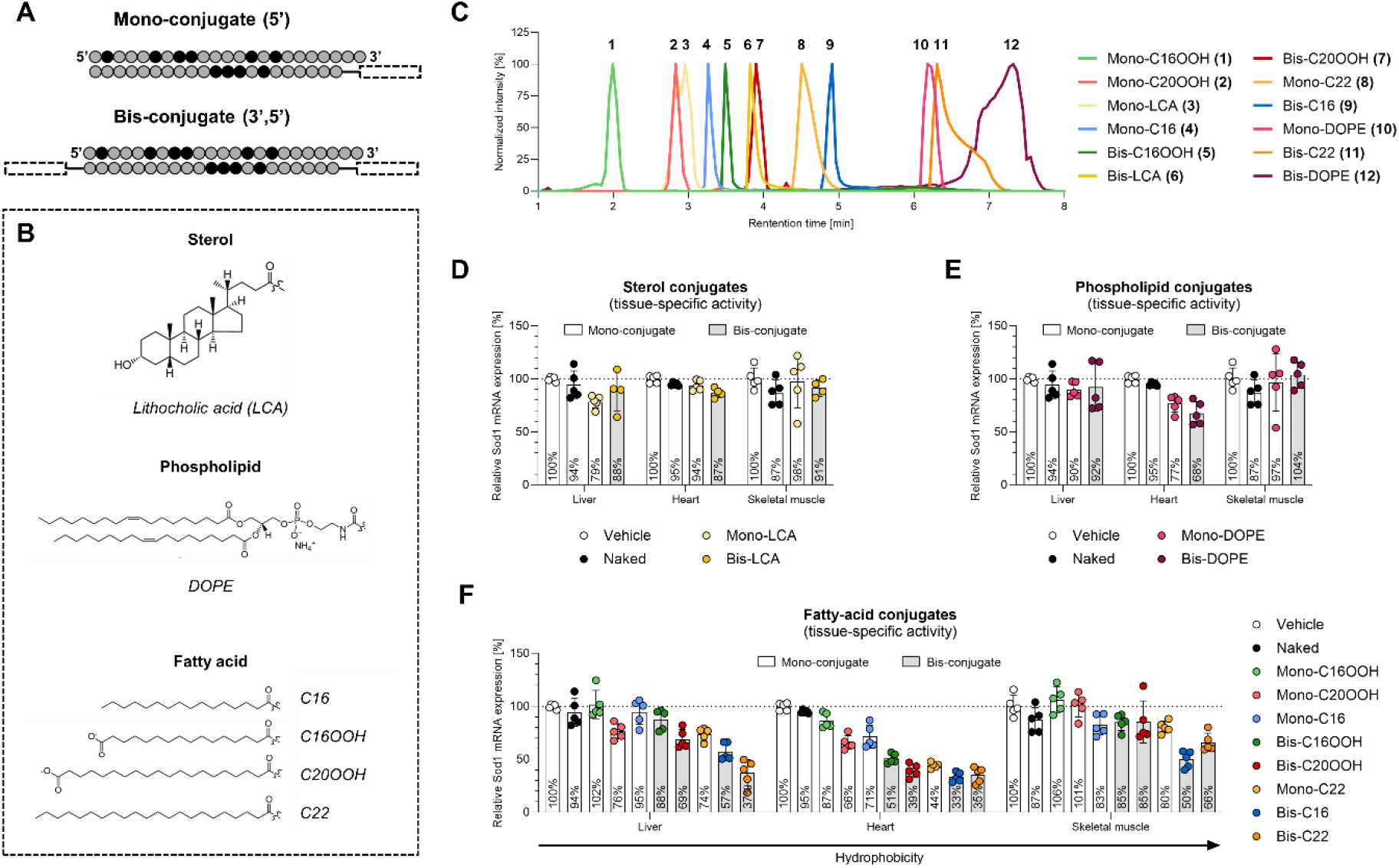
Characterization and in vivo evaluation of mono- and bis-lipid siRNA conjugates. **A)** Schematic representation of the conjugation architecture and the Sod1 siRNA, which contains 2′-deoxy-2′-fluoro (2′-F, black) and 2′-O-methyl (2′-OMe, grey) ribosugar modifications. Ligands were attached to the sense strand resulting in (5′) mono- or (3′,5′) bis-conjugates. **B)** Chemical structures of lipid ligands. **C)** Reversed-phase HPLC chromatograms showing retention times of lipid ligands conjugated to the siRNA sense strand. **D-F)** Target mRNA levels measured by RT-qPCR in the indicated tissues 7 days after tail-vein injection of sterol-, phospholipid-, or fatty-acid-siRNA conjugates at a dose of 600 nmol/kg (n=5). Fatty-acid conjugates are arranged by hydrophobicity. Statistical analysis was performed using ordinary one-way ANOVA followed by Dunnett’s multiple comparisons, comparing each treatment group with the vehicle control for each tissue separately. P values are indicated as * P ≤ 0.05, ** P ≤ 0.01, *** P ≤ 0.001, **** P ≤ 0.0001.

Next, we assessed conjugate-mediated siRNA delivery in vivo by measuring siRNA activity, based on target mRNA knockdown, across several tissues. All lipid conjugates demonstrated varying degrees of activity in liver, heart or skeletal muscle tissue, but not in kidney or brain (**Supplementary Figure 3**). Mono-LCA showed negligible levels of target knockdown, which is consistent with previous reports (7, 8); and increasing ligand valency (bis-LCA) did not result in any improvement (**Figure 1D**). In contrast, conjugation of the phospholipid DOPE, which was selected for its ability to promote membrane interactions (19), resulted in modest yet selective siRNA activity in the heart at the tested dose (23 ± 8% knockdown). Compared to its mono counterpart, bis-DOPE showed minor improvements in heart-selective knockdown efficiency (32 ± 11% knockdown) (**Figure 1E**). The benefit of bis-conjugation was more pronounced for fatty-acid ligands. Conjugation of C16, C20 and C22 - regardless of terminal modification - greatly enhanced siRNA activity in the liver, heart and skeletal muscle (**Figure 1F**). Notably, all bis-fatty-acid conjugates consistently outperformed their mono counterparts, which underscores the overall advantage of increased molecular size and/or hydrophobicity. This trend aligns with previous findings linking hydrophobicity to serum protein binding, which in turn influences clearance and biodistribution of oligonucleotide-lipid conjugates (6, 7, 11). Additionally, these findings highlight the importance of ligand identity, with fatty-acid conjugates exhibiting potent siRNA delivery, particularly to extrahepatic tissues, which is in agreement with earlier observations in the field (7, 8, 20).

In addition to valency, chain length and terminal modification of ligands also influenced the activity of fatty-acid-siRNA conjugates. Interestingly, hydrophobicity appeared to drive productive uptake in the heart and, to a similar extent, in the liver, as evidenced by increased knockdown for fatty-acid conjugates with longer HPLC retention times (**Figure 1C** and **1F**). In fact, bis-C16OOH, bis-C20OOH and mono-C22 demonstrated relatively selective cardiac activity, with favorable tissue-to-liver ratios (≤ 0.6) (**Supplementary Figure 4**), which could be of interest for cardiac indications. In skeletal muscle tissue, only bis-C16 and bis-C22 exhibited substantial activity at this dose (50 ± 8% and 34 ± 9% knockdown, respectively), while all other fatty-acid conjugates performed poorly (**Figure 1F**). For bis-C22, however, the improvements in extrahepatic knockdown were accompanied by increased hepatic activity, resulting in unfavorable tissue-to-liver ratios (0.95 in heart and 1.78 in skeletal muscle) (**Supplementary Figure 4**). Notably, previous studies have reported that more hydrophobic lipid-siRNA conjugates tend to associate with lipoproteins, thereby promoting hepatic uptake (6, 11, 21). Additionally, bis-C22 exhibited the lowest critical aggregation concentration (CAC; 2.12 μM) among these fatty-acid conjugates (**Supplementary Figure 5**), indicating a stronger tendency to form aggregates, which may limit the availability of monomeric species for albumin interactions (22). These observations suggest a threshold for hydrophobicity (and aggregation) beyond which extrahepatic activity is compromised. Apart from that, bis-C16 demonstrated the strongest overall extrahepatic performance among all conjugates, achieving not only the highest knockdown in heart (67 ± 5%) but also in skeletal muscle (50 ± 8%). It is likely that albumin binding drives this effect as has been demonstrated for lipid-ASO conjugates (11, 12). Together, these observations indicate that hydrophobicity alone is not the determining factor that dictates productive delivery of fatty-acid-conjugated siRNAs to specific tissues.

Building on these observations, we explored whether modifications to the terminus of fatty-acid conjugates could further modulate tissue-specific siRNA activity. We hypothesized that introducing a terminal carboxyl group might tune delivery properties through pH-dependent ionization that could affect endosomal escape, as well as albumin interactions. However, the introduction of terminal carboxyl groups (C16OOH and C20OOH) appeared to reduce the overall activity in relation to unmodified fatty-acid conjugates (**Figure 1F**). This decrease may result from impaired cellular uptake caused by electrostatic repulsion between the negatively charged carboxyl termini and the cell membrane. Consistent with this notion, carboxyl-terminated lipids have been shown to reduce cellular uptake of siRNA conjugates in vitro (14). Despite the reduction in overall activity, comparing tissue-to-liver ratios of bis-C16 and bis-C16OOH revealed that carboxyl-terminated lipids maintained relative heart-specific siRNA activity (0.58) while losing relative skeletal muscle activity (0.87 vs. 0.97 respectively) (**Supplementary Figure 4**). These findings indicate that terminal modifications can fine-tune extrahepatic distribution. Such insights can be useful in guiding therapeutic strategies aimed at enhancing tissue-specific siRNA activity.

Similarly, previous studies have shown that inserting polar phosphatidylcholine (PC) groups between a ligand and the siRNA can modulate the tissue-accumulation profile of lipid-siRNA conjugates (8, 23). Building on this concept, we sought to examine the role of linker chemistry on the performance of a subset of our lipid-siRNA conjugates. To this end, we designed a linker (L4+, **Supplementary Figure 6**) based on lysine derivatives that provides one permanent positive charge and could potentially promote extrahepatic delivery in a similar manner to PC. Compared to the neutral PEG3 linker (L1), incorporation of L4+ had no significant impact on the activity of bis-C16 and bis-C22 conjugates (**Supplementary Figure 7**). For mono-C16 conjugates, switching from the L1 to the L4+ linker resulted in minor improvements in activity across liver, heart, and skeletal muscle, whereas for mono-C22 conjugates, the opposite effect was observed. However, since improvements were minimal and relative target knockdown in extrahepatic tissues remained mostly unchanged, the linker was not further investigated.

### C16 conjugation architecture matters for siRNA activity

To further explore the potential of C16-siRNA conjugates for skeletal muscle and heart delivery, we investigated alternative conjugate configurations, including PEG linkers and multi-lipid arrangements (**Figure 2A** and **2B**). PEGs are commonly used to improve solubility, which can benefit hydrophobic conjugates. However, introducing PEG into mono-C16 conjugates had negligible effects on activity, with PEG2000 even slightly increasing productive uptake in the liver while reducing extrahepatic delivery (**Figure 2E**). Similarly, incorporation of PEG2000 abolished DOPE-mediated siRNA activity in the heart (**Figure 2C** and **2F**). Notably, long PEG linkers (≥ 30 units) have been shown to reduce the in vivo selectivity of lipid-siRNA conjugates for albumin over lipoproteins thereby diminishing the pharmacokinetic advantages conferred by albumin association (14). This behavior was attributed to aggregation, consistent with our observation that PEG2000 incorporation lowered the CAC of both DOPE and mono-C16 conjugates (**Supplementary Figure 5**). Even introduction of PEG4 into the bis-C16 conjugate substantially reduced the CAC (from 8.43 μM to 0.9 μM), which likely explains the overall loss of activity relative to bis-C16.

**Figure 2:**
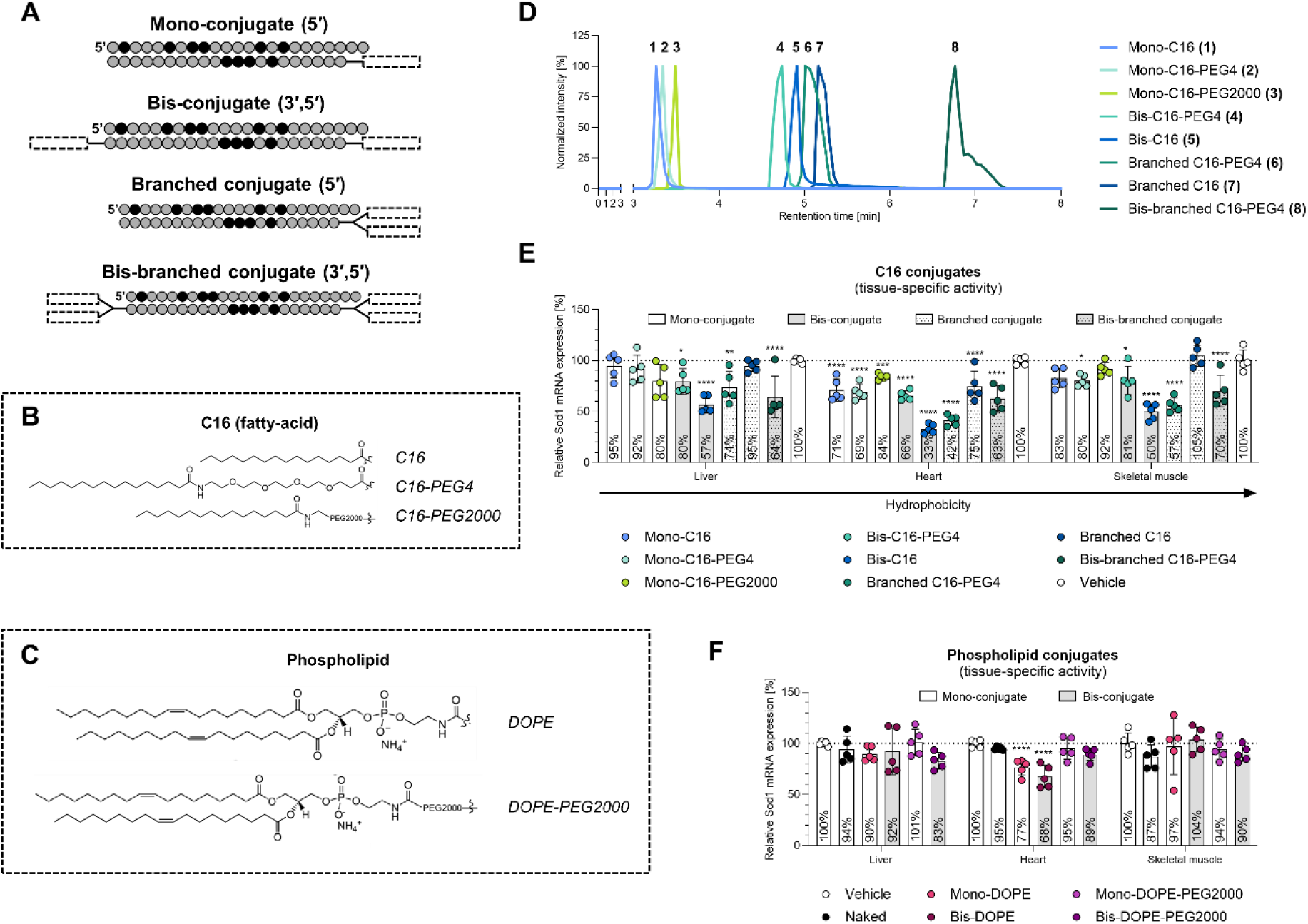
Effect of C16 conjugation architecture on tissue-specific siRNA activity. **A)** Schematic representation of the conjugation architectures and Sod1 siRNA. Ligands were attached to the sense strand, resulting in (5′) mono-, (3′,5′) bis-, (5′) branched, or (3′,5′) bis-branched conjugates. **B, C)** Chemical structures of C16 and DOPE ligands. **D)** Reversed-phase HPLC chromatograms showing retention times of C16 conjugated to the siRNA sense strand. **E, F)** Target mRNA levels measured by RT-qPCR in the indicated tissues 7 days after tail-vein injection at a dose of 600 nmol/kg (n=5). Statistical analysis was performed using ordinary one-way ANOVA followed by Dunnett’s multiple comparisons, comparing each treatment group with the vehicle control for each tissue separately. P values are indicated as * P ≤ 0.05, ** P ≤ 0.01, *** P ≤ 0.001, **** P ≤ 0.0001.

We next evaluated how different C16 arrangements influence siRNA delivery. Attaching two lipids on the same terminus (branched C16) offered no advantage over the mono-C16 conjugate and was inferior to bis-C16. Branched C16-PEG4, on the other hand, performed comparably to bis-C16, suggesting that PEG4 may provide flexibility critical to albumin interactions in this configuration. This aligns with prior findings indicating that having a proximal branchpoint with PEG spacers between the siRNA and lipids is more beneficial for branched C18 conjugates (14). Of note, tissue-to-liver ratios were largely unaffected by configuration architecture, as evidenced by similar ratios for conjugates with comparable hydrophobicity (**Figure 2D** and **Supplementary Figure 4**). Lastly, when we integrated these configurations into a bis-branched C16-PEG4 construct (CAC = 0.44 μM, **Supplementary Figure 5**), its activity shifted toward the liver. This, again, points to a hydrophobic (and aggregation-related) threshold that limits extrahepatic delivery. Overall, these results highlight the importance of lipid arrangements and linker design in balancing ligand flexibility and hydrophobicity/aggregation to optimize extrahepatic siRNA delivery with C16 ligands.

Apart from that, to determine whether extrahepatic activity was influenced by dosing regimen, selected conjugates (bis-C16, branched C16-PEG4, bis-DOPE, and mono-DOPE-PEG2000) were evaluated under a repeat-dose schedule (3 × 400 nmol/kg, once weekly). Knockdown levels were comparable to those observed after a single dose across liver, heart, and skeletal muscle (**Supplementary Figure 8**), indicating that extrahepatic silencing is reproducible and not a consequence of dose rate–dependent delivery mechanisms.

### Peptide ligands provide modest benefits for lipid-siRNA conjugates

Beyond lipids, which rely largely on passive, protein-assisted delivery, we next evaluated peptides for their ability to drive more active siRNA delivery. Peptides can support receptor-mediated uptake and promote endosomal escape. Here, we evaluated a cardiomyocyte-targeting peptide (CTP), which has previously been shown to enhance cardiac accumulation of phages (24) and promote cardiomyocyte-specific uptake of CTP-engineered extracellular vesicles (25). Moreover, to introduce both membrane-interacting and endosomal-escape capabilities, we designed a novel peptide. This peptide consists of ELL, a membrane-interacting amphiphilic sequence inspired by GALA-based motifs (26, 27), and GWWG, an endosomal-escape domain (28), linked via an aliphatic spacer (C6). To determine how these peptides influence siRNA delivery, we generated homo-functional (peptide-only) conjugates as well as hetero-bifunctional conjugates containing both a peptide and a lipid ligand (**Figure 3A** and **3B**).

**Figure 3:**
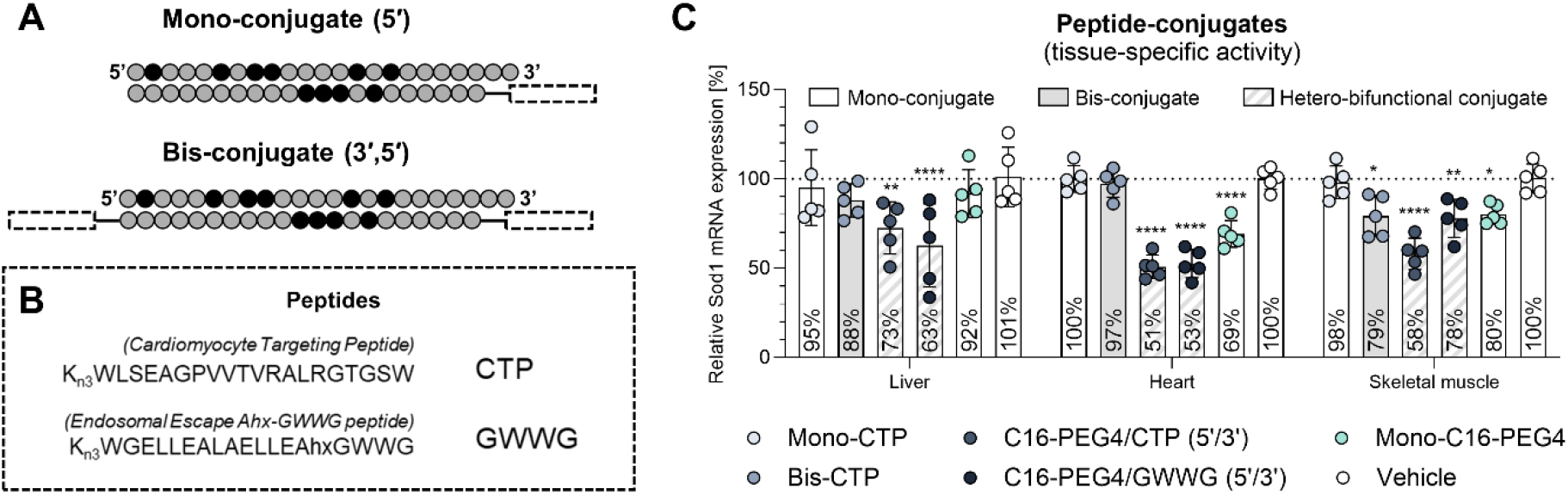
Evaluation of peptide conjugates and peptide-lipid combinations for siRNA delivery. **A)** Schematic representation of the conjugation architecture and the Sod1 siRNA. Ligands were attached to the sense strand resulting in (5′) mono- or (3′,5′) bis-conjugates. **B)** Peptide sequences. **C)** Target mRNA levels measured by RT-qPCR in the indicated tissues 7 days after tail-vein injection of siRNA conjugates at a dose of 600 nmol/kg (n=5). Statistical analysis was performed using ordinary one-way ANOVA followed by Dunnett’s multiple comparisons, comparing each treatment group with the vehicle control for each tissue separately. P values are indicated as * P ≤ 0.05, ** P ≤ 0.01, *** P ≤ 0.001, **** P ≤ 0.0001.

Homo-functional conjugates, particularly mono- and bis-CTP, showed negligible siRNA activity across tested tissues, aside from a modest effect in skeletal muscle for bis-CTP (21 ± 12% knockdown) (**Figure 3C**). While siRNA accumulation was not quantified, the lack of detectable knockdown in the heart suggests that any CTP-mediated uptake into cardiomyocytes, if it occurs, is non-productive. Notably, combining (5′) CTP with C16-PEG4 at the 3′ terminus slightly increased overall potency without altering relative tissue distribution profiles in comparison to C16-PEG4 alone (**Supplementary Figure 4**). Thus, CTP appears unlikely to impart cardiomyocyte-targeting capabilities. In contrast, conjugation of GWWG to C16-PEG4 siRNA altered tissue distribution by increasing activity in cardiac and hepatic tissues but not in skeletal muscle (**Figure 3C** and **Supplementary Figure 4**). Of note, a hetero-bifunctional peptide conjugate (5′ CTP and 3′ GWWG) was successfully synthesized but showed no activity in vitro (**Supplementary Figure 2**). Overall, these findings suggest that combining lipid and peptide ligands can improve delivery efficacy, but their impact on tissue distribution requires careful ligand selection and systematic evaluation.

### Bis-fatty-acid conjugation does not translate to ASOs

Inspired by the promising results of bis-lipid conjugation for siRNA delivery, we next examined whether this approach could translate to gapmer ASOs, another important class of clinically used oligonucleotide therapeutics. To this end, we generated (5′) mono conjugates of C16 and C22, as well as a bis-C16 conjugate, using a Malat1-targeting, fully phosphorothioate-modified LNA–DNA–LNA (4-9-3) gapmer (**Figure 4A**). Following intravenous administration at 1 µmol/kg, liver, heart, and skeletal muscle tissues were collected for analysis seven days later. Unconjugated ASOs produced robust knockdown in liver, with more modest effects in skeletal muscle and heart (**Figure 4B**). Such activity is expected, as fully PS-modified ASOs are known to interact extensively with plasma proteins which facilitates delivery (29, 30). Relative to the parent ASO, mono-conjugation with either C16 or C22 enhanced activity in liver and skeletal muscle, and to a lesser extent in heart, which is (partially) consistent with previous reports (11, 12, 31, 32). In contrast, bis-C16 showed markedly reduced potency across all tissues, effectively abolishing activity (**Figure 4B**). This loss of activity may arise from excessive hydrophobicity leading to aggregation (CAC = 0.18 μM, **Figure 4C**) and consequent alterations in protein-binding profiles (22). Therefore, while bis-fatty-acid conjugation offers significant improvements for siRNA delivery this configuration does not apply to ASOs, most likely due to the physicochemical properties of the oligonucleotide itself.

**Figure 4:**
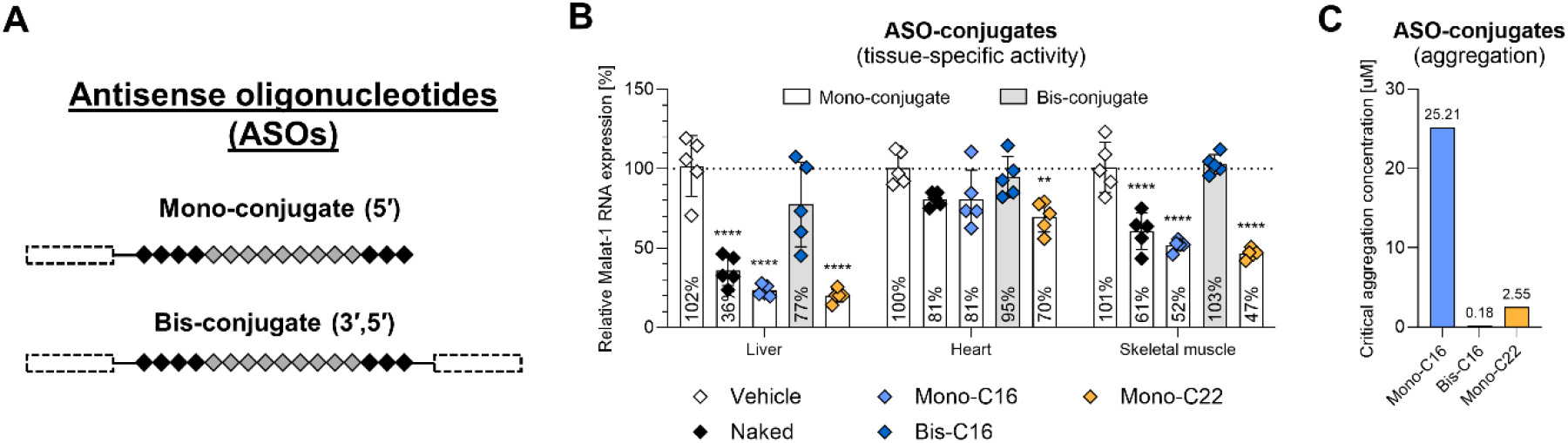
Evaluation of mono- and bis-lipid conjugation for ASO delivery. **A)** Schematic representation of ASO conjugation architecture. **B)** Target mRNA levels measured by RT-qPCR in the indicated tissues 7 days after tail-vein injection of ASO conjugates at a dose of 1 µmol/kg (n=5). Statistical analysis was performed using ordinary one-way ANOVA followed by Dunnett’s multiple comparisons, comparing each treatment group with the vehicle control for each tissue separately. **C)** Critical aggregation concentration (CAC) of ASO conjugates determined using Nile Red assay. Statistical analysis was performed using ordinary one-way ANOVA followed by Dunnett’s multiple comparisons, comparing each treatment group with the vehicle control for each tissue separately. P values are indicated as * P ≤ 0.05, ** P ≤ 0.01, *** P ≤ 0.001, **** P ≤ 0.0001.

## CONCLUSION

Using a modular click-chemistry platform, we mapped how ligand identity, valency, and architecture shape conjugate-mediated siRNA delivery. Fatty-acid ligands, particularly C16-based conjugates, emerged as the most effective designs for productive gene silencing in heart and skeletal muscle. Subtle architectural changes, such as PEG-supported branching, further demonstrated that balancing hydrophobicity, flexibility, and aggregation is essential for optimizing activity. In contrast, peptide ligands provided only modest benefits, and the lack of translatability of bis-fatty-acid designs to ASOs highlights how oligonucleotide chemistry fundamentally constrains conjugate efficacy.

While these findings provide a framework for rational conjugate design, several limitations should be considered. First, our observations are based on intravenous administration; highly hydrophobic constructs may behave differently following subcutaneous dosing, where injection site retention can substantially affect systemic exposure (20). Second, although hydrophobicity (partially) influences biodistribution, factors such as aggregation behavior, membrane-insertion propensity, and other physicochemical properties also appear to shape cellular uptake and productive silencing (20, 22). Third, conclusions were drawn from a single siRNA sequence and target gene; apparent potency may be shaped by biological context, including differences in target expression, turnover, and other cellular factors across tissues (33), emphasizing the importance of validation with additional sequences. Finally, the repeat-dose experiment demonstrated reproducibility of extrahepatic silencing across dosing regimens; however, as tissues were analyzed at a single timepoint, the durability of silencing remains to be established. Longitudinal biodistribution and pharmacodynamic studies will be important for linking delivery properties to duration of action and for defining the optimal dosing regimen for extrahepatic applications. In addition, systematic toxicology and tolerability studies will be essential to ensure the safety of these conjugates.

Together, these findings outline design principles and caveats for expanding siRNA therapeutics beyond hepatic targets.

## ACKNOWLEDGEMENTS

The authors thank Mattias Bood and Anders Dahlén, both from AstraZeneca, for providing the siRNA oligonucleotides used in this study. The graphical abstract was created in BioRender. Roudi, S. (2026) https://BioRender.com/g24ka7a.

## AUTHOR CONTRIBUTIONS

J.A.R. (Investigation, Formal analysis, Writing – original draft, Writing – review & editing, Supervision), E.F. (Investigation, Writing – review & editing), A.M. (Investigation, Writing – review & editing), M.O. (Investigation, Writing – review & editing), T.C. (Investigation, Writing – review & editing), A.H. (Supervision, Writing – review & editing), N.A. (Investigation), S.R. (Investigation, Supervision), C.B. (Investigation), Y.H. (Investigation), O.S. (Investigation), M.W. (Supervision), R.Z. (Supervision), M.H. (Conceptualization, Investigation, Supervision, Writing – review & editing), S.E.A. (Conceptualization, Supervision, Writing – review & editing). All authors approved the final manuscript.

## SUPPLEMENTARY DATA

Supplementary Data are available at NAR online.

## DATA AVAILABILITY

The data underlying this article will be shared on reasonable request to the corresponding author.

## REFERENCES

1. Khvorova, A. and Watts, J.K. (2017) The chemical evolution of oligonucleotide therapies of clinical utility. Nature Biotechnology 2017 35:3, 35, 238–248.

2. Tang, Q. and Khvorova, A. (2024) RNAi-based drug design: considerations and future directions. Nature Reviews Drug Discovery 2024 23:5, 23, 341–364.

3. Nair, J.K., Willoughby, J.L.S., Chan, A., Charisse, K., Alam, M.R., Wang, Q., Hoekstra, M., Kandasamy, P., Kelin, A. V., Milstein, S., et al. (2014) Multivalent N-Acetylgalactosamine-Conjugated siRNA Localizes in Hepatocytes and Elicits Robust RNAi-Mediated Gene Silencing. J. Am. Chem. Soc., 136, 16958–16961.

4. Anand, P., Zhang, Y., Patil, S. and Kaur, K. (2025) Metabolic Stability and Targeted Delivery of Oligonucleotides: Advancing RNA Therapeutics Beyond The Liver. J. Med. Chem., 68, 6870–6896.

5. Soutschek, J., Akinc, A., Bramlage, B., Charisse, K., Constien, R., Donoghue, M., Elbashir, S., Gelck, A., Hadwiger, P., Harborth, J., et al. (2004) Therapeutic silencing of an endogenous gene by systemic administration of modified siRNAs. Nature 2004 432:7014, 432, 173–178.

6. Wolfrum, C., Shi, S., Jayaprakash, K.N., Jayaraman, M., Wang, G., Pandey, R.K., Rajeev, K.G., Nakayama, T., Charrise, K., Ndungo, E.M., et al. (2007) Mechanisms and optimization of in vivo delivery of lipophilic siRNAs. Nature Biotechnology 2007 25:10, 25, 1149–1157.

7. Osborn, M.F., Coles, A.H., Biscans, A., Haraszti, R.A., Roux, L., Davis, S., Ly, S., Echeverria, D., Hassler, M.R., Godinho, B.M.D.C., et al. (2019) Hydrophobicity drives the systemic distribution of lipid-conjugated siRNAs via lipid transport pathways. Nucleic Acids Res., 47, 1070–1081.

8. Biscans, A., Coles, A., Haraszti, R., Echeverria, Di., Hassler, M., Osborn, M. and Khvorova, A. (2019) Diverse lipid conjugates for functional extra-hepatic siRNA delivery in vivo. Nucleic Acids Res., 47, 1082–1096.

9. Biscans, A., Caiazzi, J., McHugh, N., Hariharan, V., Muhuri, M. and Khvorova, A. (2021) Docosanoic acid conjugation to siRNA enables functional and safe delivery to skeletal and cardiac muscles. Molecular Therapy, 29, 1382–1394.

10. Fakih, H.H., Lochmann, C., Gagnon, R., Summers, A., Caiazzi, J., Buchwald, J.E., Tang, Q., wit Maru, B., Hildebrand, S.R., Zain Abideen, M.U., et al. (2025) Potent and durable gene modulation in heart and muscle with chemically defined lipophilic siRNAs. Nucleic Acids Res., 53, 13–14.

11. Prakash, T.P., Mullick, A.E., Lee, R.G., Yu, J., Yeh, S.T., Low, A., Chappell, A.E., Østergaard, M.E., Murray, S., Gaus, H.J., et al. (2019) Fatty acid conjugation enhances potency of antisense oligonucleotides in muscle. Nucleic Acids Res., 47, 6029–6044.

12. Chappell, A.E., Gaus, H.J., Berdeja, A., Gupta, R., Jo, M., Prakash, T.P., Oestergaard, M., Swayze, E.E. and Seth, P.P. (2020) Mechanisms of palmitic acid-conjugated antisense oligonucleotide distribution in mice. Nucleic Acids Res., 48, 4382–4395.

13. Sarett, S.M., Werfel, T.A., Lee, L., Jackson, M.A., Kilchrist, K. V., Brantley-Sieders, D. and Duvall, C.L. (2017) Lipophilic siRNA targets albumin in situ and promotes bioavailability, tumor penetration, and carrier-free gene silencing. Proc. Natl. Acad. Sci. U. S. A., 114, E6490–E6497.

14. Hoogenboezem, E.N., Patel, S.S., Lo, J.H., Cavnar, A.B., Babb, L.M., Francini, N., Gbur, E.F., Patil, P., Colazo, J.M., Michell, D.L., et al. (2024) Structural optimization of siRNA conjugates for albumin binding achieves effective MCL1-directed cancer therapy. Nature Communications 2024 15:1, 15, 1–20.

15. Sorets, A.G., Schwensen, K.R., Francini, N., Kjar, A., Lyons, S., Park, J.C., Palmer, D., Abdulrahman, A.M., Cowell, R.P., Katdare, K.A., et al. (2025) Intravenous lipid-siRNA conjugate mediates gene silencing at the blood-brain barrier and blood-CSF barrier. Journal of Controlled Release, 387, 114226.

16. Fakih, H.H., Tang, Q., Summers, A., Shin, M., Buchwald, J.E., Gagnon, R., Hariharan, V.N., Echeverria, D., Cooper, D.A., Watts, J.K., et al. (2023) Dendritic amphiphilic siRNA: Selective albumin binding, in vivo efficacy, and low toxicity. Mol. Ther. Nucleic Acids, 34, 102080.

17. Fakih, H.H., Tang, Q., Summers, A., Gross, K.Y., Rachid, M.O., Okamura, K., Martinez, N., Sleiman, H.F., Harris, J.E. and Khvorova, A. (2025) Albumin-binding dendritic siRNA improves delivery and efficacy to solid tumors in a melanoma model. Mol. Ther. Nucleic Acids, 36, 102579.

18. Brown, K.M., Nair, J.K., Janas, M.M., Anglero-Rodriguez, Y.I., Dang, L.T.H., Peng, H., Theile, C.S., Castellanos-Rizaldos, E., Brown, C., Foster, D., et al. (2022) Expanding RNAi therapeutics to extrahepatic tissues with lipophilic conjugates. Nature Biotechnology 2022 40:10, 40, 1500–1508.

19. Zheng, W., Schürz, M., Wiklander, R.J., Gustafsson, O., Gupta, D., Slovak, R., Traista, A., Coluzzi, A., Roudi, S., Barone, A., et al. (2023) Surface display of functional moieties on extracellular vesicles using lipid anchors. Journal of Controlled Release, 357, 630–640.

20. Biscans, A., Coles, A., Echeverria, D. and Khvorova, A. (2019) The valency of fatty acid conjugates impacts siRNA pharmacokinetics, distribution, and efficacy in vivo. Journal of Controlled Release, 302, 116–125.

21. Hvam, M.L., Cai, Y., Dagnæs-Hansen, F., Nielsen, J.S., Wengel, J., Kjems, J. and Howard, K.A. (2017) Fatty Acid-Modified Gapmer Antisense Oligonucleotide and Serum Albumin Constructs for Pharmacokinetic Modulation. Molecular Therapy, 25, 1710–1717.

22. Kusznir, E.A., Hau, J.C., Portmann, M., Reinhart, A.G., Falivene, F., Bastien, J., Worm, J., Ross, A., Lauer, M., Ringler, P., et al. (2023) Propensities of Fatty Acid-Modified ASOs: Self-Assembly vs Albumin Binding. Bioconjug. Chem., 34, 866–879.

23. Hariharan, V.N., Nakamura, T., Shin, M., Tang, Q., Sontakke, V., Caiazzi, J., Hildebrand, S., Khvorova, A. and Yamada, K. (2024) Phosphatidylcholine head group chemistry alters the extrahepatic accumulation of lipid-conjugated siRNA. Mol. Ther. Nucleic Acids, 35, 102230.

24. McGuire, M.J., Samli, K.N., Johnston, S.A. and Brown, K.C. (2004) In vitro Selection of a Peptide with High Selectivity for Cardiomyocytes In vivo. J. Mol. Biol., 342, 171–182.

25. Mentkowski, K.I. and Lang, J.K. (2019) Exosomes Engineered to Express a Cardiomyocyte Binding Peptide Demonstrate Improved Cardiac Retention in Vivo. Scientific Reports 2019 9:1, 9, 10041-.

26. Neundorf, I., Rennert, R., Hoyer, J., Schramm, F., Löbner, K., Kitanovic, I. and Wölfl, S. (2009) Fusion of a Short HA2-Derived Peptide Sequence to Cell-Penetrating Peptides Improves Cytosolic Uptake, but Enhances Cytotoxic Activity. Pharmaceuticals 2009, Vol. 2, Pages 49- 65, 2, 49–65.

27. Honcharenko, D., Rocha, C.S.J., Lundin, K.E., Maity, J., Milton, S., Tedebark, U., Murtola, M., Honcharenko, M., Slaitas, A., Smith, C.I.E., et al. (2022) 2′-O-(N-(Aminoethyl)carbamoyl)methyl Modification Allows for Lower Phosphorothioate Content in Splice-Switching Oligonucleotides with Retained Activity. https://home.liebertpub.com/nat, 32, 221–233.

28. Lönn, P., Kacsinta, A.D., Cui, X.S., Hamil, A.S., Kaulich, M., Gogoi, K. and Dowdy, S.F. (2016) Enhancing Endosomal Escape for Intracellular Delivery of Macromolecular Biologic Therapeutics. Scientific Reports 2016 6:1, 6, 32301-.

29. Gaus, H.J., Gupta, R., Chappell, A.E., Østergaard, M.E., Swayze, E.E. and Seth, P.P. (2019) Characterization of the interactions of chemically-modified therapeutic nucleic acids with plasma proteins using a fluorescence polarization assay. Nucleic Acids Res., 47, 1110–1122.

30. Crooke, S.T., Wang, S., Vickers, T.A., Shen, W. and Liang, X.H. (2017) Cellular uptake and trafficking of antisense oligonucleotides. Nature Biotechnology 2017 35:3, 35, 230–237.

31. Østergaard, M.E., Jackson, M., Low, A., E Chappell, A., G Lee, R., Peralta, R.Q., Yu, J., Kinberger, G.A., Dan, A., Carty, R., et al. (2019) Conjugation of hydrophobic moieties enhances potency of antisense oligonucleotides in the muscle of rodents and non-human primates. Nucleic Acids Res., 47, 6045–6058.

32. Ait Benichou, S., Jauvin, D., De Serres-Bérard, T., Bennett, F., Rigo, F., Gourdon, G., Boutjdir, M., Chahine, M. and Puymirat, J. (2022) Enhanced Delivery of Ligand-Conjugated Antisense Oligonucleotides (C16-HA-ASO) Targeting Dystrophia Myotonica Protein Kinase Transcripts for the Treatment of Myotonic Dystrophy Type 1. https://home.liebertpub.com/hum, 33, 810–820.

33. Davis, S.M., Hildebrand, S., Macmillan, H.J., Monopoli, K.R., Buchwald, J., Sousa, J., Cooper, D., Ly, S., Echeverria, D., Mchugh, N., et al. (2025) Systematic analysis of siRNA and mRNA features impacting fully chemically modified siRNA efficacy. Nucleic Acids Res., 53, 53.

