## Supplementary Material for "Click Chemistry-Based Strategy for Modular Ligand Attachment to siRNAs: Toward Extrahepatic RNAi"

#### TABLE OF CONTENTS

##### Supplementary Figures

Supplementary Figure 1: Schematic representation of the conjugation strategy.

Supplementary Figure 2: In vitro assessment of siRNA conjugates.

Supplementary Figure 3: Target knockdown of all 28 siRNA-conjugates across indicated tissues.

Supplementary Figure 4: Tissue-to-liver ratios of all 28 siRNA conjugates.

Supplementary Figure 5: Critical aggregation concentrations of siRNA conjugates.

Supplementary Figure 6: Overview of linkers.

Supplementary Figure 7: The impact of linker chemistry on a subset of fatty-acid conjugates.

Supplementary Figure 8: The effect of dosing regimen on tissue-specific siRNA activity.

##### Supplementary Tables

Supplementary Table 1: Sod1 siRNA sequence and chemistry.

Supplementary Table 2: Overview of all siRNA conjugates used in this study.

Supplementary Table 3: Overview of all ASO conjugates used in this study.

##### Synthetic procedures

##### HPLC chromatograms and MS data

#### SUPPLEMENTARY FIGURES

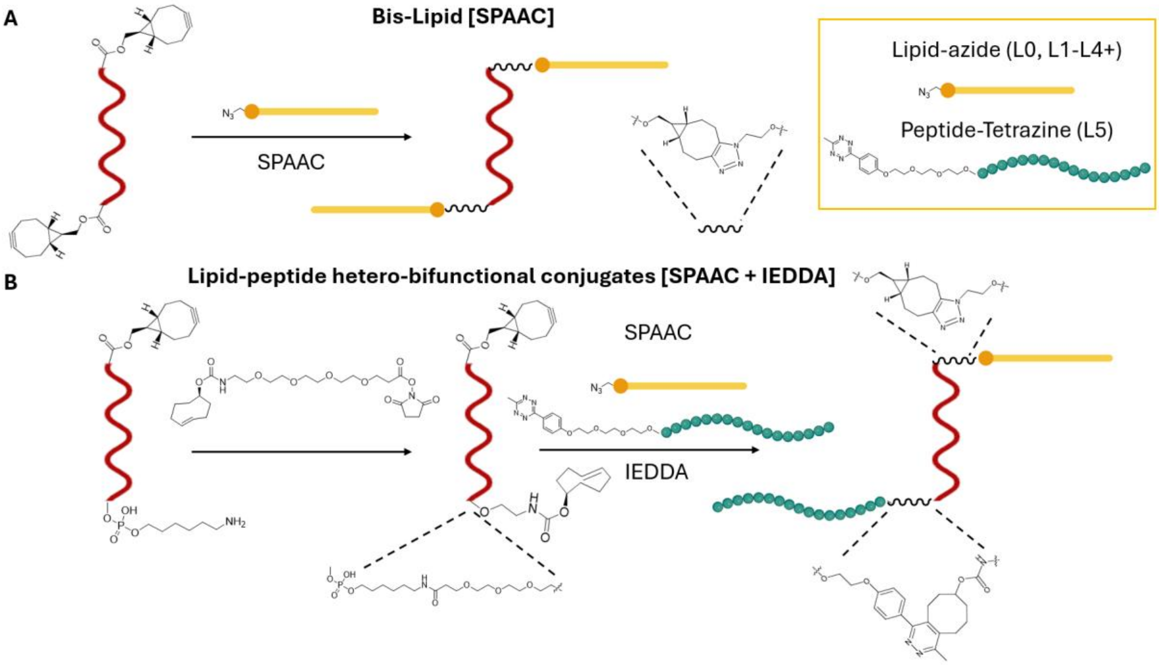


**Supplementary Figure 1: Schematic representation of the conjugation strategy. A) 3′,5′** Double Strain Promoted Azide Alkyne Cycloaddition (SPAAC) used to obtain 3′-5′ bis-lipid conjugates. **B)** Combination of SPAAC and inverse-electron-demand Diels-Alder (IEDDA) conjugation used to obtain 3′-peptide-5′-lipid hetero-bifunctional conjugates.

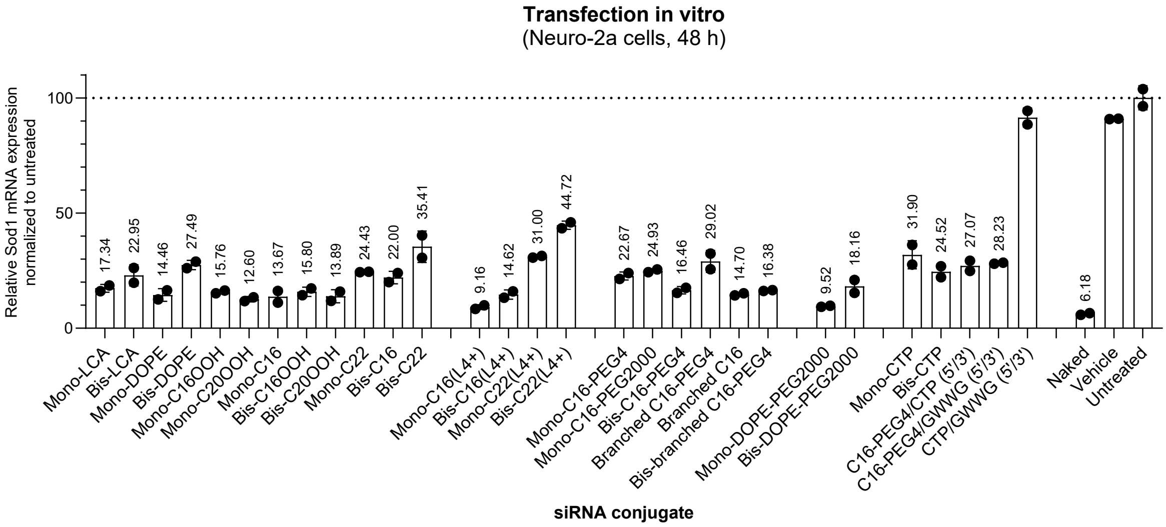


**Supplementary Figure 2: In vitro assessment of siRNA conjugates.** Neuro-2a cells were transfected with 1 pmol/well of the indicated siRNA conjugates using RNAiMAX and analyzed 48 h later. Sod1 mRNA levels were quantified by TaqMan qPCR and normalized to Gapdh, relative to the untreated control.

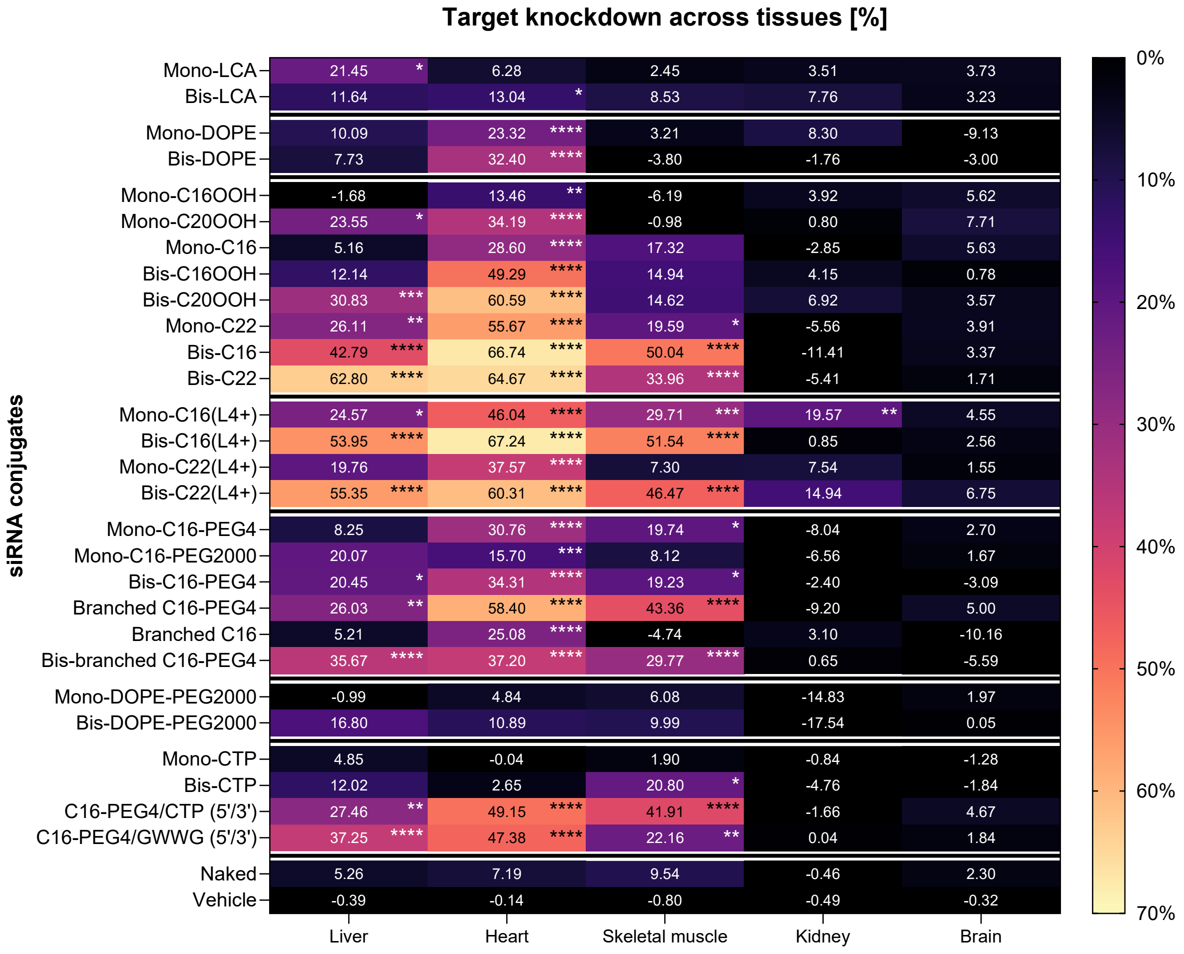


**Supplementary Figure 3: Target knockdown of all 28 siRNA-conjugates across indicated tissues.** Percent target knockdown evaluated by RT-qPCR in the indicated tissues 7 days after tail-vein injection of siRNA conjugates at a dose of 600 nmol/kg (n=5). Statistical analysis was performed using ordinary one-way ANOVA followed by Dunnett's multiple comparisons, comparing each treatment group with the vehicle control for each tissue separately. * P ≤ 0.05, ** P ≤ 0.01, *** P ≤ 0.001, **** P ≤ 0.0001.

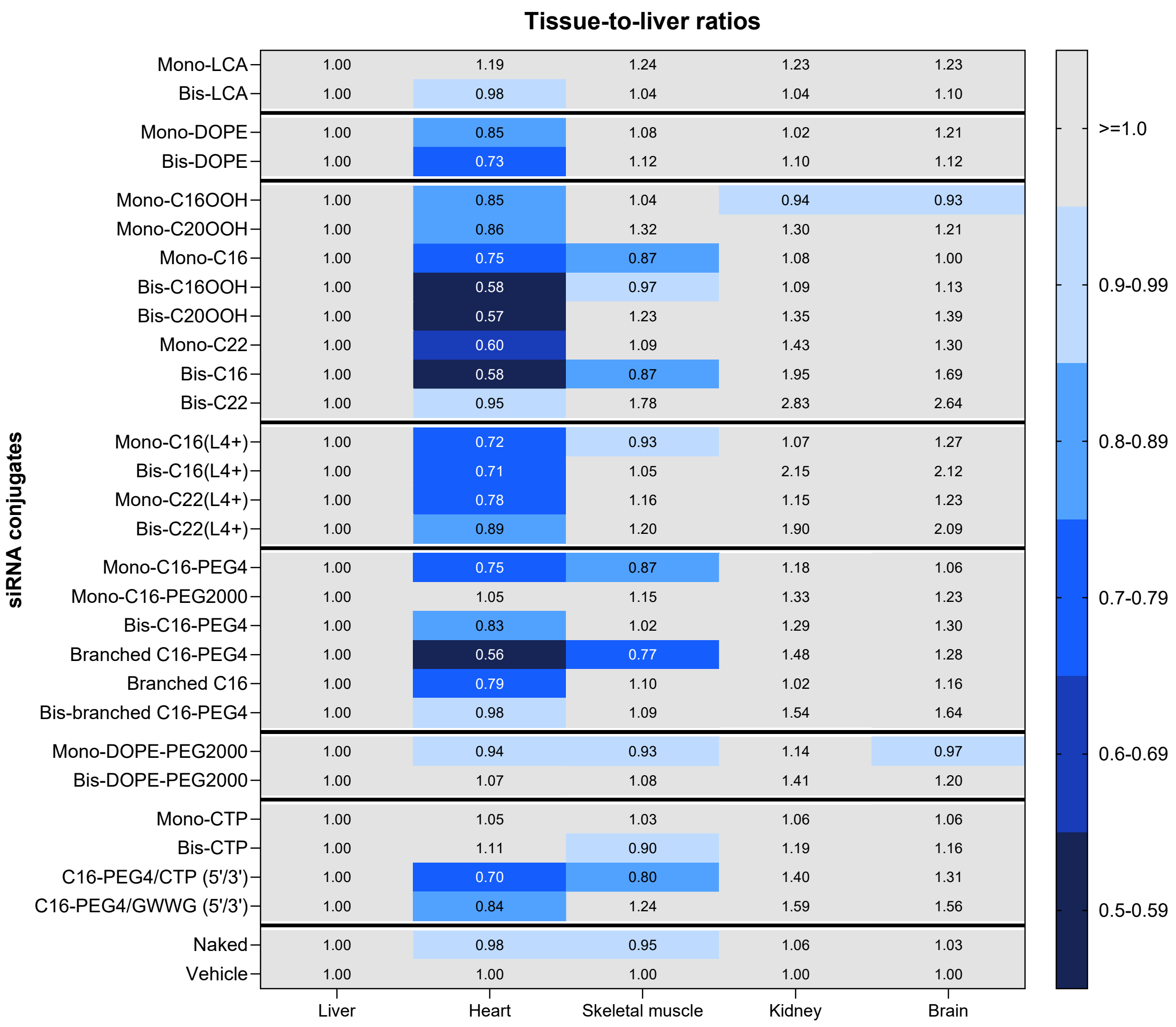


**Supplementary Figure 4: Tissue-to-liver ratios of all 28 siRNA conjugates.** Tissue-to-liver ratios were calculated from RT-qPCR data (shown in Figure 1D-F, Supplementary Figure 6, Figure 2E-F, and Figure 3C) using relative Sod1 mRNA expression in liver as the baseline.

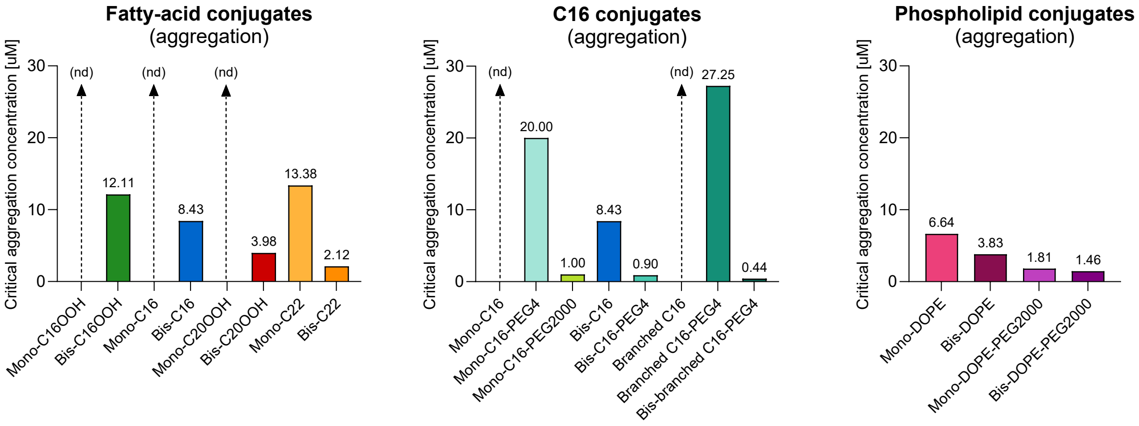


**Supplementary Figure 5: Critical aggregation concentrations of siRNA conjugates.** The critical aggregation concentration (CAC) was defined as the intersection point of the two linear regions in the plot of Nile Red fluorescence versus oligonucleotide concentration.

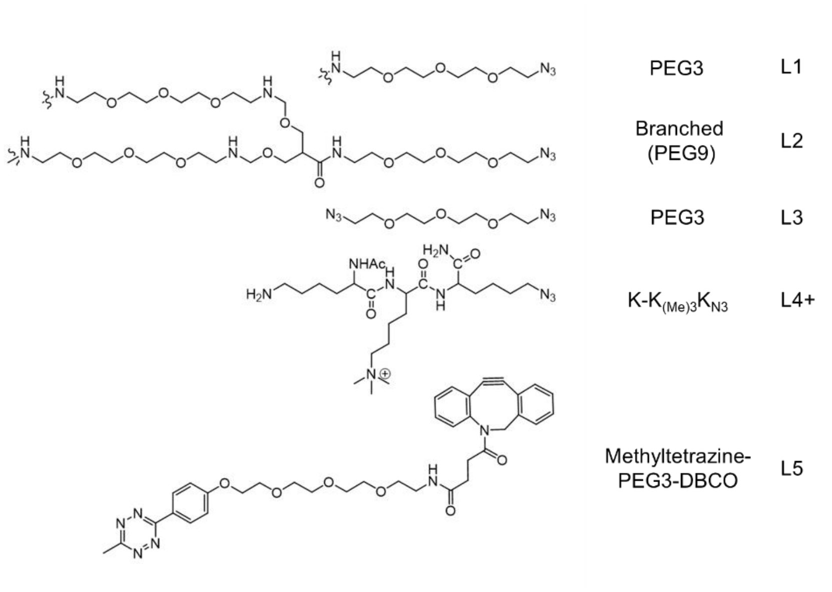


**Supplementary Figure 6: Overview of linkers**. L4+ is a linker based on lysine derivatives. *PEG = polyethylene glycol, DBCO = dibenzocyclooctyne.*

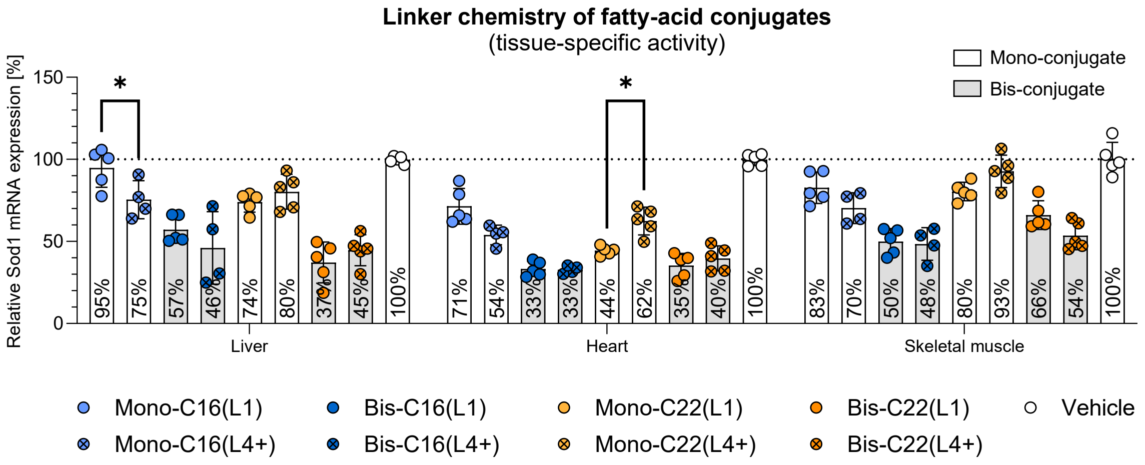


**Supplementary Figure 7: The impact of linker chemistry on a subset of fatty-acid conjugates.** Target mRNA levels measured by RT-qPCR in the indicated tissues 7 days after tail-vein injection of siRNA conjugates at a dose of 600 nmol/kg (n=5). Statistical analysis was performed using ordinary one-way ANOVA followed by Dunnett's multiple comparisons, comparing the respective L1 and L4+ conjugates for each tissue separately. * P ≤ 0.05, ** P ≤ 0.01, *** P ≤ 0.001, **** P ≤ 0.0001.

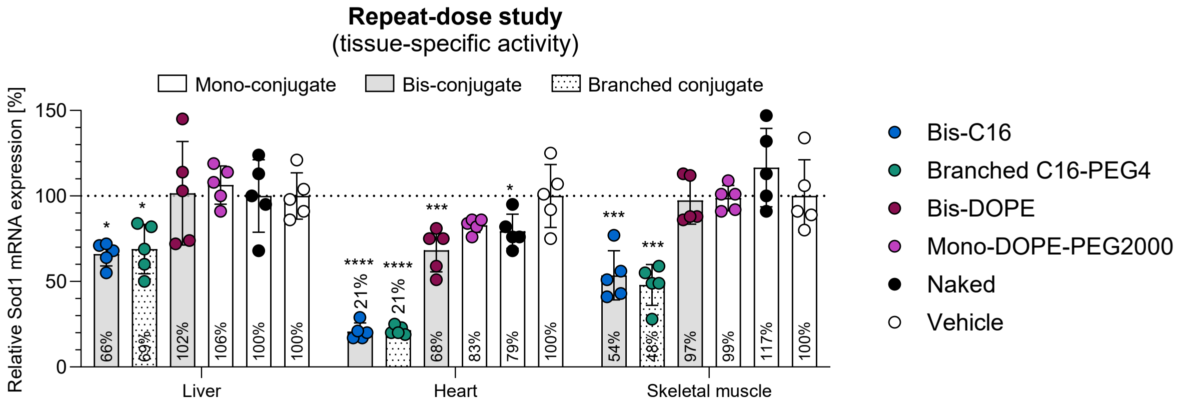


**Supplementary Figure 8: The effect of dosing regimen on tissue-specific siRNA activity.** Target mRNA levels were measured by RT-qPCR in the indicated tissues 7 days after the final tail-vein injection. siRNA conjugates were administered at 400 nmol/kg once weekly for three weeks (n=5). Statistical analysis was performed using ordinary one-way ANOVA followed by Dunnett's multiple comparisons, comparing each treatment group with the vehicle control for each tissue separately. * P ≤ 0.05, ** P ≤ 0.01, *** P ≤ 0.001, **** P ≤ 0.0001.

**SUPPLEMENTARY TABLES**

| Strand | Sequence (5′ to 3′) | Retention time [min] | MS [M] [MaxEnt] |
|---|---|---|---|
| Antisense | u•U•uagAgUGaggaUuAaaaug•a•g | 0.2 | 7772 |
| Sense | BCN-c•a•uuuuAaUCCucacucuaaa | 0.3 | 7400 |

**Supplementary Table 1: Sod1 siRNA sequence and chemistry.** Upper-case and lower-case letters indicate 2′-deoxy-2′-fluoro (2′-F) and 2′-O-methyl (2′-OMe) ribosugar modifications, respectively; • indicate phosphorothioate linkage. The sense strand is modified with a 5′-bicyclo[6.1.0]non-4-yne group (BCN).

| Simplified name | Detailed name | Single-stranded siRNA conjugates | | Double-stranded siRNA conjugates | |
|---|---|---|---|---|---|
| | | Retention time [min] | MS [M] [MaxEnt] | Formula weight [g/mol] | Injected dose [mg/kg] |
| Mono-LCA | 5′-LCA-L1 | 3 | 7976 | 16716 | 10.03 |
| Bis-LCA | 3′,5′-bis-(LCA-L1) | 3.8 | 9152 | 17914 | 10.75 |
| Mono-DOPE | 5′-DOPE-L3 | 6.2 | 8632 | 16316 | 9.79 |
| Bis-DOPE | 3′,5′-bis-(DOPE)-L3 | 7.3 | 10464 | 16917 | 10.15 |
| Mono-C16OOH | 5′-C16OOH-L1 | 2 | 7888 | 16680 | 10.01 |
| Mono-C20OOH | 5′-C20OOH-L1 | 2.8 | 7944 | 16625 | 9.97 |
| Mono-C16 | 5′-C16-L1 | 3.3 | 7857 | 16576 | 9.95 |
| Bis-C16OOH | 3′,5′-bis-(C16OOH-L1) | 3.5 | 8968 | 17844 | 10.71 |
| Bis-C20OOH | 3′,5′-bis-(C20OOH-L1) | 3.9 | 9088 | 17732 | 10.64 |
| Mono-C22 | 5′-C22-L1 | 4.5 | 7940 | 16681 | 10.01 |
| Bis-C16 | 3′,5′-bis-(C16-L1) | 4.9 | 8914 | 17653 | 10.59 |
| Bis-C22 | 3′,5′-bis-(C22-L1) | 6.3 | 9080 | 17846 | 10.71 |
| Mono-C16(L4+) | 5′-C16-L4+ | 3.4 | 7980 | 16720 | 10.03 |
| Bis-C16(L4+) | 3′,5′-bis-(C16-L4+) | 3.8 | 9165 | 17927 | 10.76 |
| Mono-C22(L4+) | 5′-C22-L4+ | 4.1 | 8064 | 16804 | 10.08 |
| Bis-C22(L4+) | 3′,5′-bis-(C22-L4+) | 5.6 | 9329 | 18091 | 10.85 |
| Mono-C16-PEG4 | 5′-C16-PEG4-L1 | 3.3 | 8104 | 16824 | 10.09 |
| Mono-C16-PEG2000 | 5′-C16-PEG2000-L1 | 3.5 | 10850 | 18612 | 11.17 |
| Bis-C16-PEG4 | 3′,5′-bis-(C16-PEG4-L1) | 4.7 | 9407 | 17386 | 10.43 |
| Branched C16-PEG4 | 5′-di(C16-PEG4)-L2 | 5 | 9150 | 18133 | 10.88 |
| Branched C16 | 5′-di(C16)-L2 | 5.2 | 8647 | 16906 | 10.14 |

| | | | | | |
|---|---|---|---|---|---|
| Bis-branched C16-PEG4 | 3′,5′-bis-[di-(C16-PEG4)]-L2 | 6.8 | 11498 | 18751 | 11.25 |
| Mono-DOPE-PEG2000 | 5′-DOPE-PEG2000 | 6.6 | 10211 | 18932 | 11.36 |
| Bis-DOPE-PEG2000 | 3′,5′-bis-(DOPE-PEG2000) | na | 13622 | 22339 | 13.40 |
| Mono-CTP | 5′-CTP | 1.9 | 9735 | 18452 | 11.07 |
| Bis-CTP | 3′,5′-bis-(CTP) | 2.4 | 12672 | 21389 | 12.83 |
| C16-PEG4/CTP (5′/3′) | 5′-C16-PEG4-L1, 3′-CTP-L5 | 3.8 | 11553 | 20296 | 12.18 |
| C16-PEG4/GWWG (5′/3′) | 5′-C16-PEG4-L1, 3′-GWWG-L5 | 4.5 | 11497 | 20240 | 12.14 |
| CTP/GWWG (5′/3′) | 5′-CTP, 3′-GWWG-L5 | 3.8 | 13129 | 21872 | |

**Supplementary Table 2: Overview of all siRNA conjugates used in this study.** Simplified names are used throughout the main text and figures, whereas detailed names appear in the methods section. Retention times were determined by reverse-phase HPLC and molecular masses by mass spectrometry (MS). Injected dose (mg/kg) correspond to an equimolar dose of 600 nmol/kg.

| Simplified name | Detailed name | Retention time [min] | MS [M] [MaxEnt] |
|---|---|---|---|
| Mono-C16 | 5′-C16 | 3.67 | 6497 |
| Bis-C16 | 3′, 5′-bis-C16 | 5.77 | 7653 |
| Mono-C22 | 5′-C22 | 5.14 | 6582 |
| Naked | Malat1 ASO | 0.37 | 5328 |

**Supplementary Table 3: Overview of all ASO conjugates used in this study.** Simplified names are used throughout the main text and figures, whereas detailed names appear in the methods section. Retention times were determined by reverse-phase HPLC and molecular masses by mass spectrometry (MS).

#### SYNTHETIC PROCEDURES

##### Synthesis of azido modified ligands

*Azido-modified palmitic acid (C16-PEG3-$N_3$, C16-PEG4-PEG3-$N_3$, C16-PEG2000-PEG3-$N_3$)*

Synthesis from palmitoyl NHS ester.

Palmitic acid N-hydroxysuccinimide ester (C16-NHS) or palmitoyl(ethylcarbamoyl-PEG4 acid N-hydroxysuccinimide ester (C16-PEG4-NHS) or palmyitoyl(ethylcarbaomyl-PEG2000 acid N-hydroxysuccinimide ester (C16-PEG2000-NHS) (2 equiv, 3 µmol) was dissolved in 20 µL of DMF and mixed with azido-PEG3-amine (1 equiv, 1.5 µmol, 0.327 mg) if precipitation occurred, gentle heating and sonication were applied. The clear reaction mixture was incubated at 50°C for 16 h. The crude product was used for subsequent reactions without further purification.

*Azido-modified docosanoic and lithocholic acid (C22-PEG3-$N_3$, LCA-PEG3-$N_3$).*

Synthesis via HBTU activation.

Docosanoic acid (DCA) or Lithocholic acid (LCA) (0.85 equiv) was dissolved in THF (0.5 mL). HBTU (1.7 equiv) and pyridine were added. Due to partial precipitation, DMF was added until complete dissolution was achieved. Azido-PEG3-amine (1.1 equiv) was added after 30 min, and the reaction mixture was stirred overnight at 80°C. The reaction mixture was diluted with dichloromethane (DCM) and washed with 0.5 M HCl (3x), the organic phase was dried and evaporated and used without further purification.

*Azido-modified double-branched palmitic acid (di(C16)L2).*

Synthesis from palmitoyl NHS ester

Palmitic acid N-hydroxysuccinimide ester (C16-NHS) (2 equiv, 0.0267 mmol, 9.4 mg) was dissolved in DMF (300 µL) was added to azido-PEG2-bis-PEG3-amine (1 equiv, 0.013 mmol, 10 mg) in DMF (100 µL). The reaction mixture was sonicated and gently heated for 16 h. The crude product was used without further purification.

*Azido-modified double-branched palmitic acid (di(C16-peg4)L2).*

Synthesis from palmitic acid PEG4 NHS ester.

Palmitoyl(ethylcarbamoyl-PEG4 acid N-hydroxysuccinimide ester (C16-PEG4-NHS, 3 equiv, 0.039 mmol, 23 mg) was dissolved in DMSO (115 µL) and added to azido-PEG2-bis-PEG3-amine (1 equiv, 0.013 mmol, 10 mg) in DMSO (50 µL). The reaction was sonicated and gently heated for 16 h. The crude product was used without further purification.

*Negative charge ligands (C16OOH-PEG3-$N_3$ and C20OOH-PEG3-$N_3$)*

16-(Azido-PEG3-ethylcarbamoyl)pentadecanoic tert-butyl ester (C16OOtBu-PEG3-$N_3$) or 20-(azide-PEG3-ethylcarbamoyl)nonadecanoic tert-butyl ester (C20OOtBu-PEG3-$N_3$) was dissolved in 80% acetic acid. After 1 h at RT the tert-butyl protecting group was removed. The resulting azido-functionalized carboxylic acids were used without further purification.

*Azido-modified peptides (CTP, ahxGWWG, L4+)*

Peptides (Ahx-GWWG, positively charged linker (L4+) and cardiomyocyte targeting CTP) were synthesized using Fmoc solid-phase peptide synthesis on Biotage Initiator microwave peptide synthesizer (Biotage) under nitrogen. Rink Amide ChemMatrix resin (213 mg, 100 µmol) was swollen in NMP for 20 min in 70 °C. Fmoc deprotection was performed using 20% piperidine in NMP for 13 min at room temperature. Amido acid couplings were carried out using 5 equiv of Fmoc-protected amino acid, 5 equiv DIC and 5 equiv Oxyma in NMP for 6 min in 75°C. Capping was performed using NMP-lutidine-acetic anhydrate (89:6:5, v\v\v) for 2 min at room temperature. After completion of the synthesis, the resin was washed with NMP and DCM. Cleavage and global deprotection were performed using TFA-TIS-$H_2O$ (95:2.5:2.5 v\v\v) for 2 h at room temperature. The filtrate was precipitated in cold methyl tert-butyl ether (20 mL). The precipitated crude peptide was collected by centrifugation, then re-dissolved in 10 mL of $H_2O$-acetonitrile (1:1, v\v) and lyophilized. Purification was performed by RP-HPLC as described in 1.2.

Ahx-GWWG-$N_3$ LCMS (ESI+): calc = 2279.17, found: 2279.15
CTP-$N_3$ LCMS (ESI+): calc = 2336.25, found: 2336.27; 1167.62
L4+-$N_3$ LCMS (ESI+): calc = 513.66, found: 512.72

*MethylTetratzine-modified peptides (MetTET-peptides)*

Azide-functionalized peptides (CTP-$N_3$, GWWG-$N_3$,1 equiv) dissolved in DMSO were reacted with DBCO-PEG3-methyltetrazine (1 equiv) in DMSO at RT for 1 h. Reaction completion was confirmed by LC-MS analysis. The product was purified by RP-HPLC as described in 1.2. Fraction containing the product were combined and lyophilized.

CTP-MetTET LCMS (ESI+): calc = 2870, found 2870

GWWG-MetTET LCMS (ESI+): calc = 2814, found 2814

*Lipid-modified positive linker L4+ (C16-L4+ and C22-L4+)*

Linker L4+ was mixed with C16-NHS or activated HBTU activated DCA in 1:3 molar ratio in DMF/pyridine 3:1 (v:v) at 50°C and left overnight. After reaction came to completion the crude product was further used without purification.

C16-L4+-$N_3$ LCMS (ESI+): calc [M+H] = 580, found 580

C22-L4+-$N_3$ LCMS (ESI+): calc [M+H] = 664, found 664

**Synthesis of 5′-conjugates**

Sod1 oligonucleotide containing 5′-bicyclo[6.1.0]non-4-yne group (5′-BCN-Sod1) was reacted with ligand containing azide group by mixing both entities in 1:1 ratio in water. After reaction completion the 5′-Sod1 conjugate were isolated by RP-HPLC as described in 1.2. Fractions containing product were freeze-dried.

5′-C16-L1-Sod1 LCMS (ESI-): calc = 7857, found 7857 $t_R$=3.3 min
5′-C16-PEG4-L1-Sod1 LCMS (ESI-): calc = 8104, found 8104, $t_R$= 3.3 min
5′-C16-PEG2000-L1-Sod1 LCMS (ESI-): calc = 10850, found 10850, $t_R$= 3.5 min

5′-C16-L4+-Sod1 LCMS (ESI-): $[M-4H]^{4-}$ calc = 1993, found 1993, $t_R$ = 3.4 min
5′-C16OOH–L1-Sod1 LCMS (ESI-): $[M-5H]^{5-}$ calc = 1576, found 1576, $t_R$ = 2.0 min
5′-C22-L1-Sod1 LCMS (ESI-): $[M-4H]^{4-}$ calc = 1984, found 1984, $t_R$ = 4.5 min
5′-C22-L4+-Sod1 LCMS (ESI-): $[M-5H]^{5-}$ calc = 1611, found 1611, $t_R$ = 4.1 min
5′-C20OOH–L1-Sod1 LCMS (ESI-): $[M-5H]^{5-}$ calc 1587, found 1585, $t_R$ = 2.8 min
5′-LCA-L1-Sod1 LCMS (ESI-): $[M-4H]^{4-}$ calc 1993, found 1993, $t_R$ = 3.0 min
5′-di(C16)-L2-Sod1 LCMS (ESI-): [M-1] calc = 8647, found 8647, $t_R$ = 5,2 min
5′-di(C16-PEG4)-L2-Sod1 LCMS (ESI-): [M-1] calc = 9150, found 9150, $t_R$= 5.0 min
5′-DOPE-PEG2000-Sod1 LCMS (ESI-): calc = 10211, found 10211, $t_R$= 6.6 min
5′-CTP-Sod1 LCMS (ESI): [M] calc = 9735, found 9735, $t_R$ = 1.9 min

Sod1 oligonucleotide containing 5′-bicyclo[6.1.0]non-4-yne group (5′-BCN-Sod1) (1 equiv, 0.0026 mmol, 20mg) in 180 uL of water was mixed with azido-PEG3-azide (2 equiv, 0.0051 mmmol, 1.01 mg) in 20 uL DMSO:H2O 1:1, for 1 hour at RT. The product was purified by RP-HPLC as described in 1.2. Fractions containing product were lyophilized.

5′-N3-Sod1 LCMS (ESI-): [M-1] calc = 7601, found -5/5 = 1519

Sod1 oligonucleotide containing azido moiety at 5′-end (5′-N3-Sod1) (1 equiv, 0.00136 mmol, 11 mg) in 400 uL of water was mixed with DOPE-DBCO (1 equiv, 0.00136 mmol, 1.43 mg) in 20 uL DMSO for 3 h at RT. After completion of reaction the product was lyophilized.

5′-DOPE-Sod1 LCMS (ESI-): [M-1] calc = 8632, found -5/5 = 1725, $t_R$= 6.2 min

Malat1 oligonucleotide containing 6-aminohexanyl moiety at 5′-end was dissolve in bicarbonate buffer ph 8.8 and DBCO N-hydroxysuccinimide ester (5 equiv, 0.0012 mmol, 6 mg) in 10uL DMSO was added. Reaction was agitated at RT overnight. After completion of the reaction product was isolated by RP-HPLC as described in 1.2. Product 5′-DBCO-Malat1 was lyophilized.

5′-DBCO-Malat1 LCMS (ESI-): [M-1] calc = 6039.82, found 6039.82

Malat1 oligonucleotide containing 5′- dibenzocyclooctyne group (5′-DBCO-Malat1) was reacted with ligand containing azide group by mixing both entities in 1:1 ratio in water. After reaction completion the 5′-Malat1 conjugate were isolated by RP-HPLC as described in 1.2. Fractions containing product were freeze-dried.

5′-C16-Malat1 LCMS (ESI-): [M-1] calc = 6497.12, found 6497.21, $t_R$ = 3.67 min

5′-C22-Malat1 LCMS (ESI-): [M-1] calc = 6582.09, found 6582.12, $t_R$ = 5.14 min

**Synthesis of 3′,5′-bis-conjugates.**

Sod1 oligonucleotide containing 3′,5′-bis(bicyclo[6.1.0]non-4-yne) modification (3′,5′-BCN-Sod1) was reacted with ligand containing azide group by mixing both entities in 1:2 ratio at RT in water for 2h. After reaction completion, the 3′,5′-Sod1 bis-conjugate was purified by RP-HPLC as described in 1.2. Fractions containing product were lyophilized.

3′,5′-bis-(C16-L1)-Sod1, LCMS (ESI-): [MS] calc = 8914, found 8914, $t_R$ = 4.9 min
3′,5′-bis-(C16-PEG4-L1)-Sod1, LCMS (ESI-): [MS] calc = 8914, found 8914, $t_R$ = 4.9 min
3′,5′-bis-(C16-L4+)-Sod1 LCMS (ESI-): calc = 9165, found 9165, $t_R$ = 3.8 min
3′,5′-bis-(C16OOH-L1)-Sod1 LCMS (ESI-): calc = 8968, found 8968, $t_R$ = 3.5 min
3′,5′-bis-(C22-L1)-Sod1, LCMS (ESI-): [MS] calc = 9083, found 9083, $t_R$ = 6.3 min
3′,5′-bis-(C22-L4+)-Sod1 LCMS (ESI-): calc = 9329, found 9329, $t_R$ = 5.6 min
3′,5′-bis-(C20OOH-L1)-Sod1 LCMS (ESI-): calc = 9088, found 9088, $t_R$ = 3.9 min
3′,5′-bis-(LCA-L1)-Sod1 LCMS (ESI-): calc = 9152, found 9152, $t_R$ = 3.8 min
3′,5′-bis(diC16-PEG4-L1)-Sod1 LCMS (ESI-): [MS] calc 9407, found 9407, $t_R$ = 5.6 min
3′,5′-bis-(DOPE-PEG2000)-Sod1 LCMS (ESI-): calc = 13622, found 13622, (not purified)
3′,5′-bis-CTP-Sod1 LCMS (ESI-): [M-1] calc = 12762, found 12762, $t_R$ = 5.6 min

Sod1 oligonucleotide containing bicyclo[6.1.0]non-4-yne group at 3′ and 5′ ends (3′,5′-BCN-Sod1) (1 equiv, 0.0024 mmol, 20mg) in 180 uL of water was mixed with azido-PEG3-azide (4 equiv, 0.0096 mmol, 1.92 mg) in 20 uL DMSO:H2O 1:1, for 1 hour at RT. The product was purified by RP-HPLC as described in 1.2. Fractions containing product were lyophilized.

3′,5′-bis-N3-Sod1 LCMS (ESI-): [M-1] calc = 8400, found -5/5 = 1679

Sod1 oligonucleotide containing azido moiety at 3′ and 5′ ends (3′,5′-bis-N3-Sod1) (1 equiv, 0.0023 mmol, 20 mg) in 400 uL of water was mixed with DOPE-DBCO (1 equiv, 0.00136 mmol, 2.4 mg) in 20 uL DMSO for 3 h at RT. After completion of reaction the product was lyophilized.

3′,5′-bis-DOPE-Sod1 LCMS (ESI-): [M-1] cal. = 10464, found -5/5 = 2091

Malat1 oligonucleotide containing 6-aminohexanyl moiety at 3′ and 5′-end was dissolved in bicarbonate buffer ph 8.8 and DBCO N-hydroxysuccinimide ester (6 equiv, 0.0012 mmol, 6 mg) in 10uL DMSO was added. Reaction was agitated at RT overnight. After completion of the reaction product was isolated by RP-HPLC as described in 1.2. Product 3′,5′-DBCO-Malat1 was lyophilized.

3′,5′-DBCO-Malat1 LCMS (ESI-): [M-1] calc = 6753.12, found 6753.21

Malat1 oligonucleotide containing dibenzocyclooctyne groups at 3′ and 5′ ends (3′,5′-DBCO-Malat1) was reacted with C16-PEG3-azide by mixing both entities in 1:3 ratio in water. After reaction completion the 3′,5′-Malat1 bis-conjugate was isolated by RP-HPLC as described in 1.2. Fractions containing product were freeze-dried.

3′,5′-bis-C16-Malat1 LCMS (ESI-): [M-1] calc = 7653.5, found 7653.23 $t_R$ = 5.77

**Synthesis of 3′,5′ hetero-functional conjugates**

Sod1 oligonucleotide containing 6-aminohexanyl moiety at 3′-end and bicyclo[6.1.0]non-4-yne) modification at 5′-end was dissolve in bicarbonate buffer ph 8.8 and *trans*-cyclooctene N-hydroxysuccinimide ester (5 equiv, 0.0012 mmol, 6 mg) in 10uL DMSO was added. Reaction was agitated in darkness at RT overnight. After completion of the reaction product was isolated by RP-HPLC as described in 1.2. Product 3′-TCO-5′-BCN-Sod1 was lyophilized and protected from light.

3′-TCO-5′-BCN-Sod1 LCMS (ESI-): [M-1] calc = 7979, found -5/5 = 1594

Sod1 oligonucleotide containing *trans*-cyclooctene moiety (TCO) at 3′-end and 5′-bicyclo[6.1.0]non-4-yne group (BCN) at 5′-end (3′-TCO-5′-BCN-Sod1) was first mixed with 1 equivalent azide modified ligand (C16-PEG4-$N_3$) in water for 3h at RT. After formation of 5′-conjugate 1.1 equivalent of methyltetrazine modified ligand (CTP-metTET, GWWG-metTET) was added for 1h at RT. After reaction completion the mixed 3′,5′-conjugate was purified by RP-HPLC as described in 1.2. Product was lyophylized

3′-TCO-5′-C16-Sod1 LCMS (ESI-): [M-1] calc = 8683, found 8683,

3′-CTP-5′-C16-PEG4-Sod1 LCMS (ESI-): [M-1] calc = 11553, found 11553, $t_R$ = 12.18 min
3′-GWWG-5′-C16-PEG4-Sod1 LCMS (ESI-):[M-1] calc = 11497, found 11497, $t_R$ = 12.14 min

3′-GWWG-5′-CTP-Sod1 LCMS (ESI-): [M-1] calc = 13129, found 13129, $t_R$ = 6.18 min

#### HPLC CHROMATOGRAMS AND MS DATA

*Cardiomyocyte targeting peptide - $K_{N3}$WLSEAGPVVTVRALRGTGSW*

MS

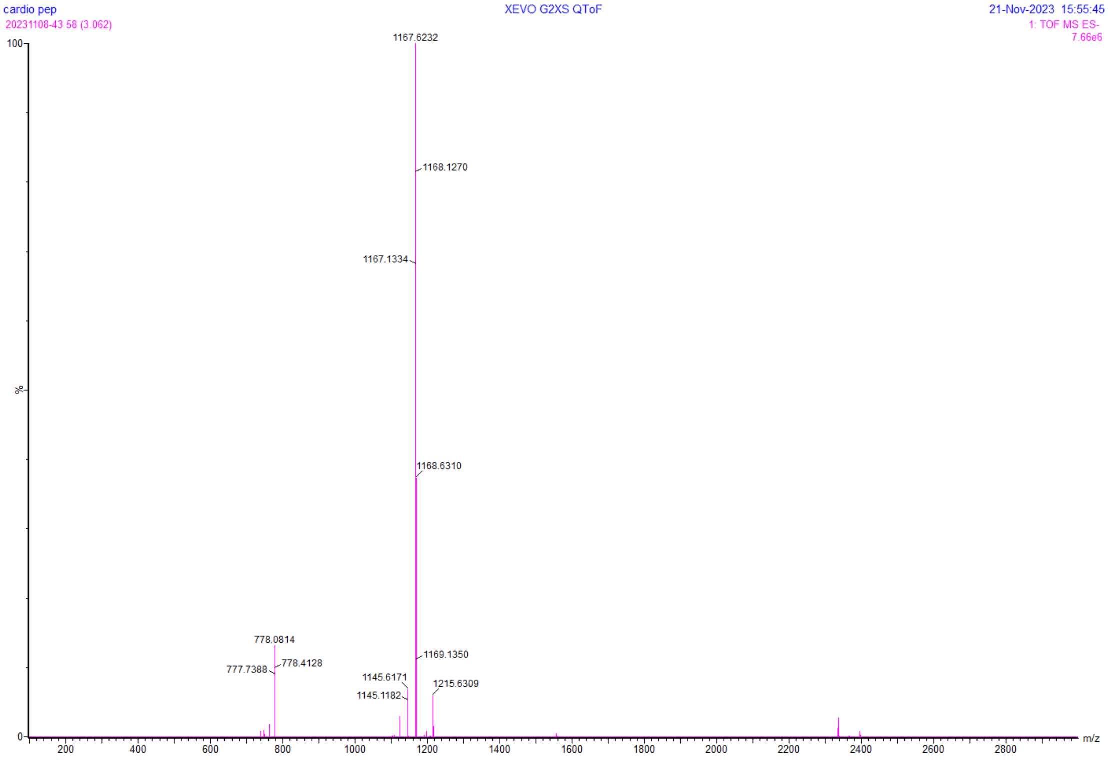


*Ahx-GWWG peptide $K_{N3}$WGELLEALAELLEAhxGWWG*

MS

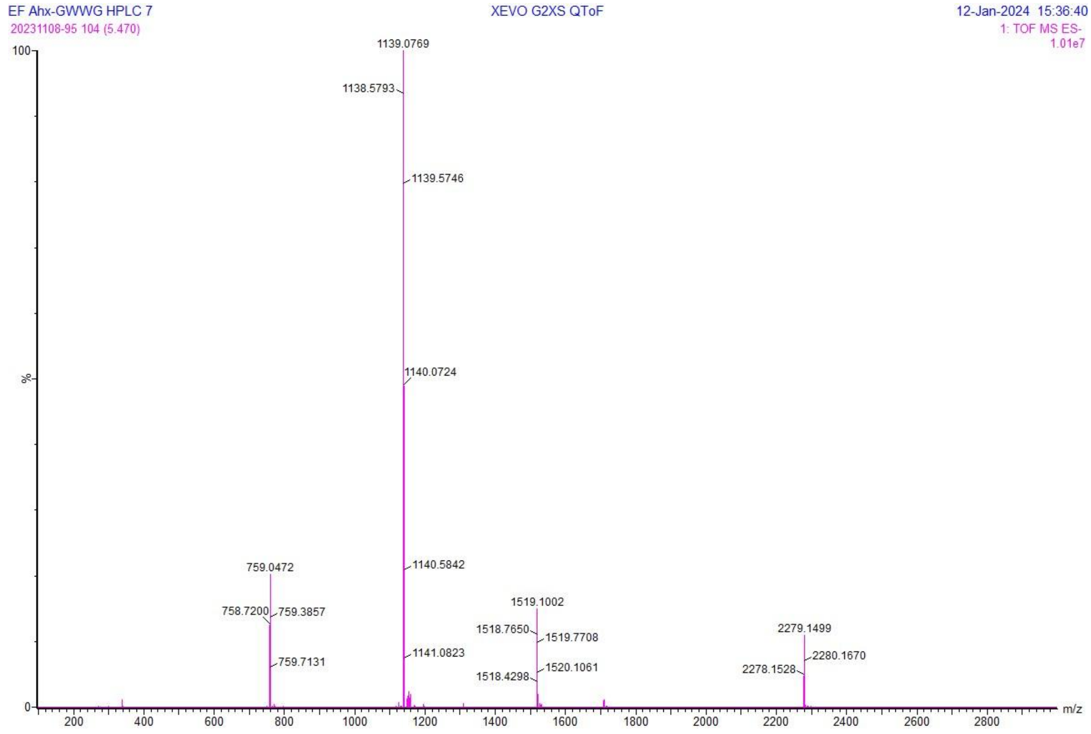


*Positively charged linker – K-K(Me)3KN3*

MS

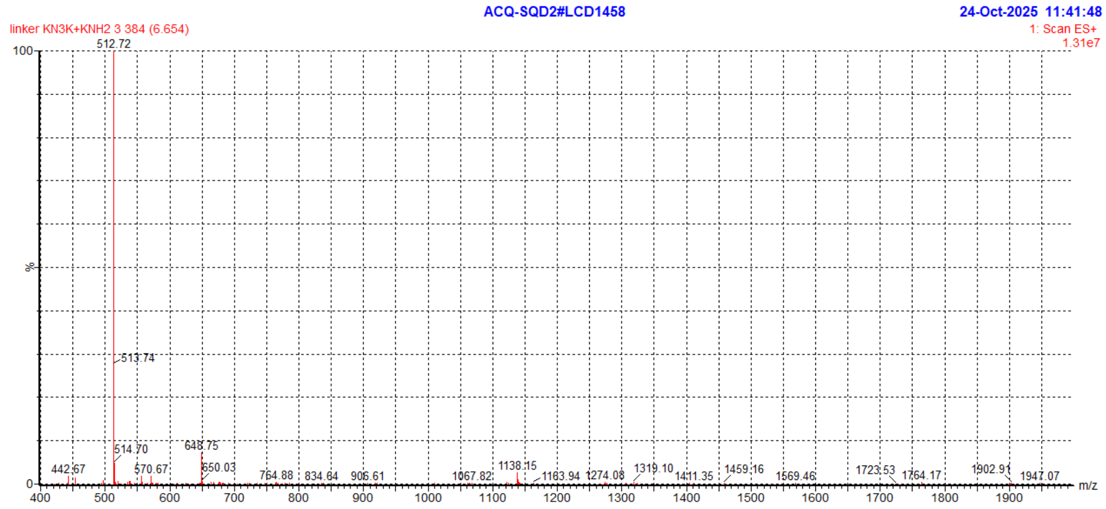

#### siRNA conjugates

##### Mono-LCA

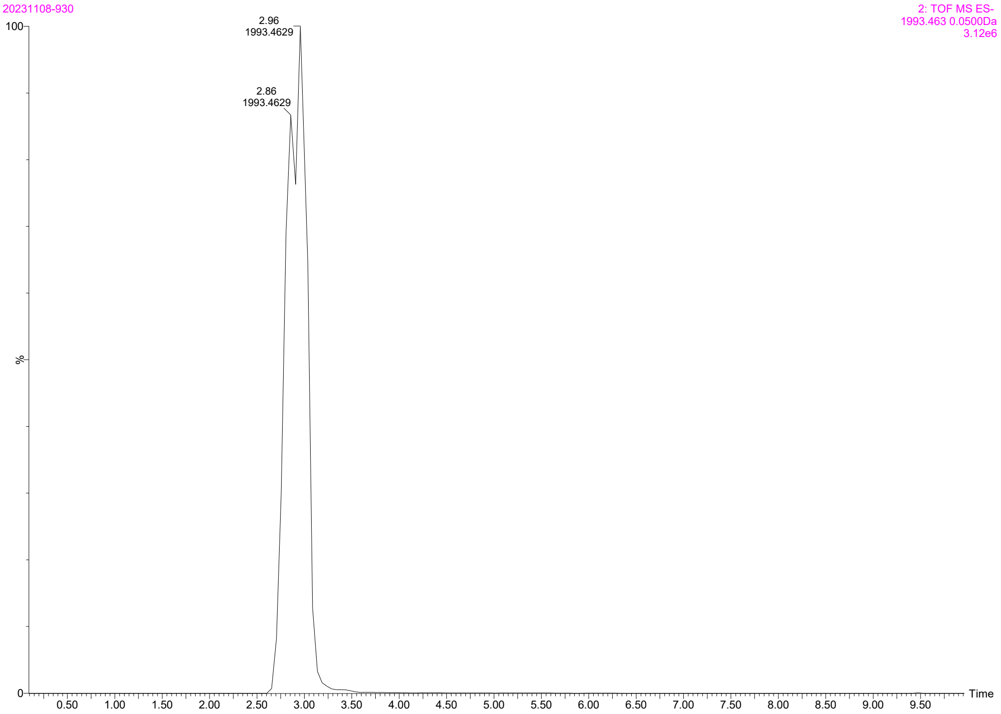


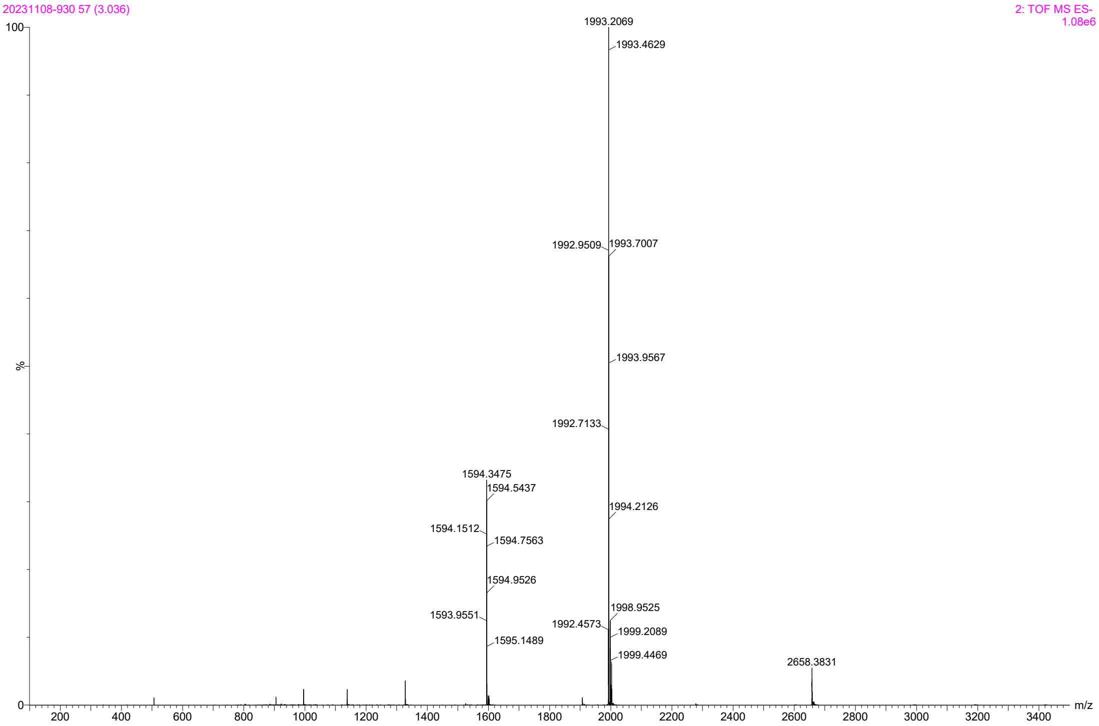

Bis-LCA

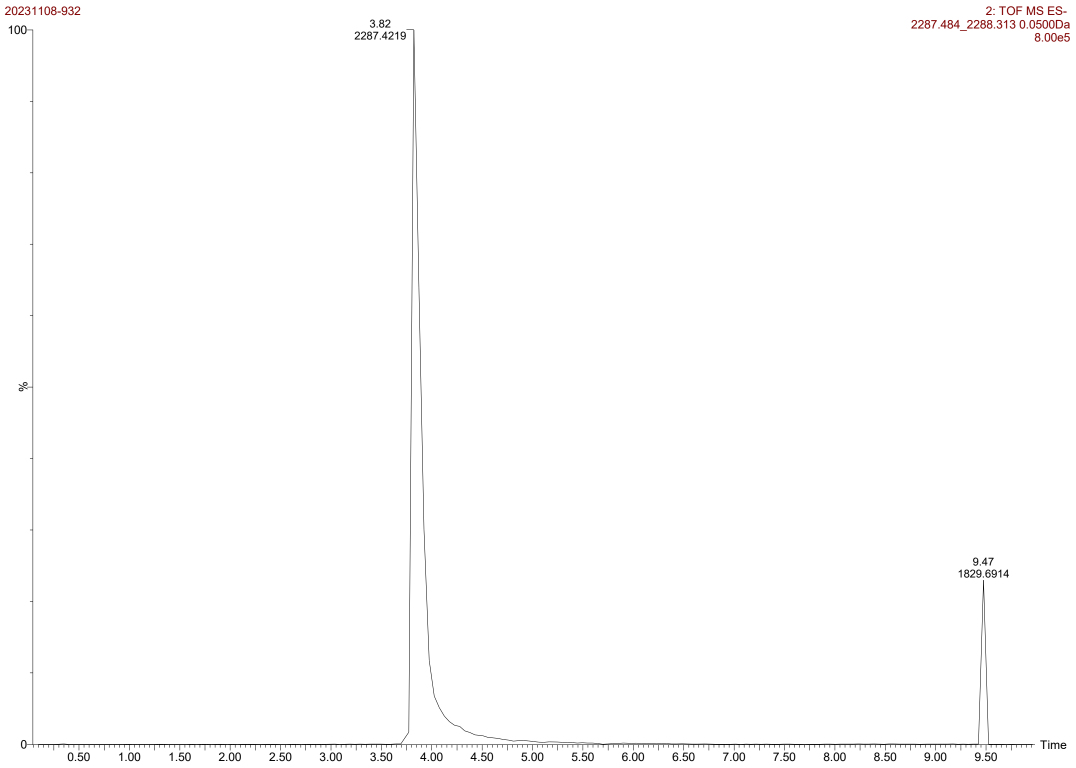
20231108-932
2: TOF MS ES-
2287.484_2288.313 0.0500Da
8.00e5
3.82
2287.4219
9.47
1829.6914
%
Time

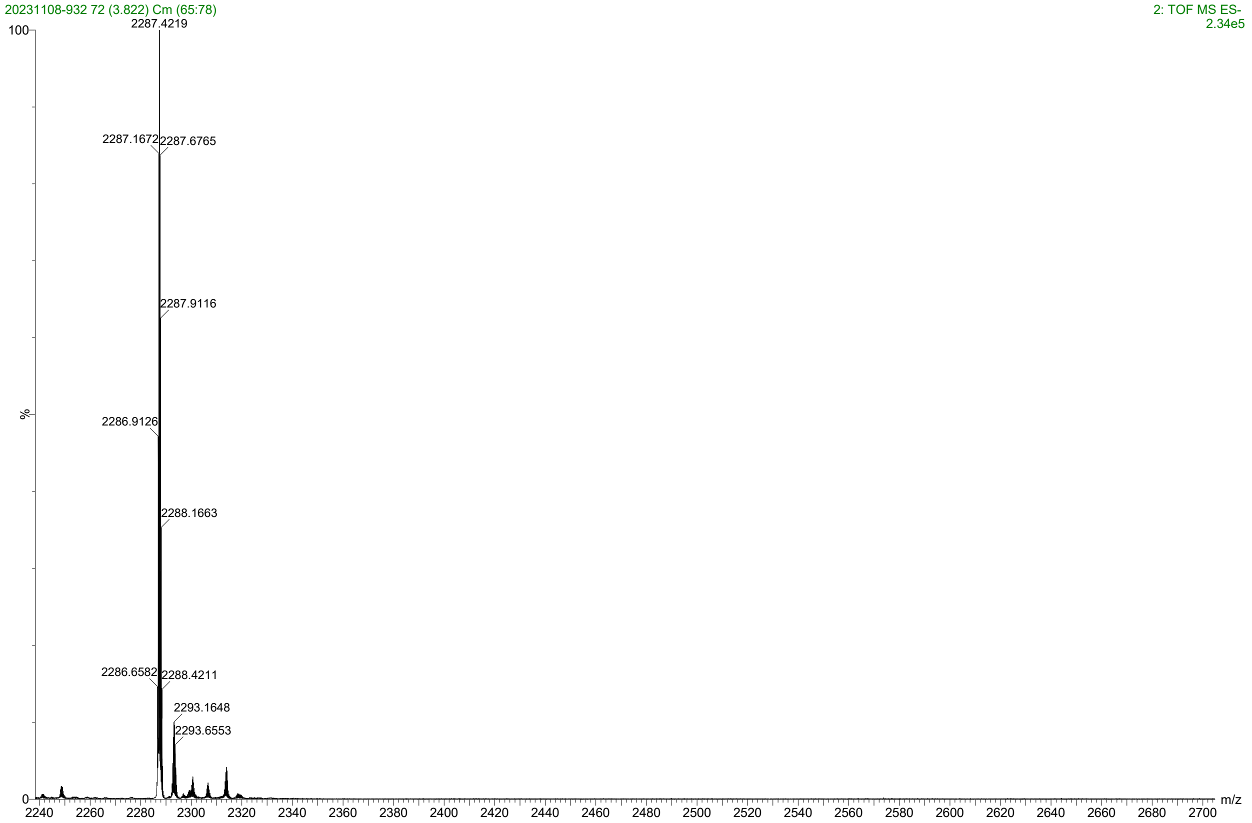
20231108-932 72 (3.822) Cm (65:78)
2: TOF MS ES-
2.34e5
2287.4219
2287.1672
2287.6765
2287.9116
2286.9126
2288.1663
2286.6582
2288.4211
2293.1648
2293.6553
%
m/z

Mono-DOPE

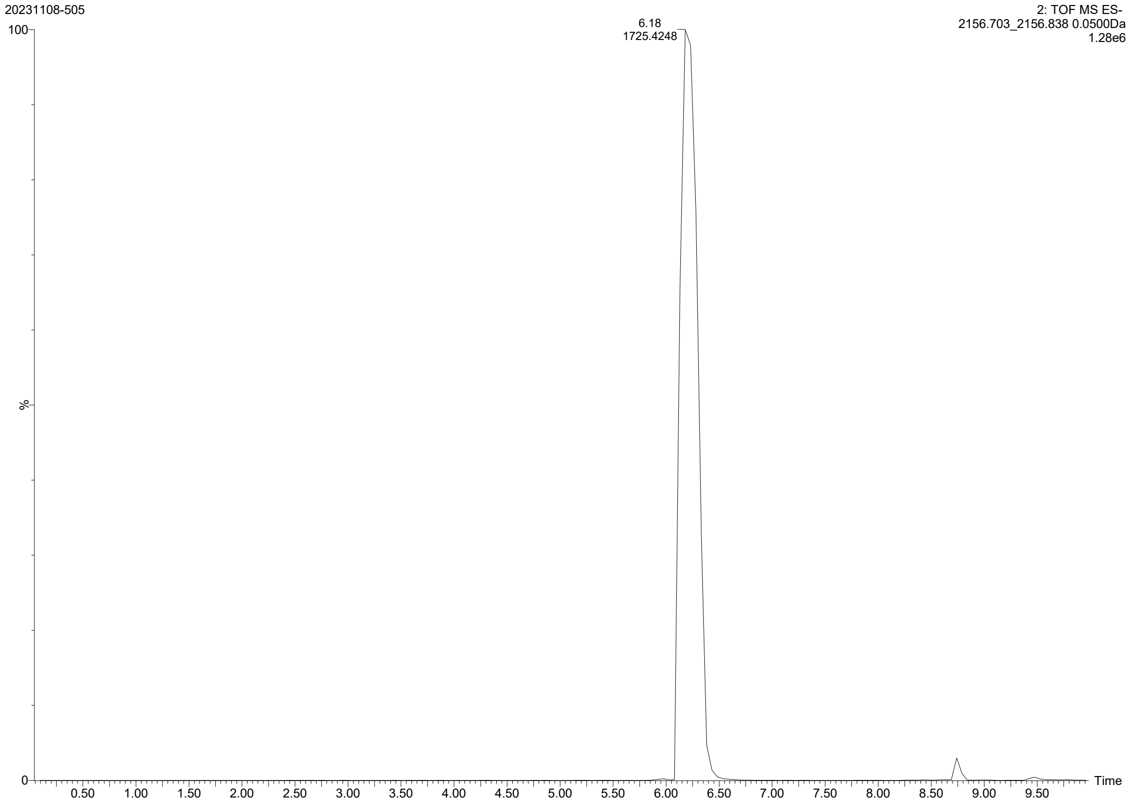

20231108-505
2: TOF MS ES-
2156.703_2156.838 0.0500Da
1.28e6
100
6.18
1725.4248
%
0
0.50
1.00
1.50
2.00
2.50
3.00
3.50
4.00
4.50
5.00
5.50
6.00
6.50
7.00
7.50
8.00
8.50
9.00
9.50
Time


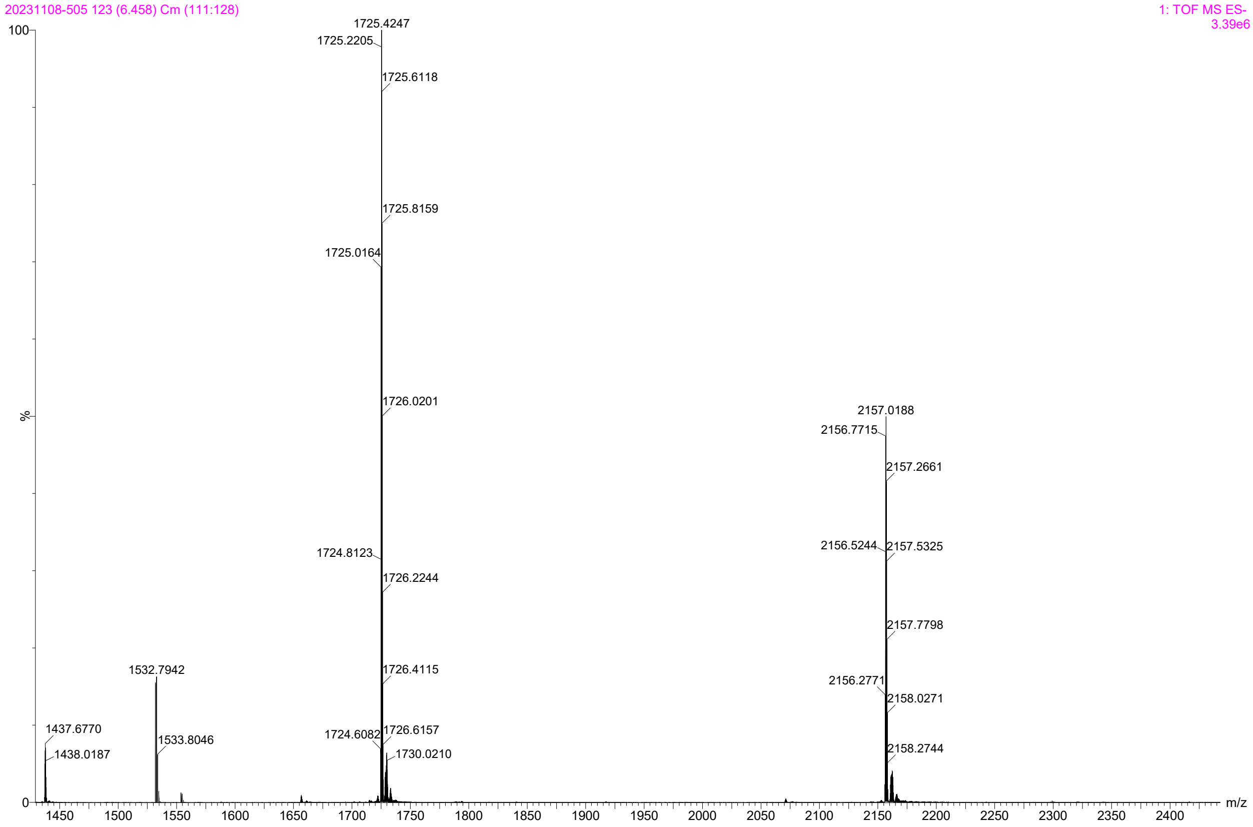

20231108-505 123 (6.458) Cm (111:128)
1: TOF MS ES-
3.39e6
100
1725.4247
1725.2205
1725.6118
1725.8159
1725.0164
1726.0201
1724.8123
1726.2244
1726.4115
1724.6082
1726.6157
1730.0210
2157.0188
2156.7715
2157.2661
2156.5244
2157.5325
2157.7798
2156.2771
2158.0271
2158.2744
1532.7942
1533.8046
1437.6770
1438.0187
%
0
1450
1500
1550
1600
1650
1700
1750
1800
1850
1900
1950
2000
2050
2100
2150
2200
2250
2300
2350
2400
m/z

Bis-DOPE

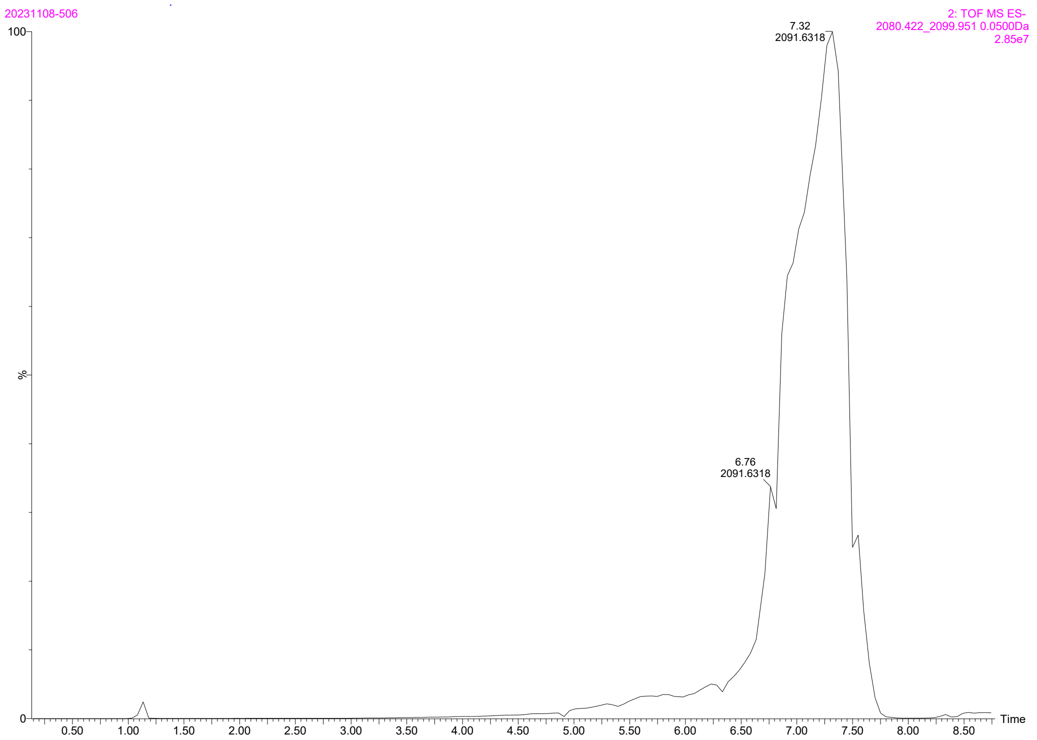

20231108-506
2: TOF MS ES-
2080.422_2099.951 0.0500Da
2.85e7
7.32
2091.6318
6.76
2091.6318
100
%
0
0.50
1.00
1.50
2.00
2.50
3.00
3.50
4.00
4.50
5.00
5.50
6.00
6.50
7.00
7.50
8.00
8.50
Time


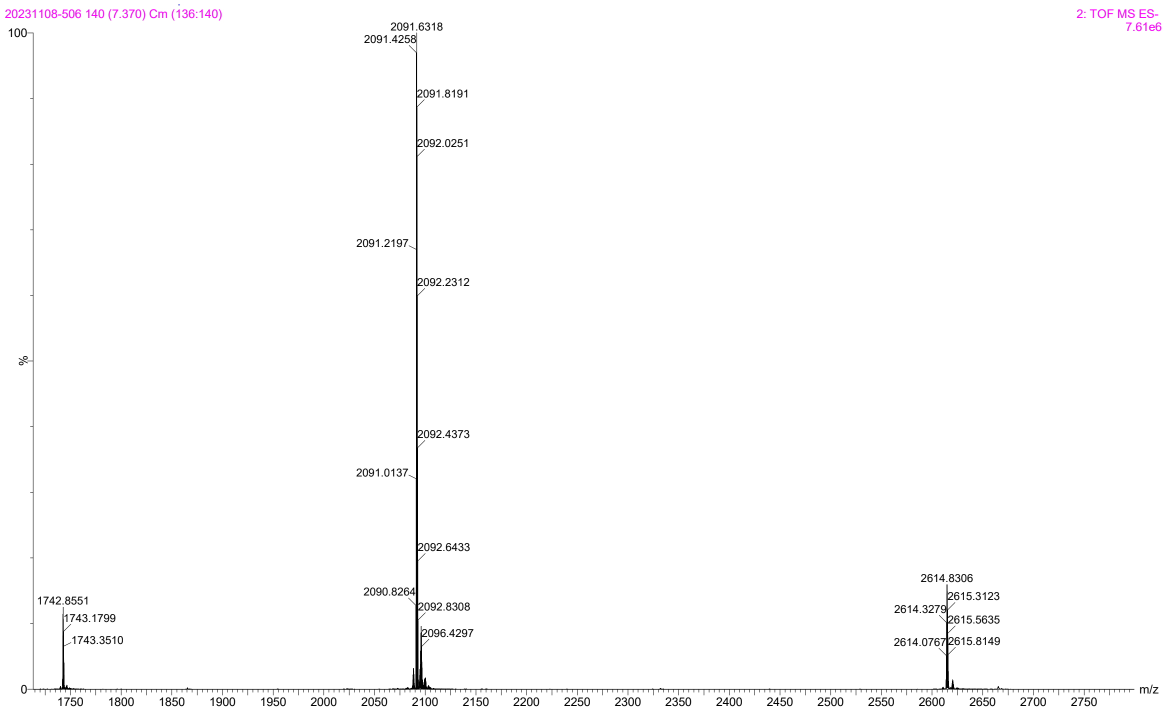

20231108-506 140 (7.370) Cm (136:140)
2: TOF MS ES-
7.61e6
2091.6318
2091.4258
2091.8191
2092.0251
2091.2197
2092.2312
2092.4373
2091.0137
2092.6433
2090.8264
2092.8308
2096.4297
1742.8551
1743.1799
1743.3510
2614.8306
2615.3123
2614.3279
2615.5635
2614.0767
2615.8149
100
%
0
1750
1800
1850
1900
1950
2000
2050
2100
2150
2200
2250
2300
2350
2400
2450
2500
2550
2600
2650
2700
2750
m/z

Mono-C16OOH

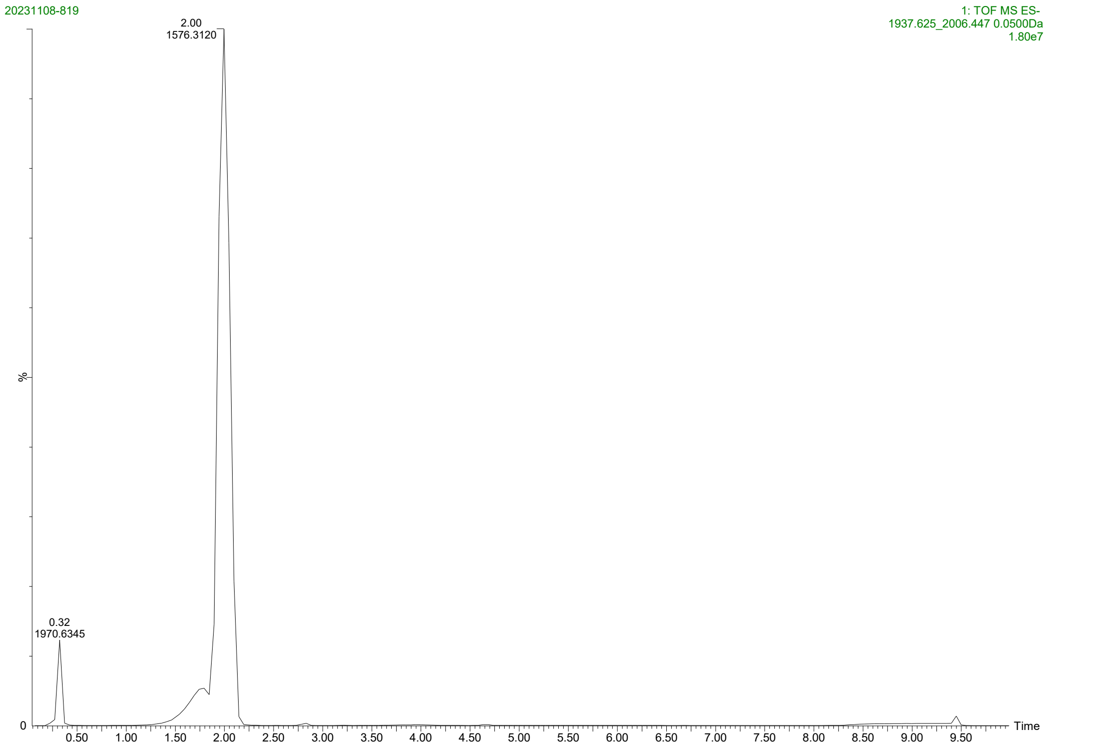
20231108-819
1: TOF MS ES-
1937.625_2006.447 0.0500Da
1.80e7
2.00
1576.3120
0.32
1970.6345
%
0
0.50
1.00
1.50
2.00
2.50
3.00
3.50
4.00
4.50
5.00
5.50
6.00
6.50
7.00
7.50
8.00
8.50
9.00
9.50
Time

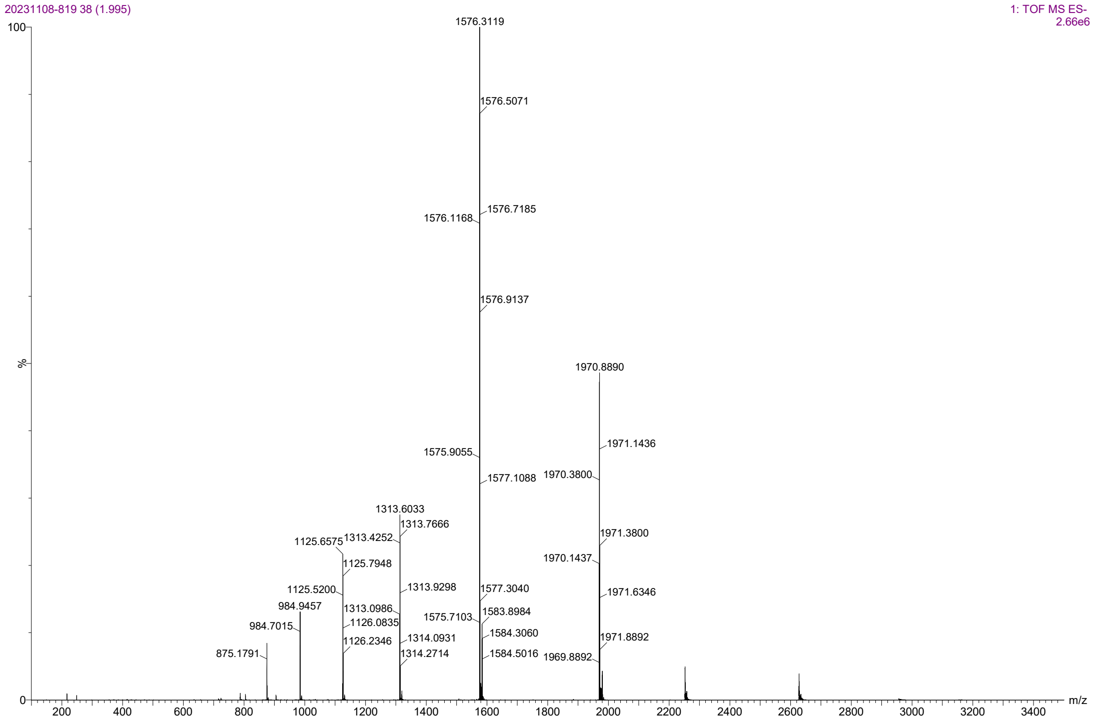
20231108-819 38 (1.995)
1: TOF MS ES-
2.66e6
100
%
0
1576.3119
1576.5071
1576.7185
1576.1168
1576.9137
1970.8890
1971.1436
1575.9055
1577.1088
1970.3800
1313.6033
1313.7666
1125.6575
1313.4252
1971.3800
1125.7948
1970.1437
1125.5200
1313.9298
1577.3040
1971.6346
984.9457
1313.0986
1583.8984
1126.0835
1575.7103
984.7015
1126.2346
1314.0931
1584.3060
1971.8892
875.1791
1314.2714
1584.5016
1969.8892
200
400
600
800
1000
1200
1400
1600
1800
2000
2200
2400
2600
2800
3000
3200
3400
m/z

### Mono-C20OOH

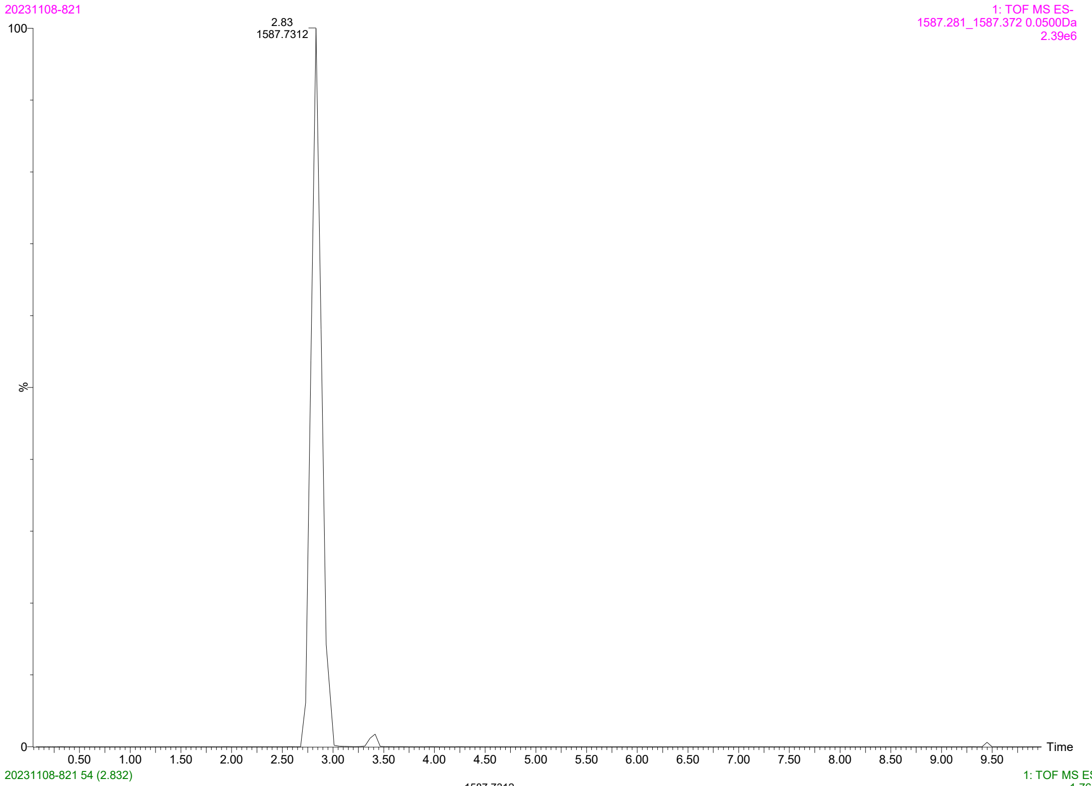

20231108-821
1: TOF MS ES-
1587.281_1587.372 0.0500Da
2.39e6
2.83
1587.7312
100
%
0
0.50 1.00 1.50 2.00 2.50 3.00 3.50 4.00 4.50 5.00 5.50 6.00 6.50 7.00 7.50 8.00 8.50 9.00 9.50
Time


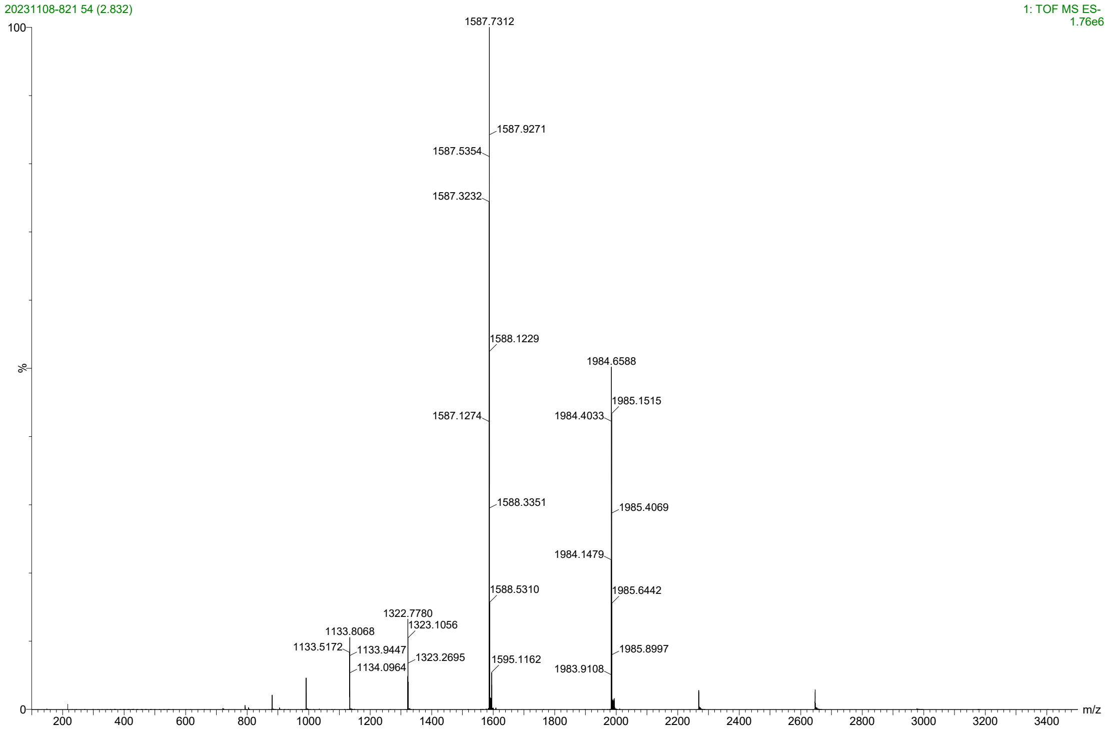

20231108-821 54 (2.832)
1: TOF MS ES-
1.76e6
1587.7312
1587.9271
1587.5354
1587.3232
1588.1229
1984.6588
1985.1515
1587.1274
1984.4033
1588.3351
1985.4069
1984.1479
1588.5310
1985.6442
1322.7780
1323.1056
1133.8068
1133.5172
1133.9447
1985.8997
1323.2695
1595.1162
1134.0964
1983.9108
100
%
0
200 400 600 800 1000 1200 1400 1600 1800 2000 2200 2400 2600 2800 3000 3200 3400
m/z

### Mono-C16

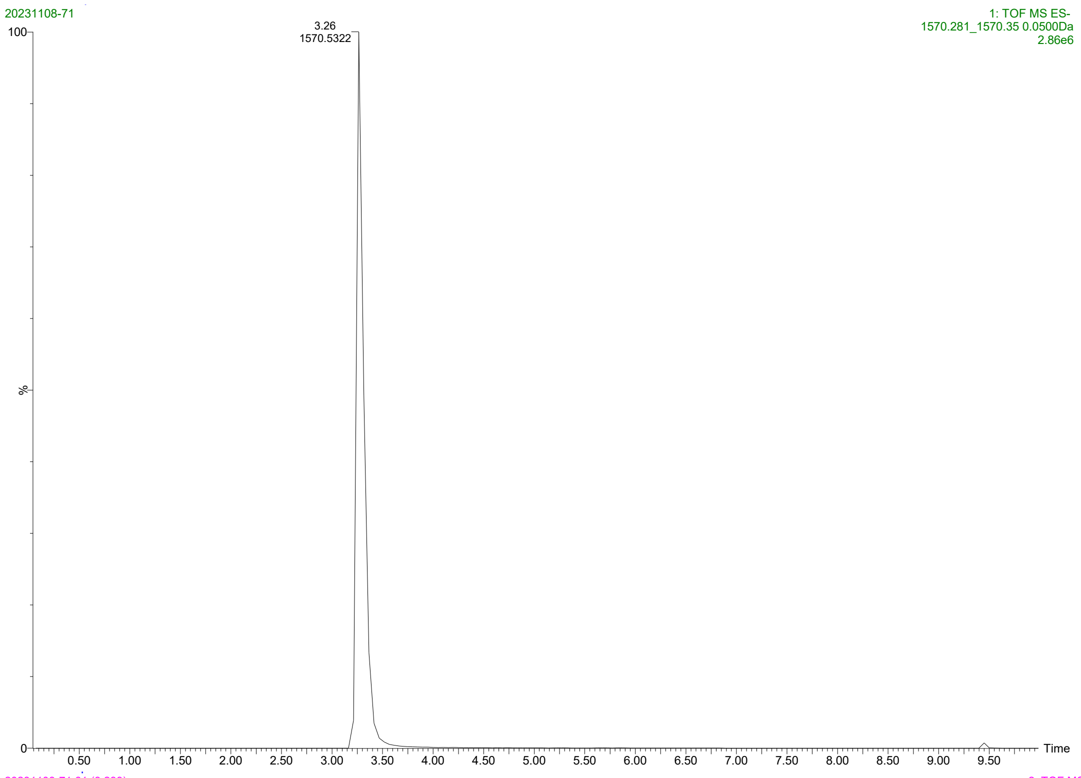

20231108-71
1: TOF MS ES-
1570.281_1570.35 0.0500Da
2.86e6
3.26
1570.5322
100
%
0
0.50 1.00 1.50 2.00 2.50 3.00 3.50 4.00 4.50 5.00 5.50 6.00 6.50 7.00 7.50 8.00 8.50 9.00 9.50
Time


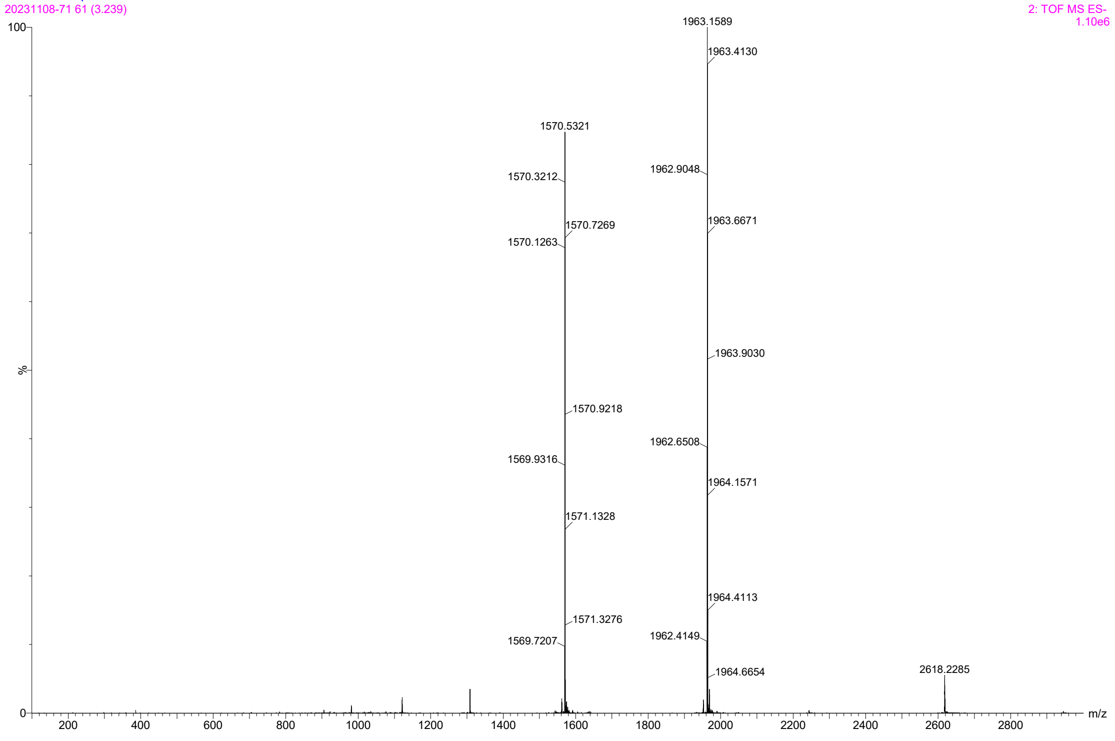

20231108-71 61 (3.239)
2: TOF MS ES-
1.10e6
1963.1589
1963.4130
1570.5321
1570.3212
1962.9048
1570.7269
1963.6671
1570.1263
1963.9030
1570.9218
1962.6508
1569.9316
1964.1571
1571.1328
1964.4113
1571.3276
1962.4149
1569.7207
1964.6654
2618.2285
100
%
0
200 400 600 800 1000 1200 1400 1600 1800 2000 2200 2400 2600 2800
m/z

Bis-C16OOH

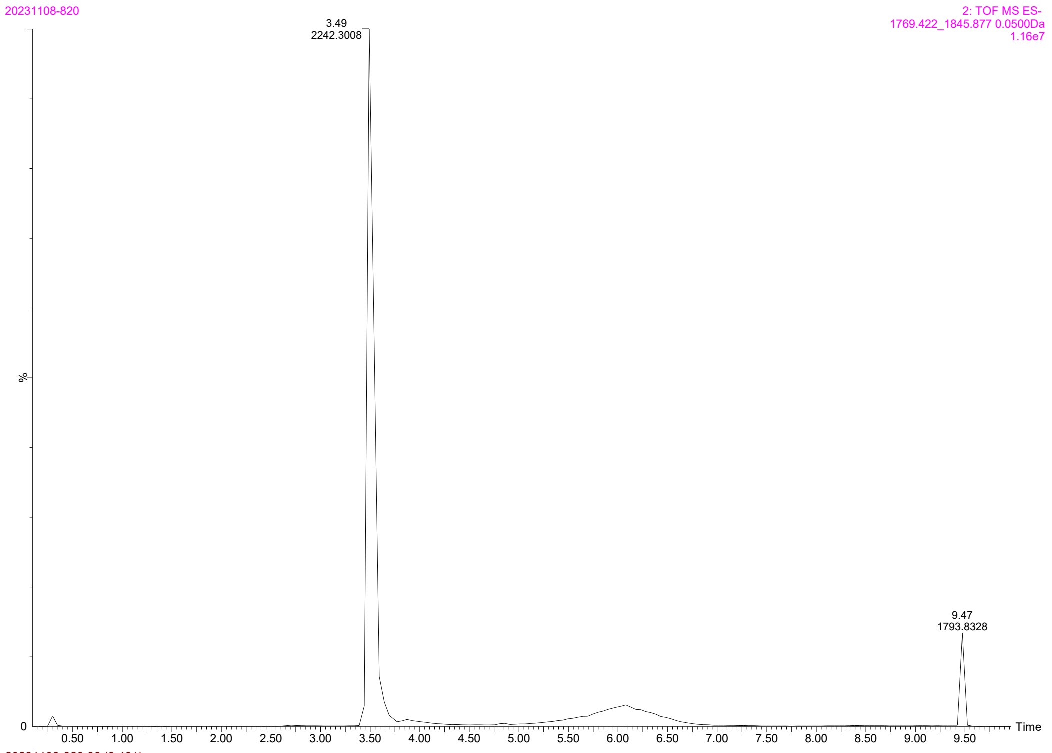

20231108-820
2: TOF MS ES-
1769.422_1845.877 0.0500Da
1.16e7
3.49
2242.3008
9.47
1793.8328
%
0
0.50
1.00
1.50
2.00
2.50
3.00
3.50
4.00
4.50
5.00
5.50
6.00
6.50
7.00
7.50
8.00
8.50
9.00
9.50
Time


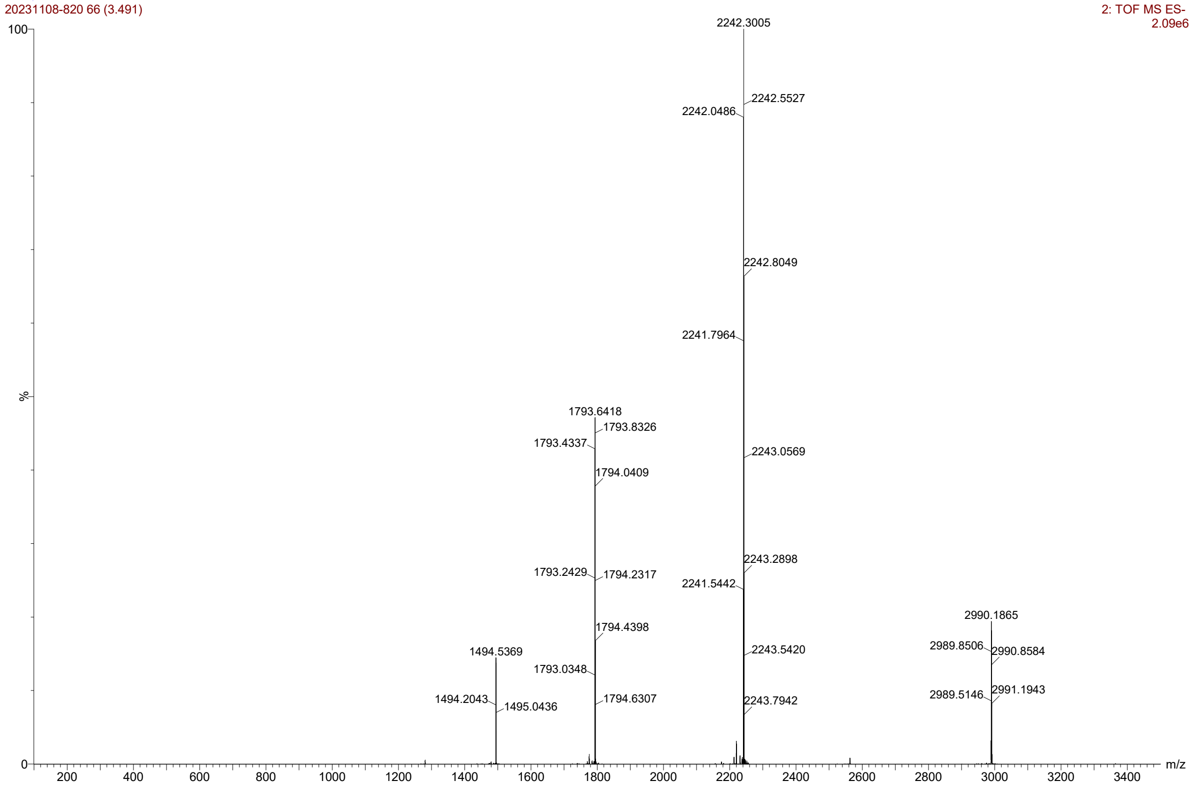

20231108-820 66 (3.491)
2: TOF MS ES-
2.09e6
100
%
0
2242.3005
2242.0486
2242.5527
2242.8049
2241.7964
2243.0569
2243.2898
2241.5442
2243.5420
2243.7942
1793.6418
1793.8326
1793.4337
1794.0409
1793.2429
1794.2317
1794.4398
1793.0348
1794.6307
1494.5369
1494.2043
1495.0436
2990.1865
2989.8506
2990.8584
2989.5146
2991.1943
200
400
600
800
1000
1200
1400
1600
1800
2000
2200
2400
2600
2800
3000
3200
3400
m/z

Bis-C20OOH

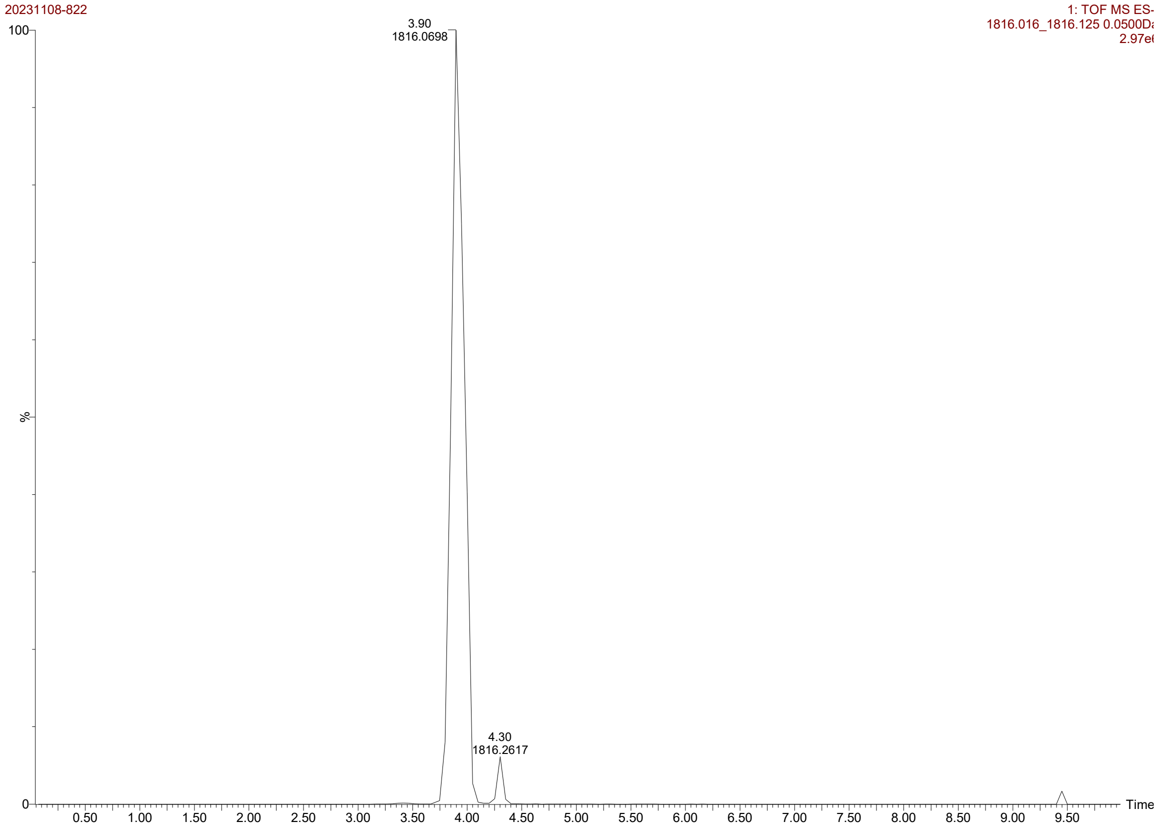

20231108-822
1: TOF MS ES-
1816.016_1816.125 0.0500Da
2.97e6
3.90
1816.0698
4.30
1816.2617
%
Time


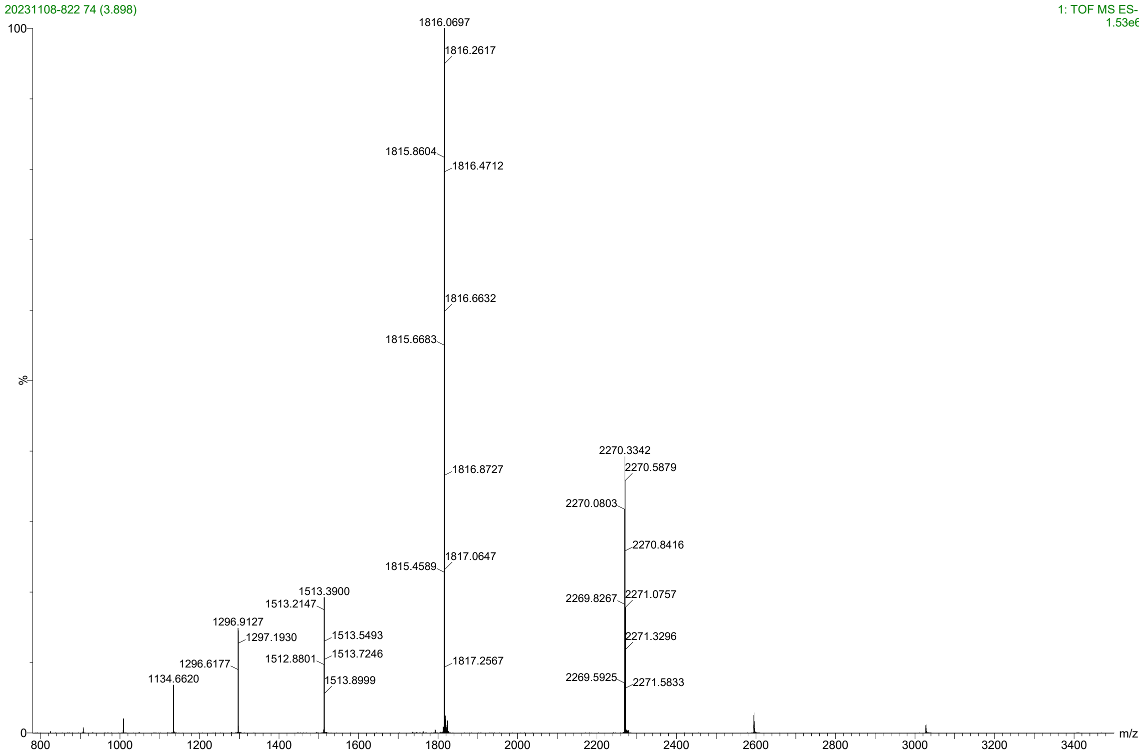

20231108-822 74 (3.898)
1: TOF MS ES-
1.53e6
1816.0697
1816.2617
1815.8604
1816.4712
1816.6632
1815.6683
2270.3342
1816.8727
2270.5879
2270.0803
2270.8416
1817.0647
1815.4589
1513.3900
1513.2147
2269.8267
2271.0757
1296.9127
1297.1930
1513.5493
2271.3296
1512.8801
1513.7246
1296.6177
1817.2567
1134.6620
1513.8999
2269.5925
2271.5833
%
m/z

### Mono-C22

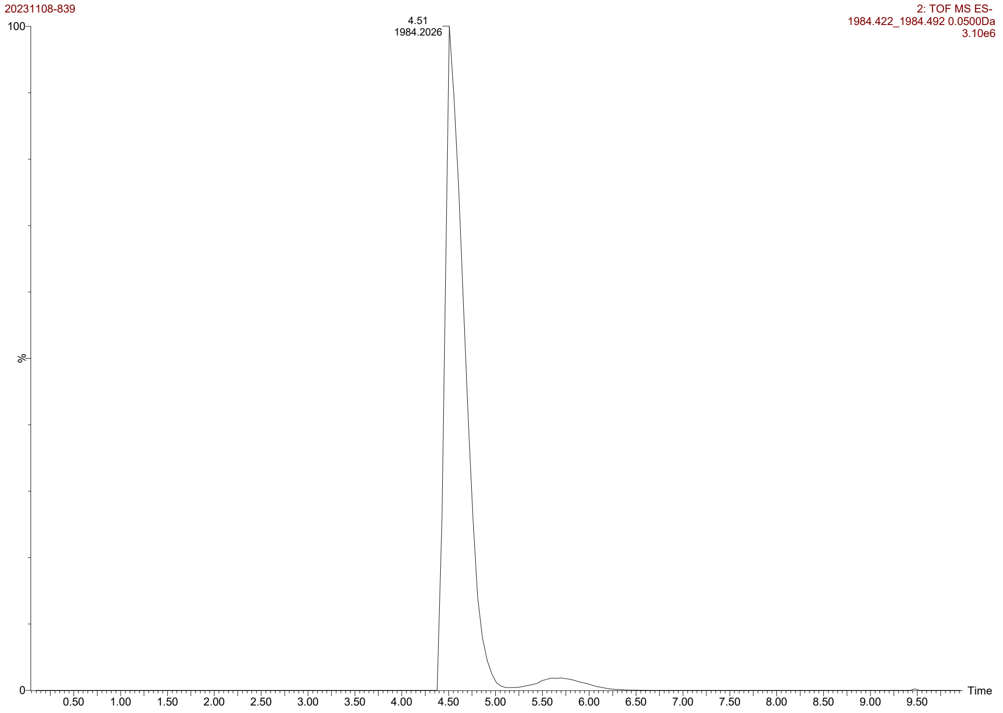

20231108-839
2: TOF MS ES-
1984.422_1984.492 0.0500Da
3.10e6
4.51
1984.2026
100
%
0
0.50
1.00
1.50
2.00
2.50
3.00
3.50
4.00
4.50
5.00
5.50
6.00
6.50
7.00
7.50
8.00
8.50
9.00
9.50
Time


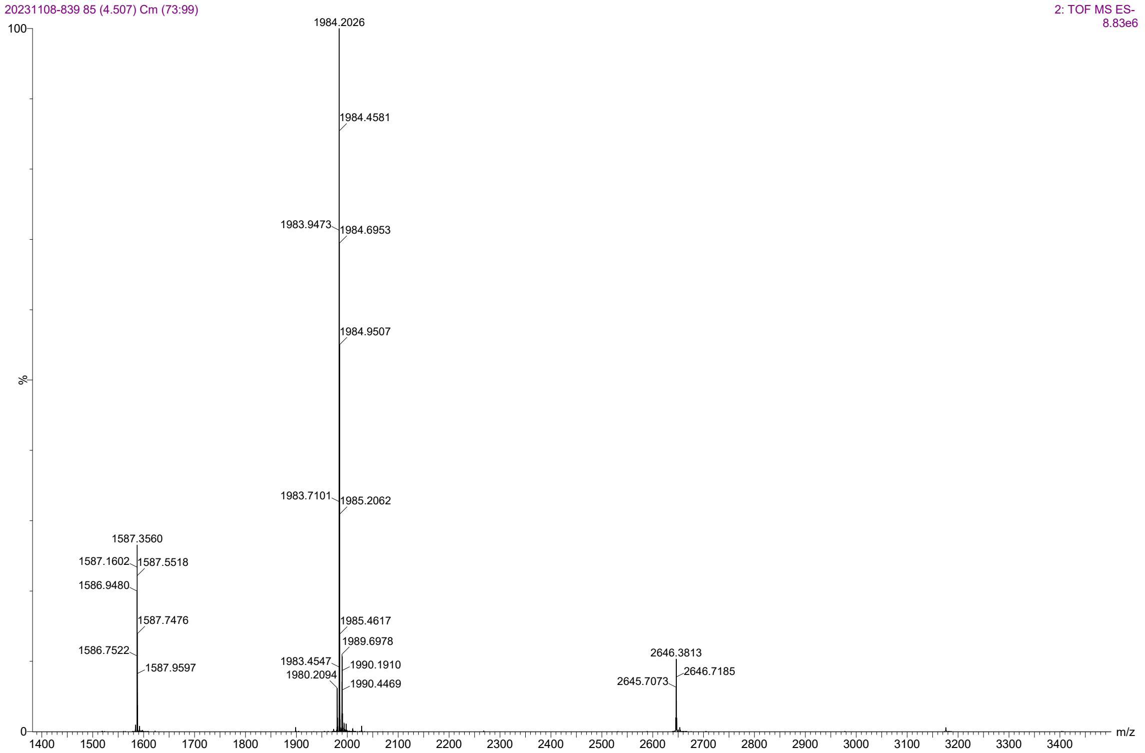

20231108-839 85 (4.507) Cm (73:99)
2: TOF MS ES-
8.83e6
1984.2026
1984.4581
1983.9473
1984.6953
1984.9507
1983.7101
1985.2062
1587.3560
1587.1602
1587.5518
1586.9480
1587.7476
1586.7522
1587.9597
1985.4617
1989.6978
1983.4547
1990.1910
1980.2094
1990.4469
2646.3813
2646.7185
2645.7073
100
%
0
1400
1500
1600
1700
1800
1900
2000
2100
2200
2300
2400
2500
2600
2700
2800
2900
3000
3100
3200
3300
3400
m/z

Bis-C16

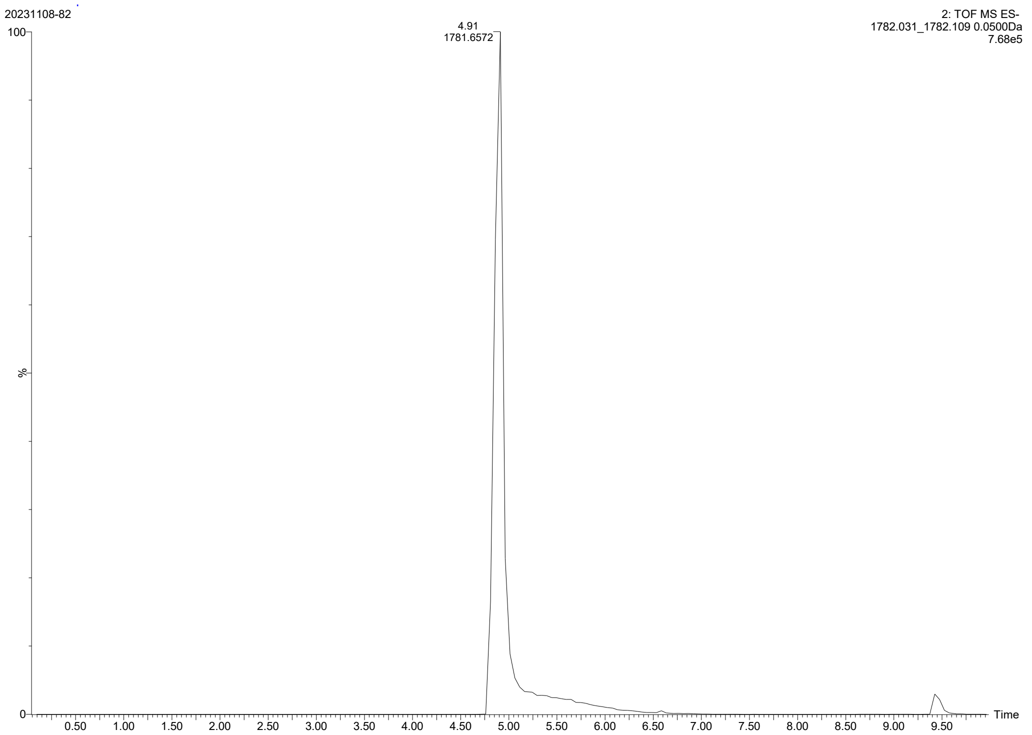
20231108-82
2: TOF MS ES-
1782.031_1782.109 0.0500Da
7.68e5
4.91
1781.6572
100
%
0
0.50
1.00
1.50
2.00
2.50
3.00
3.50
4.00
4.50
5.00
5.50
6.00
6.50
7.00
7.50
8.00
8.50
9.00
9.50
Time

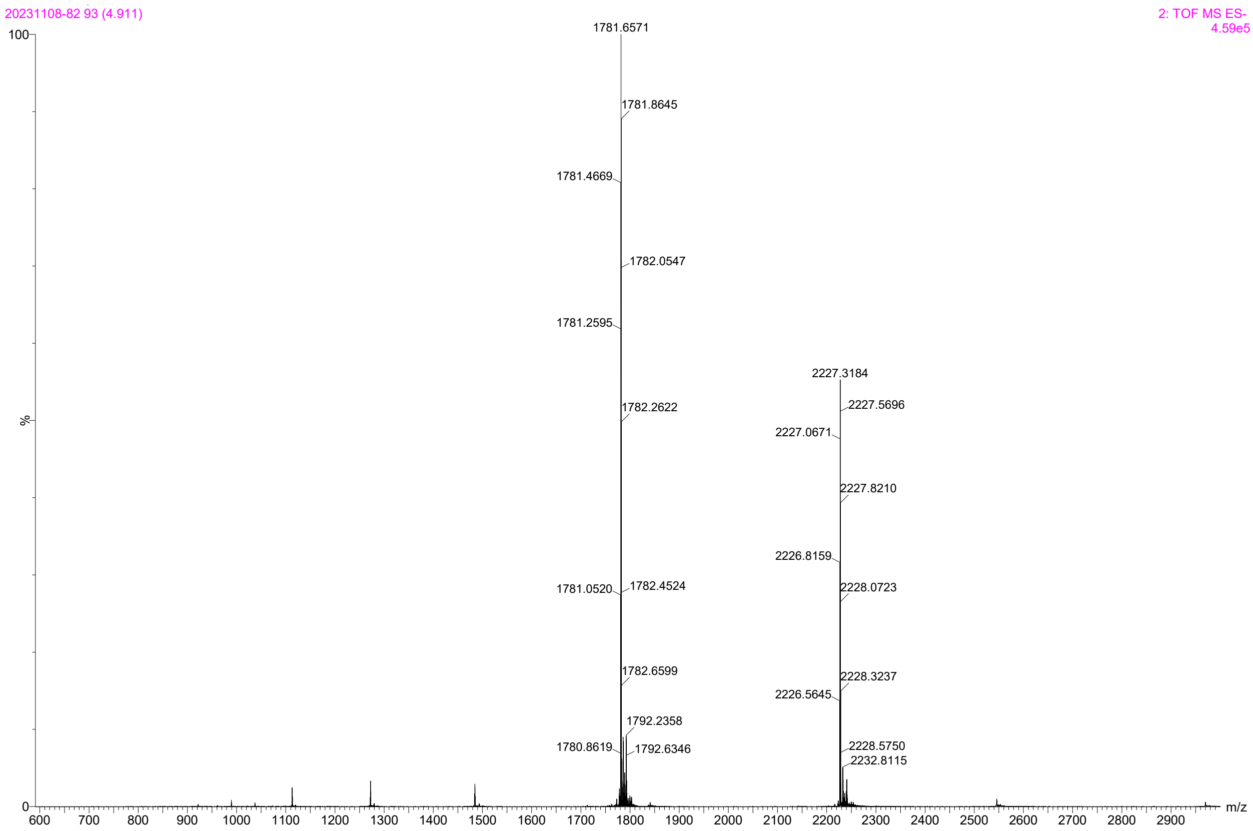
20231108-82 93 (4.911)
2: TOF MS ES-
4.59e5
1781.6571
1781.8645
1781.4669
1782.0547
1781.2595
1782.2622
1781.0520
1782.4524
1782.6599
1792.2358
1780.8619
1792.6346
2227.3184
2227.5696
2227.0671
2227.8210
2226.8159
2228.0723
2228.3237
2226.5645
2228.5750
2232.8115
100
%
0
600
700
800
900
1000
1100
1200
1300
1400
1500
1600
1700
1800
1900
2000
2100
2200
2300
2400
2500
2600
2700
2800
2900
m/z

Bis-C22

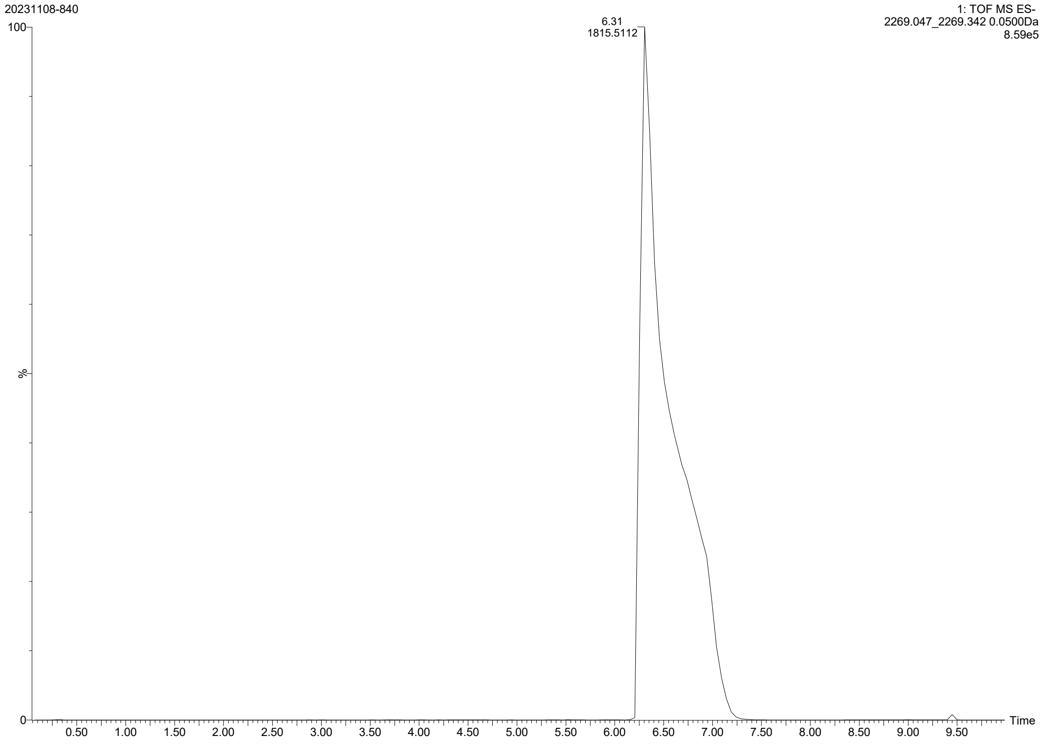
20231108-840
1: TOF MS ES-
2269.047_2269.342 0.0500Da
8.59e5
6.31
1815.5112
100
%
0
0.50
1.00
1.50
2.00
2.50
3.00
3.50
4.00
4.50
5.00
5.50
6.00
6.50
7.00
7.50
8.00
8.50
9.00
9.50
Time

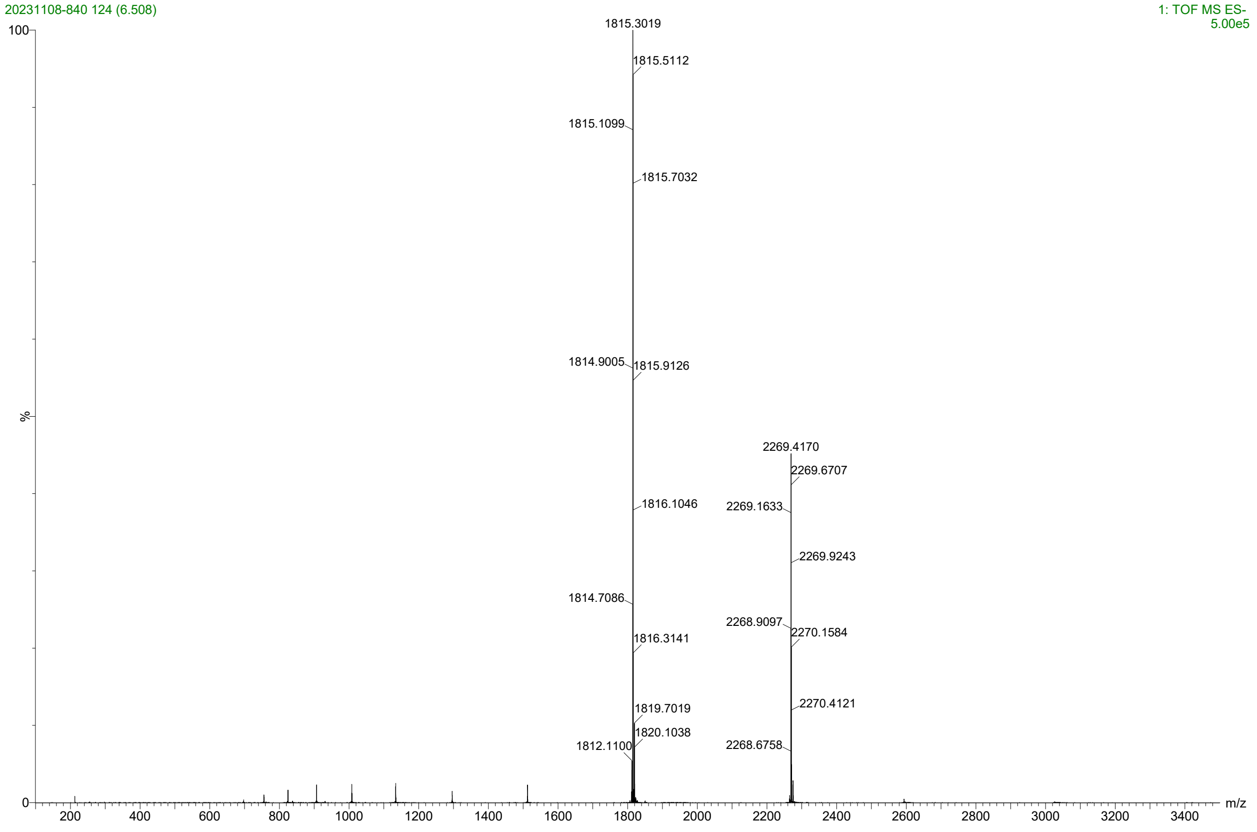
20231108-840 124 (6.508)
1: TOF MS ES-
5.00e5
1815.3019
1815.5112
1815.1099
1815.7032
1814.9005
1815.9126
2269.4170
2269.6707
1816.1046
2269.1633
2269.9243
1814.7086
2268.9097
2270.1584
1816.3141
2270.4121
1819.7019
1820.1038
1812.1100
2268.6758
100
%
0
200
400
600
800
1000
1200
1400
1600
1800
2000
2200
2400
2600
2800
3000
3200
3400
m/z

### Mono-C16(L4+)

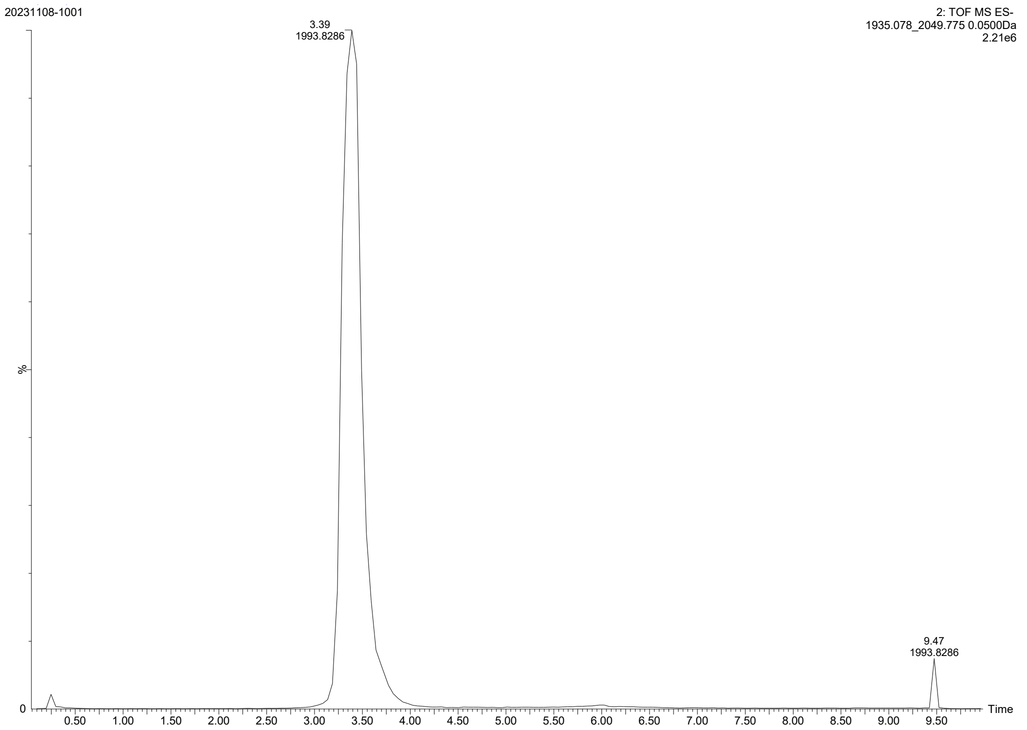
20231108-1001
2: TOF MS ES-
1935.078_2049.775 0.0500Da
2.21e6
3.39
1993.8286
9.47
1993.8286
%
0
0.50
1.00
1.50
2.00
2.50
3.00
3.50
4.00
4.50
5.00
5.50
6.00
6.50
7.00
7.50
8.00
8.50
9.00
9.50
Time

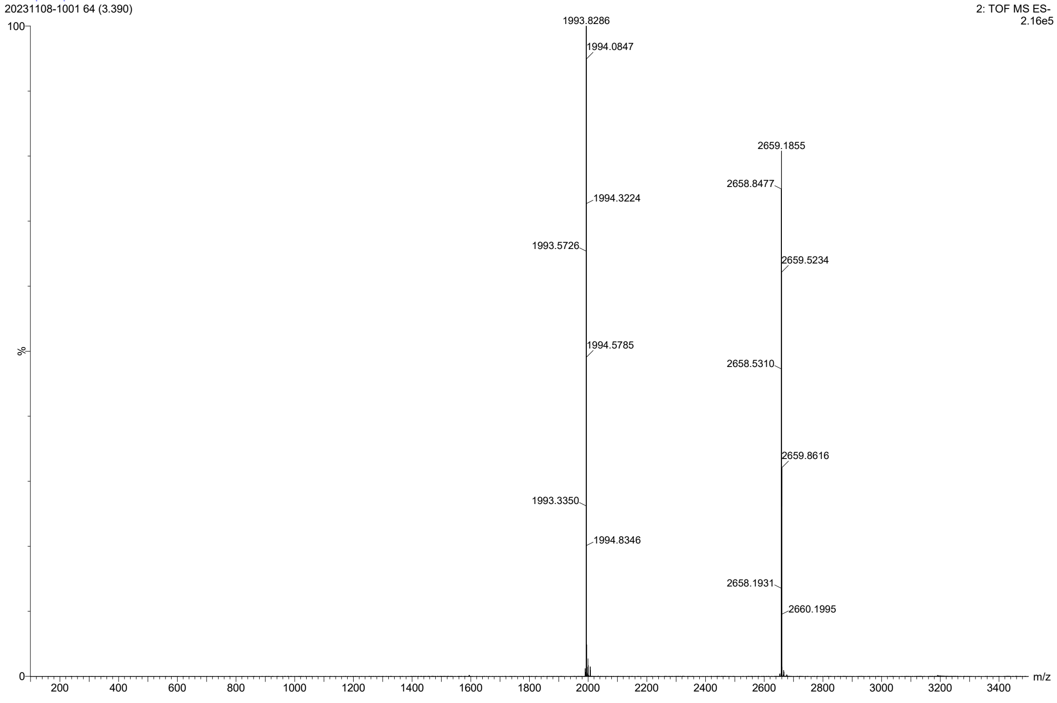
20231108-1001 64 (3.390)
2: TOF MS ES-
2.16e5
100
1993.8286
1994.0847
1994.3224
1993.5726
1994.5785
1993.3350
1994.8346
2659.1855
2658.8477
2659.5234
2658.5310
2659.8616
2658.1931
2660.1995
%
0
200
400
600
800
1000
1200
1400
1600
1800
2000
2200
2400
2600
2800
3000
3200
3400
m/z

Bis-C16(L4+)

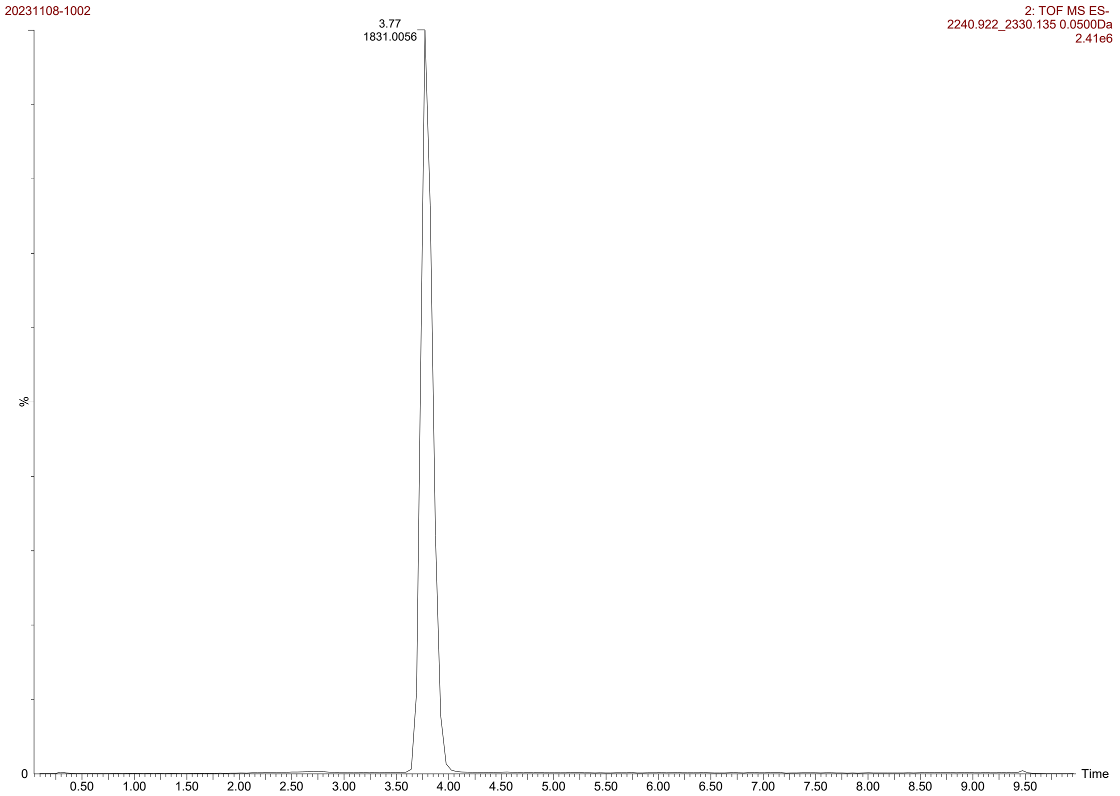
20231108-1002
2: TOF MS ES-
2240.922_2330.135 0.0500Da
2.41e6
3.77
1831.0056
%
0
0.50 1.00 1.50 2.00 2.50 3.00 3.50 4.00 4.50 5.00 5.50 6.00 6.50 7.00 7.50 8.00 8.50 9.00 9.50
Time

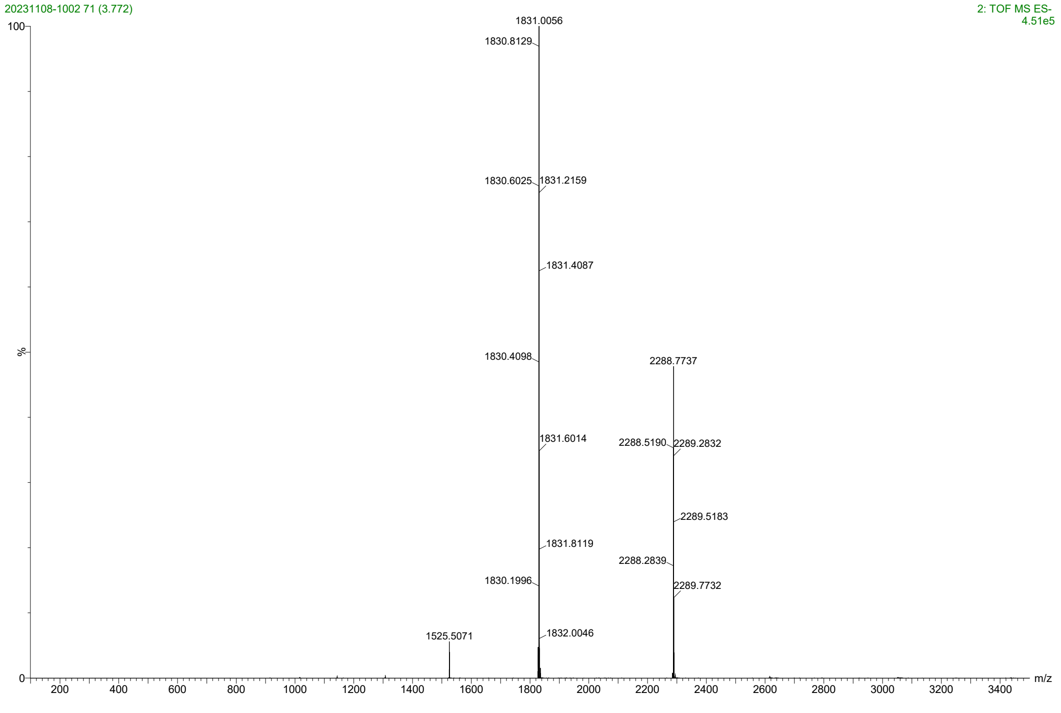
20231108-1002 71 (3.772)
2: TOF MS ES-
4.51e5
100
%
0
1831.0056
1830.8129
1830.6025
1831.2159
1831.4087
1830.4098
2288.7737
1831.6014
2288.5190
2289.2832
2289.5183
1831.8119
2288.2839
1830.1996
2289.7732
1525.5071
1832.0046
200 400 600 800 1000 1200 1400 1600 1800 2000 2200 2400 2600 2800 3000 3200 3400
m/z

Mono-C22(L4+)

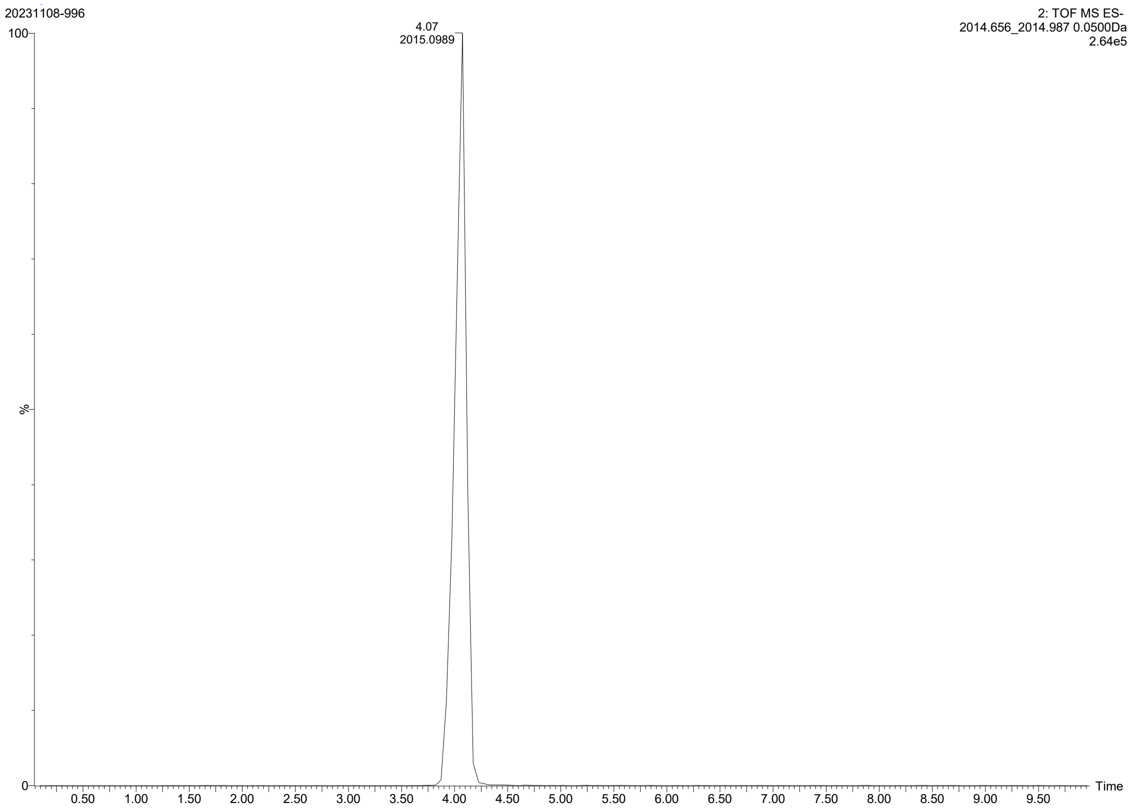

20231108-996
2: TOF MS ES-
2014.656_2014.987 0.0500Da
2.64e5
4.07
2015.0989
100
%
0
Time
0.50
1.00
1.50
2.00
2.50
3.00
3.50
4.00
4.50
5.00
5.50
6.00
6.50
7.00
7.50
8.00
8.50
9.00
9.50


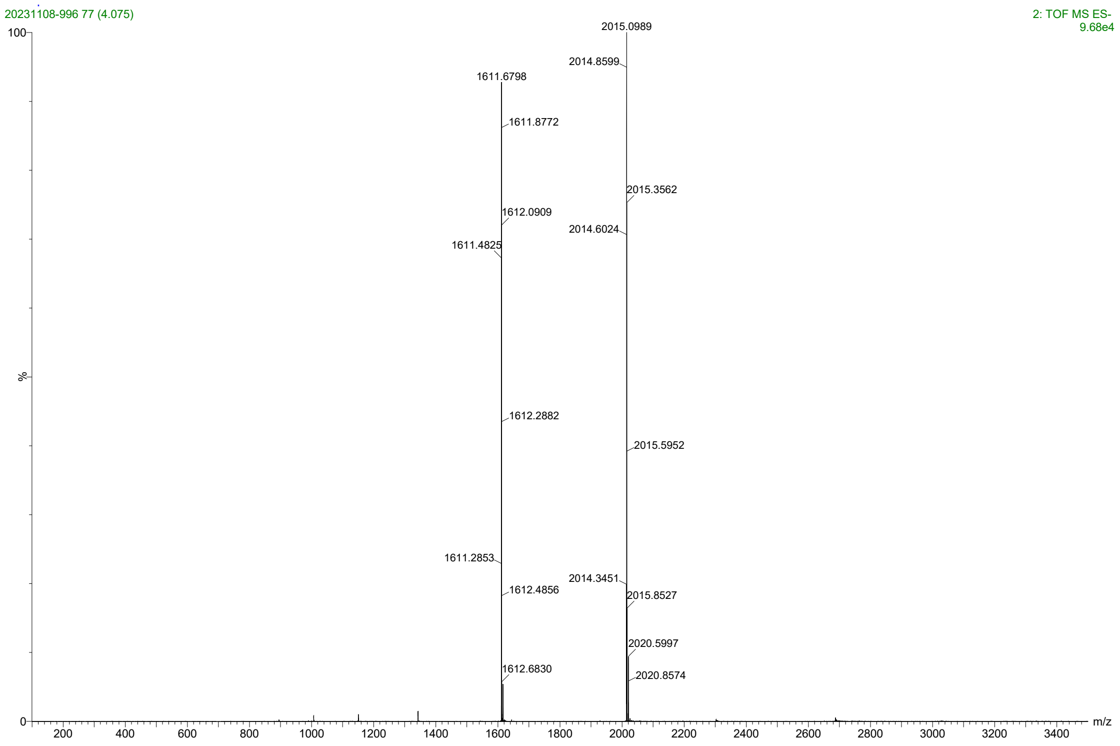

20231108-996 77 (4.075)
2: TOF MS ES-
9.68e4
2015.0989
2014.8599
1611.6798
1611.8772
2015.3562
1612.0909
2014.6024
1611.4825
1612.2882
2015.5952
1611.2853
2014.3451
1612.4856
2015.8527
2020.5997
1612.6830
2020.8574
100
%
0
m/z
200
400
600
800
1000
1200
1400
1600
1800
2000
2200
2400
2600
2800
3000
3200
3400

Bis-C22(L4+)

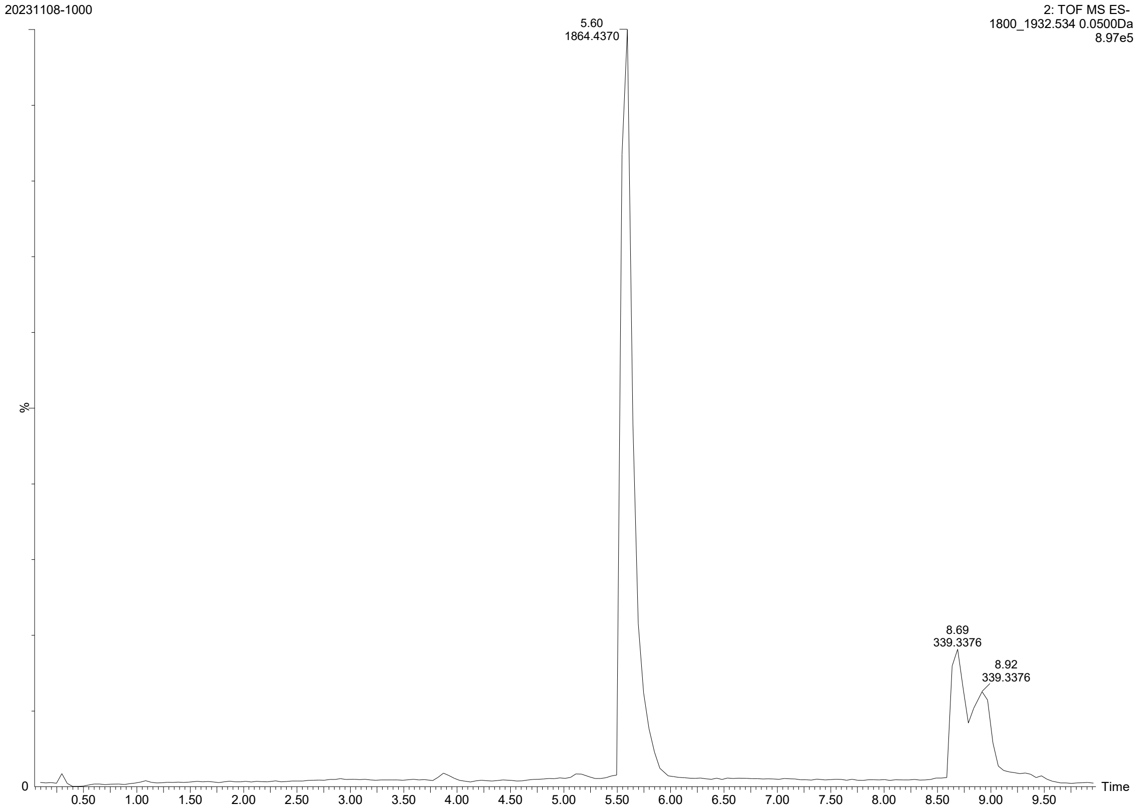

20231108-1000
2: TOF MS ES-
1800_1932.534 0.0500Da
8.97e5
5.60
1864.4370
8.69
339.3376
8.92
339.3376
%
0
0.50
1.00
1.50
2.00
2.50
3.00
3.50
4.00
4.50
5.00
5.50
6.00
6.50
7.00
7.50
8.00
8.50
9.00
9.50
Time


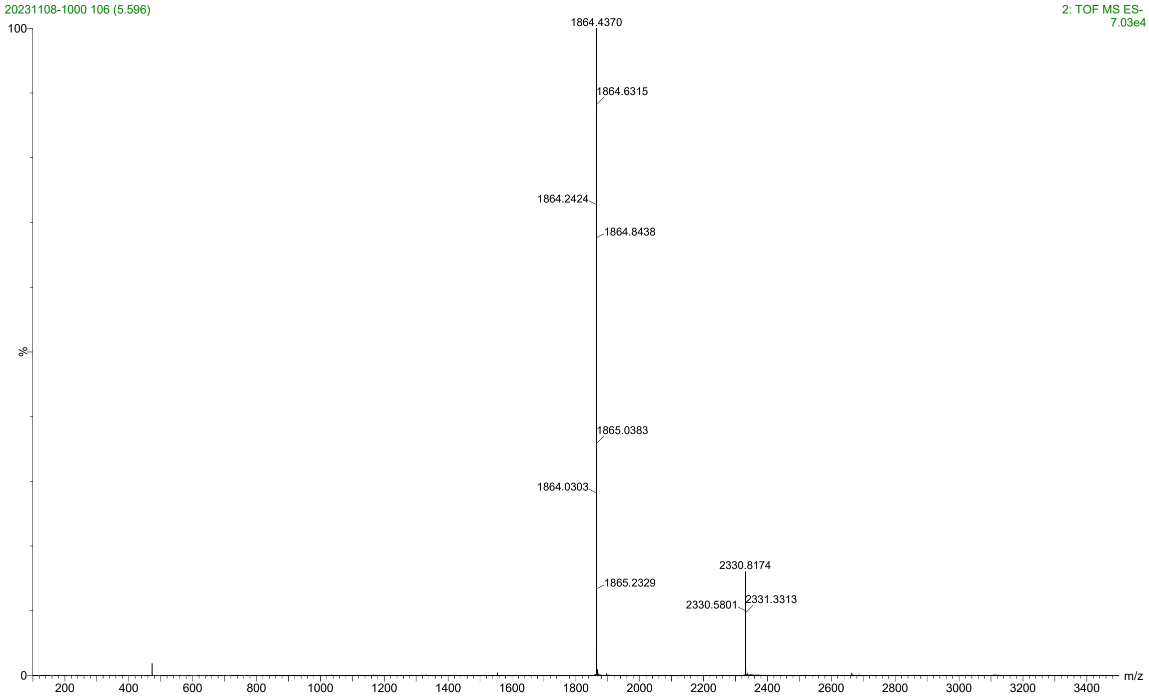

20231108-1000 106 (5.596)
2: TOF MS ES-
7.03e4
100
1864.4370
1864.6315
1864.2424
1864.8438
%
1865.0383
1864.0303
2330.8174
1865.2329
2330.5801
2331.3313
0
200
400
600
800
1000
1200
1400
1600
1800
2000
2200
2400
2600
2800
3000
3200
3400
m/z

Mono-C16-PEG4

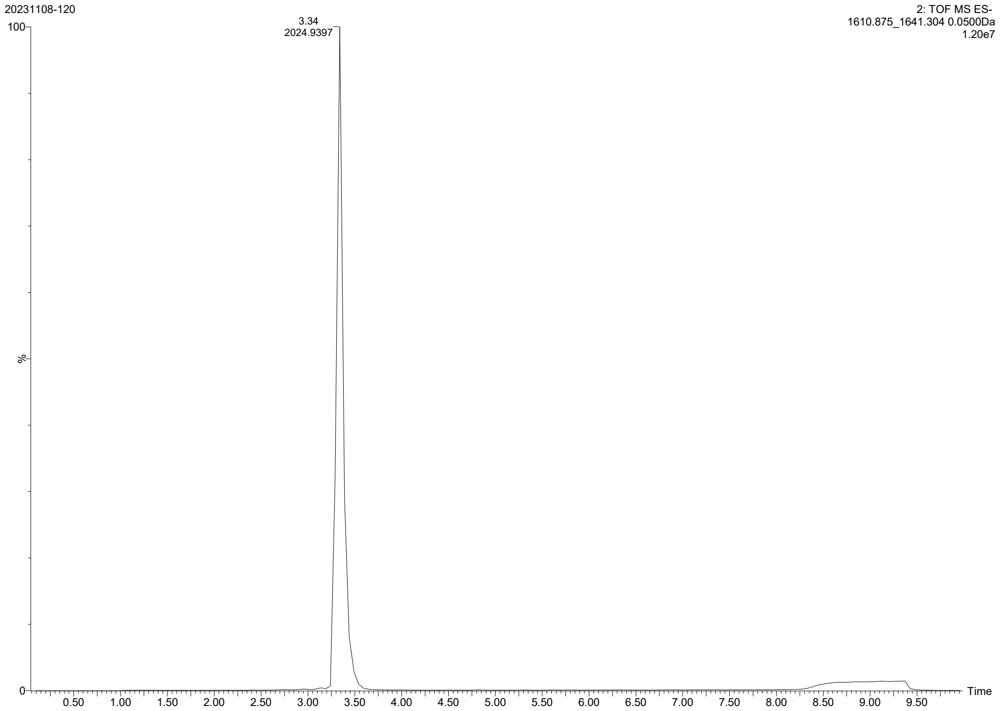


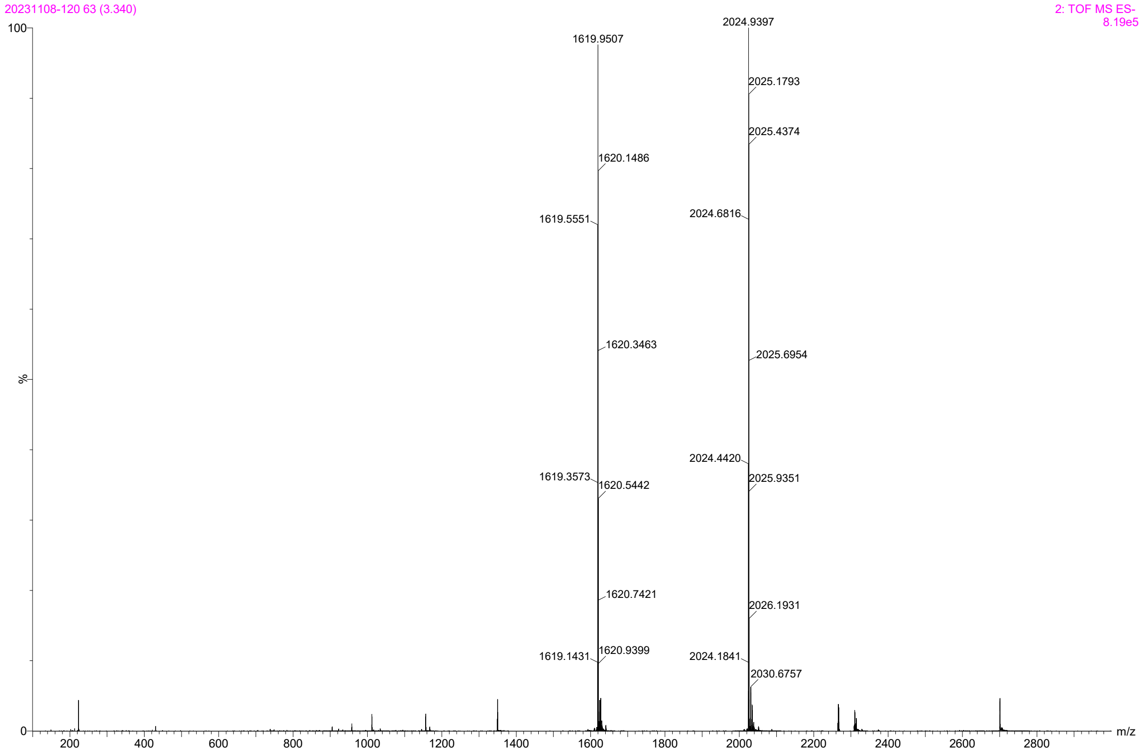

Mono-C16-PEG2000

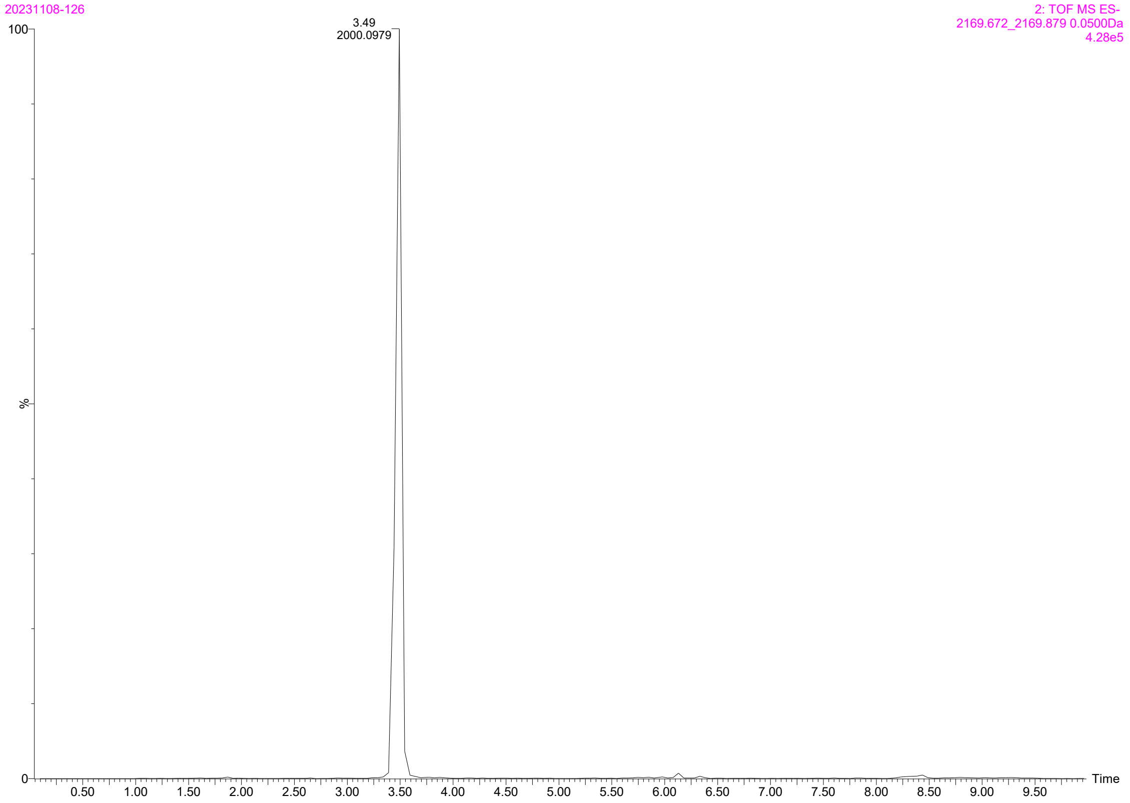


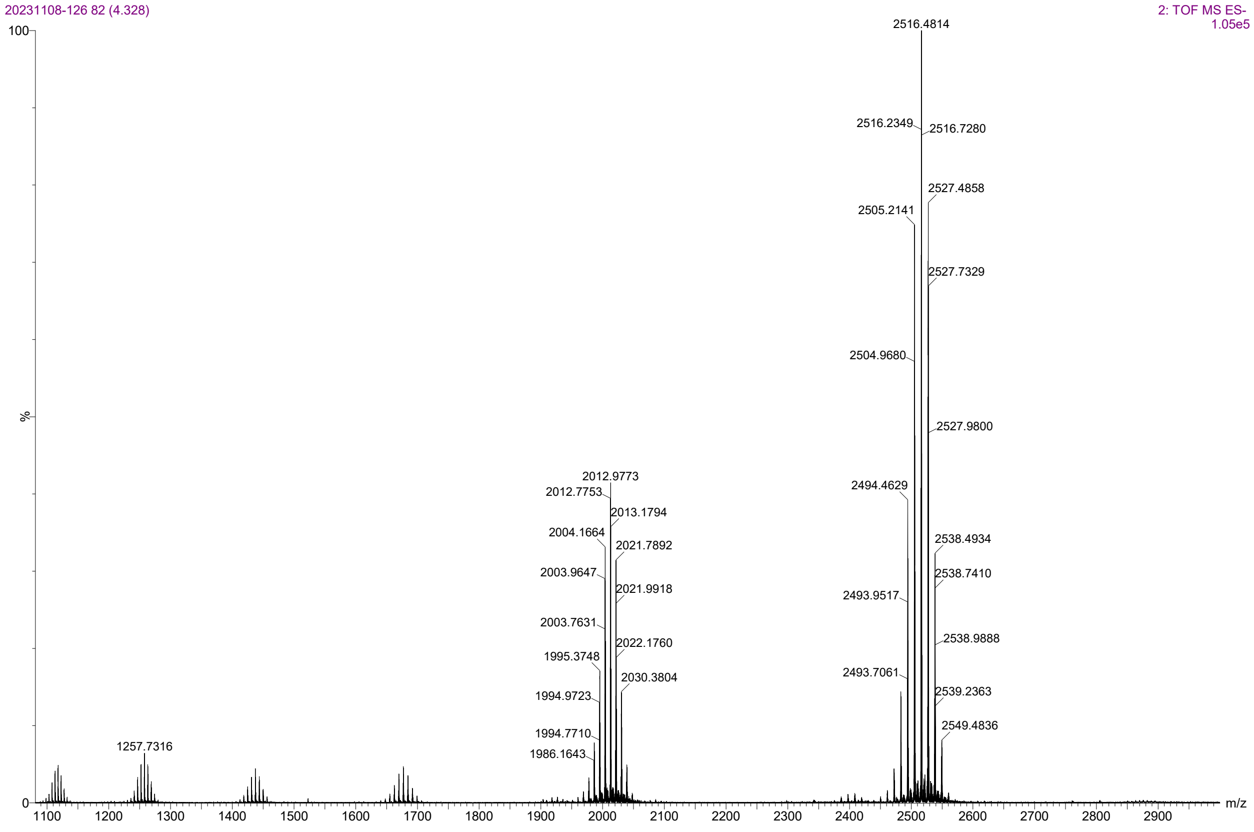

### Bis-C16-PEG4

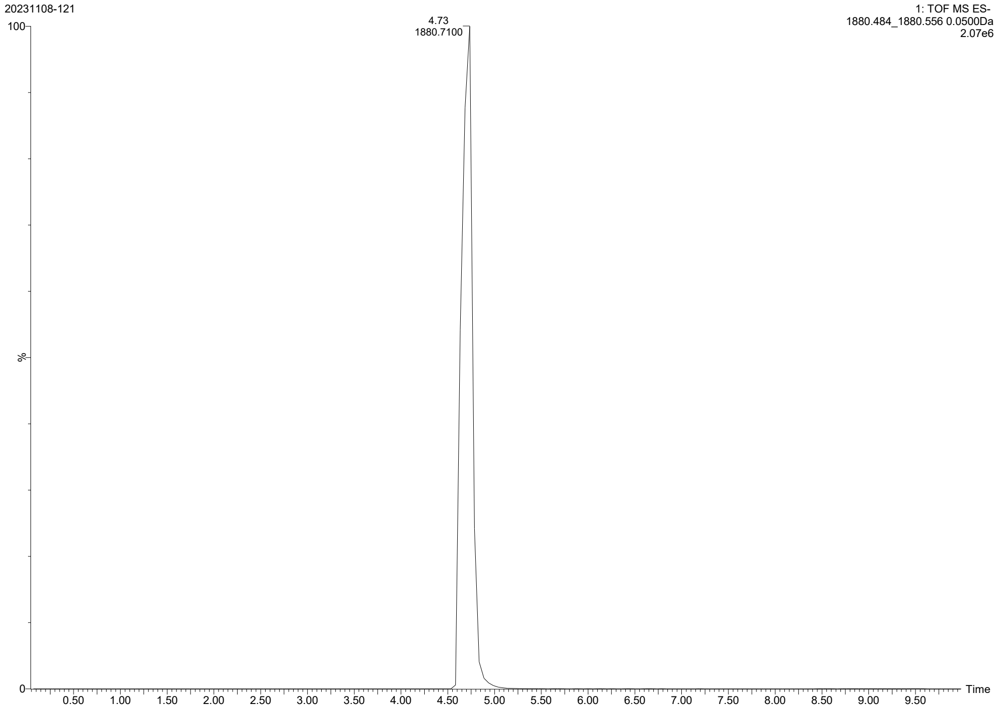
20231108-121
1: TOF MS ES-
1880.484_1880.556 0.0500Da
2.07e6
4.73
1880.7100
100
%
0
0.50
1.00
1.50
2.00
2.50
3.00
3.50
4.00
4.50
5.00
5.50
6.00
6.50
7.00
7.50
8.00
8.50
9.00
9.50
Time

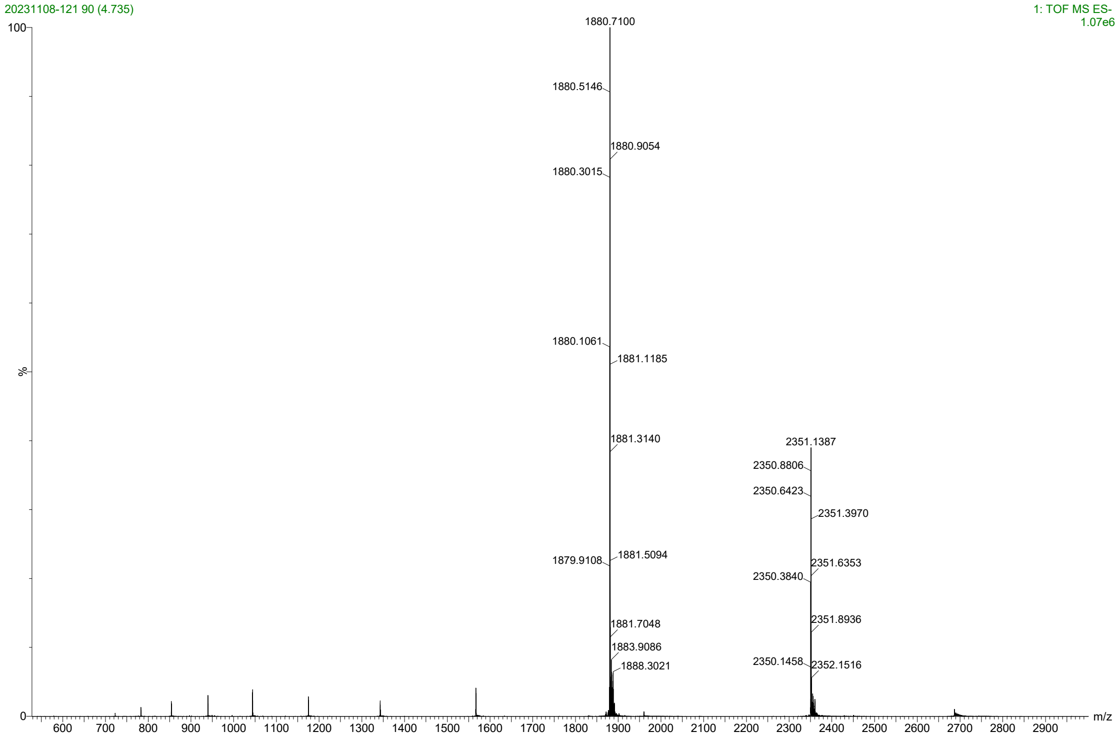
20231108-121 90 (4.735)
1: TOF MS ES-
1.07e6
1880.7100
1880.5146
1880.9054
1880.3015
1880.1061
1881.1185
1881.3140
2351.1387
2350.8806
2350.6423
2351.3970
1879.9108
1881.5094
2351.6353
2350.3840
1881.7048
2351.8936
1883.9086
2350.1458
2352.1516
1888.3021
100
%
0
600
700
800
900
1000
1100
1200
1300
1400
1500
1600
1700
1800
1900
2000
2100
2200
2300
2400
2500
2600
2700
2800
2900
m/z

### Branched C16-PEG4

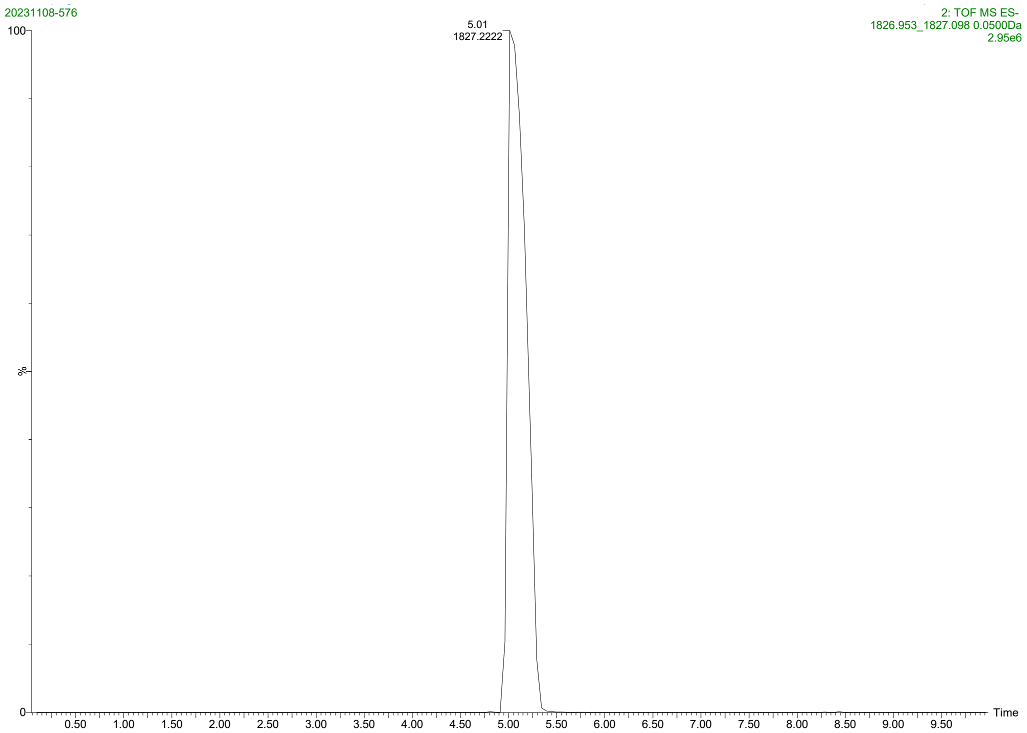


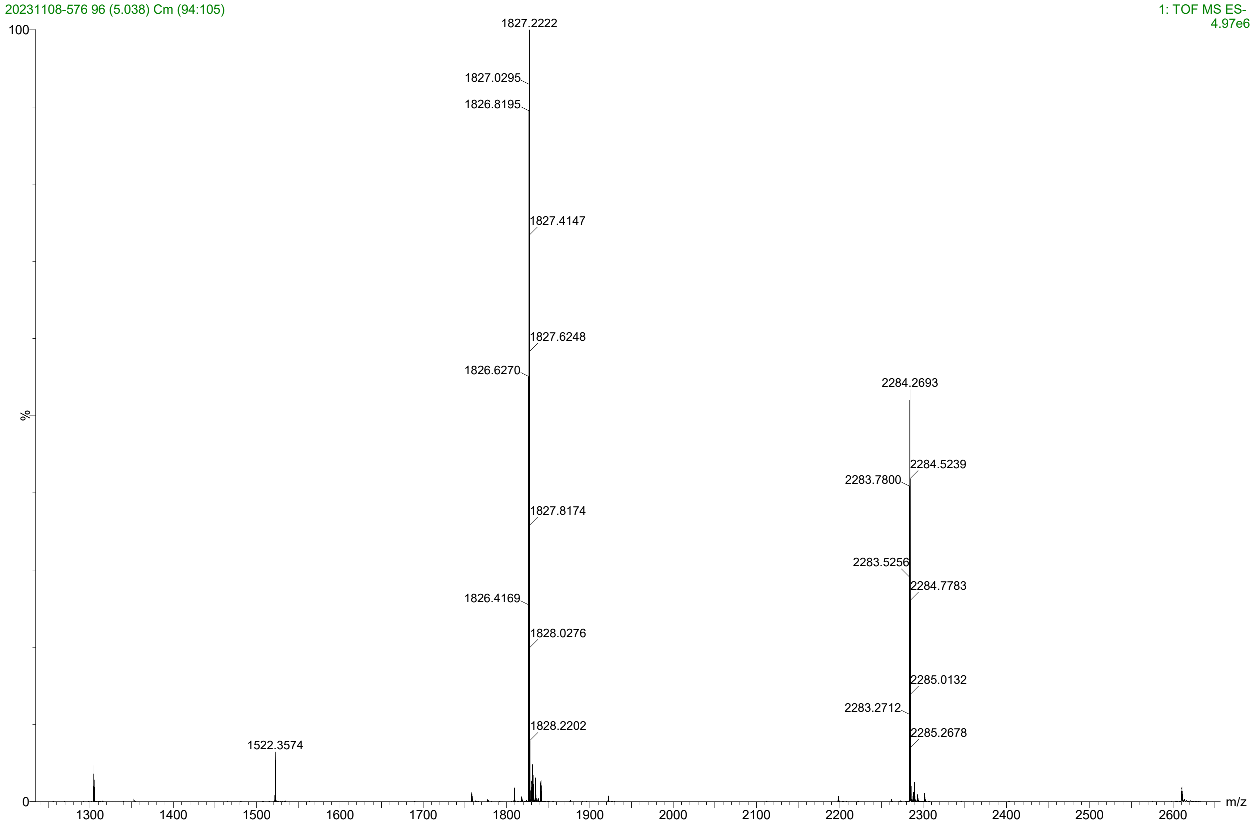

#### Branched C16

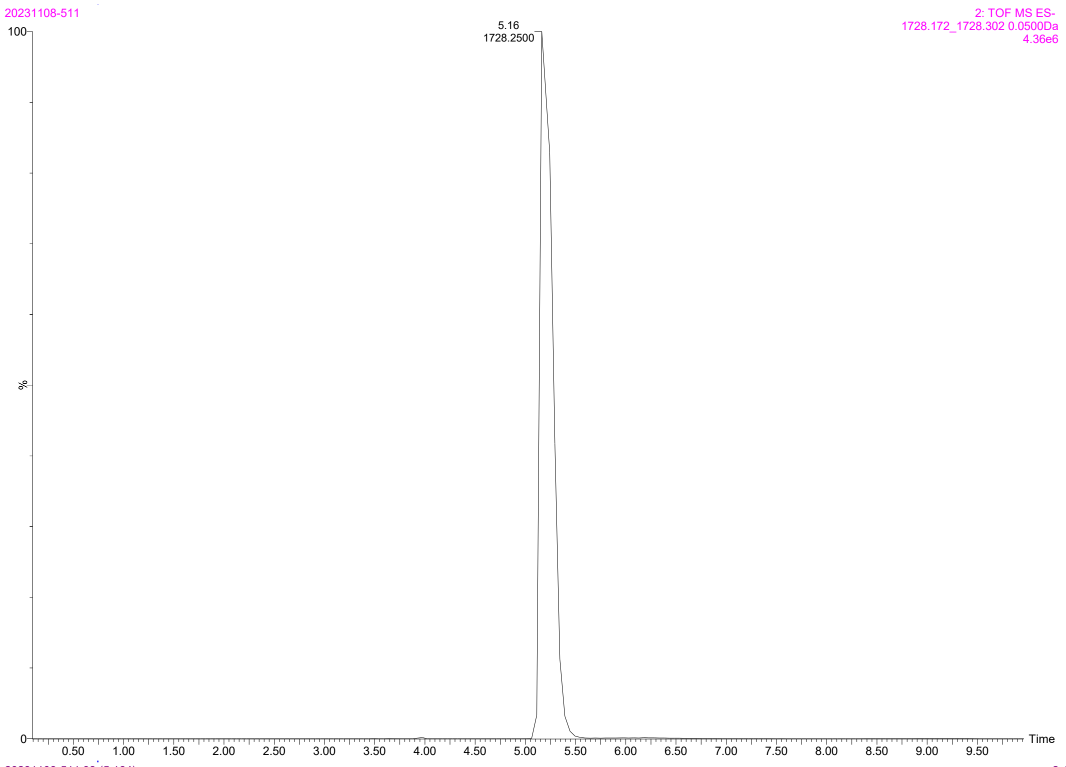


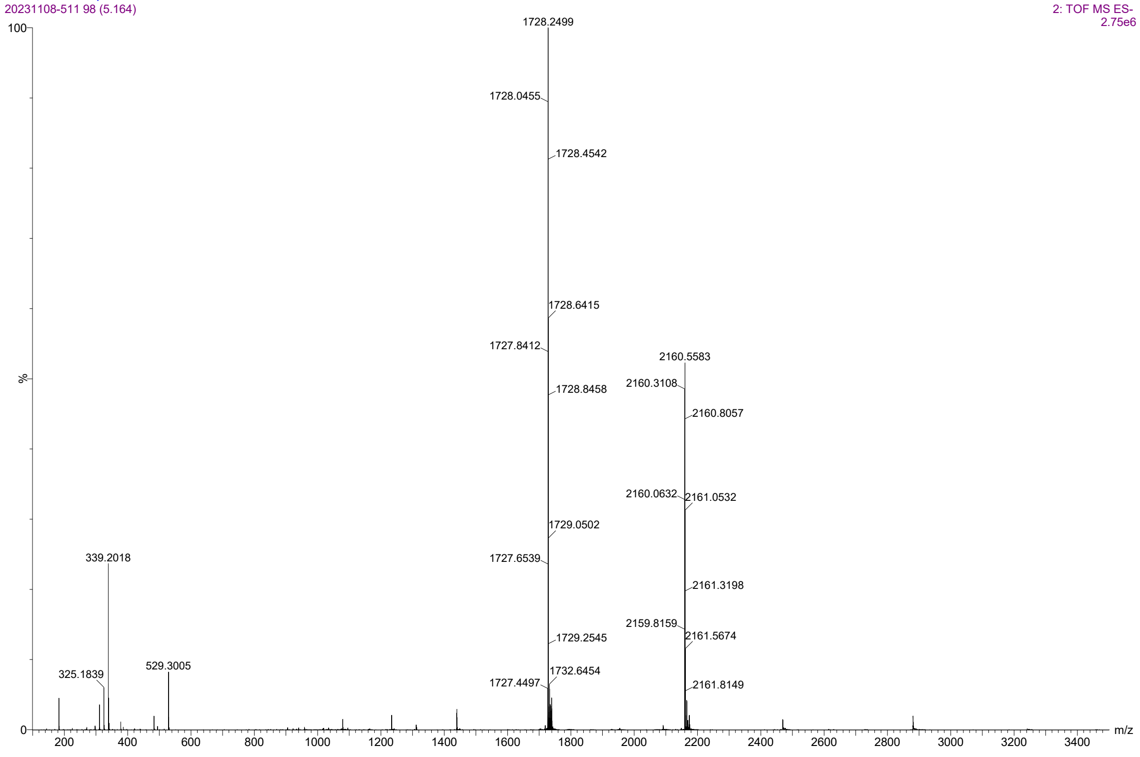

Bis-branched C16-PEG4

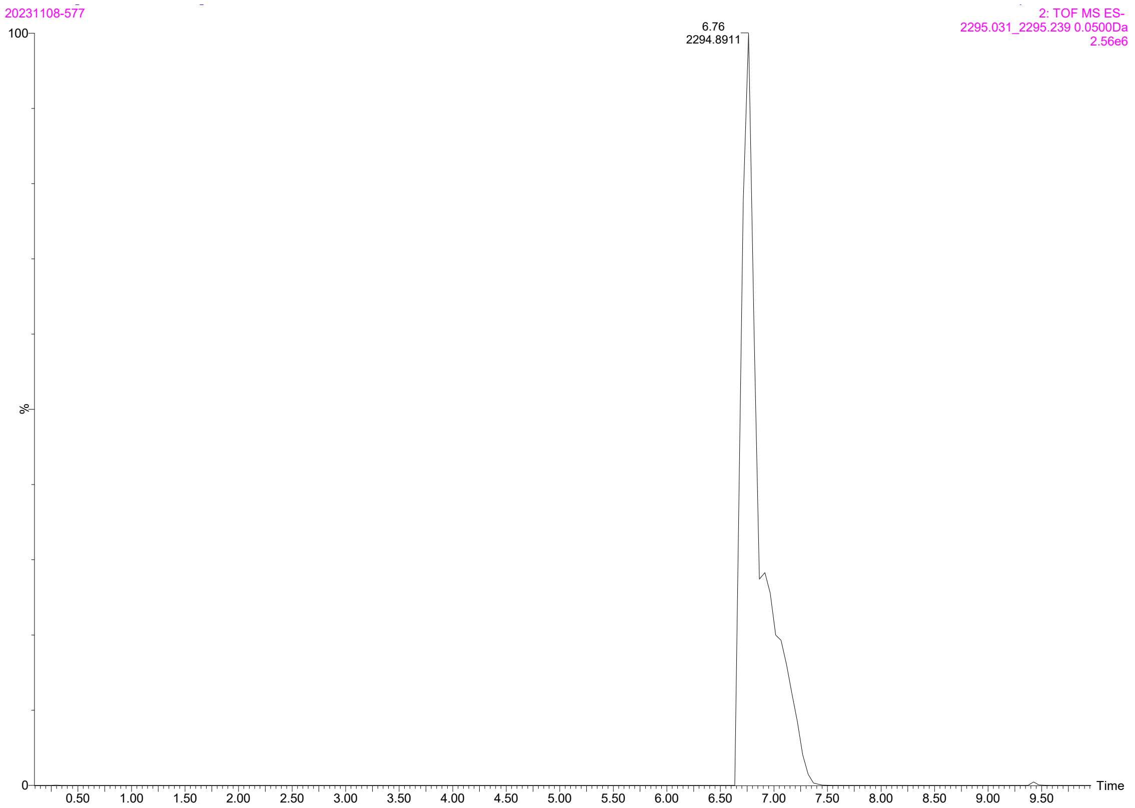


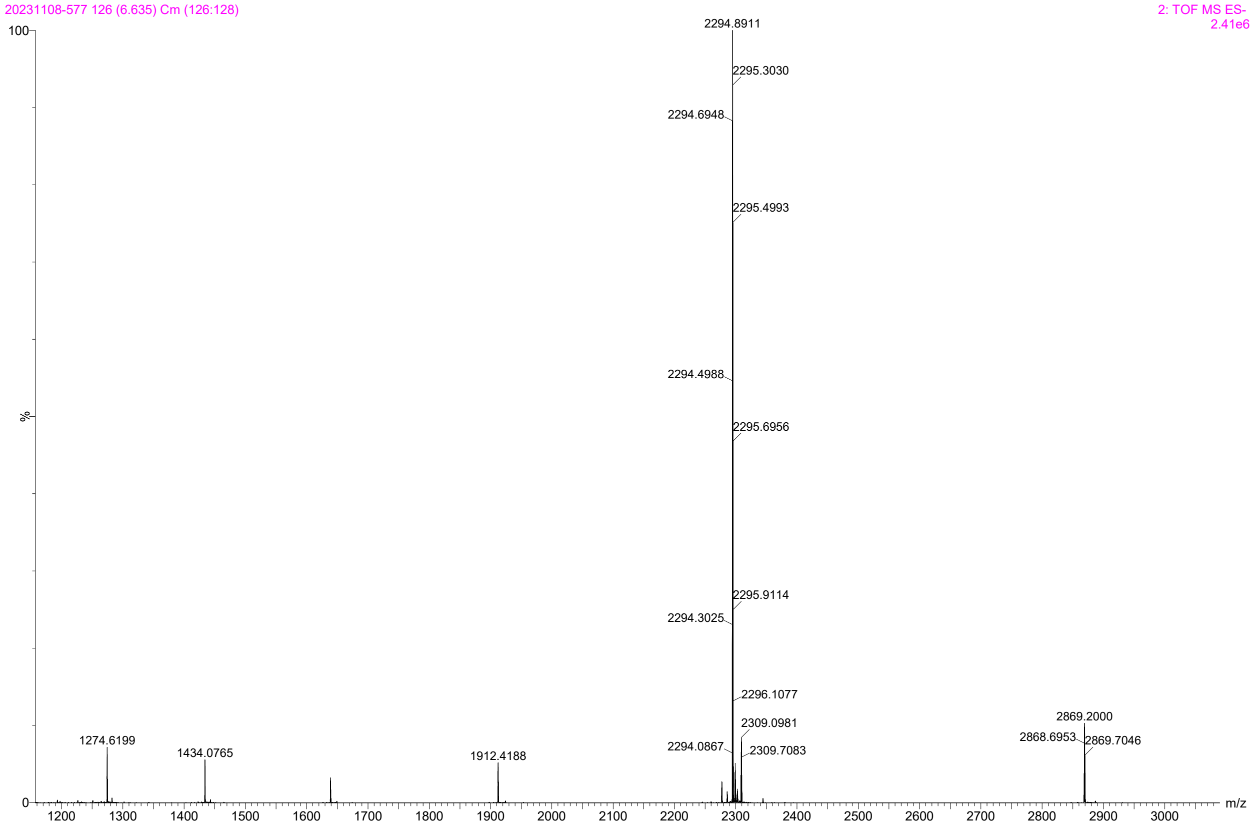

Mono-DOPE-PEG2000

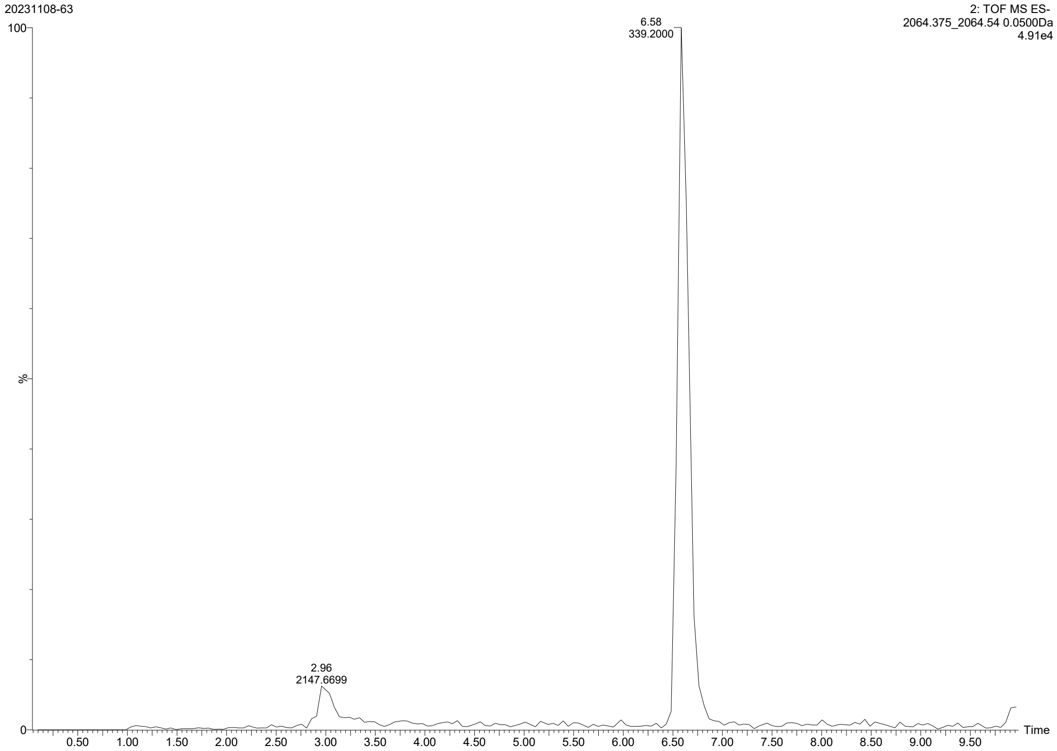


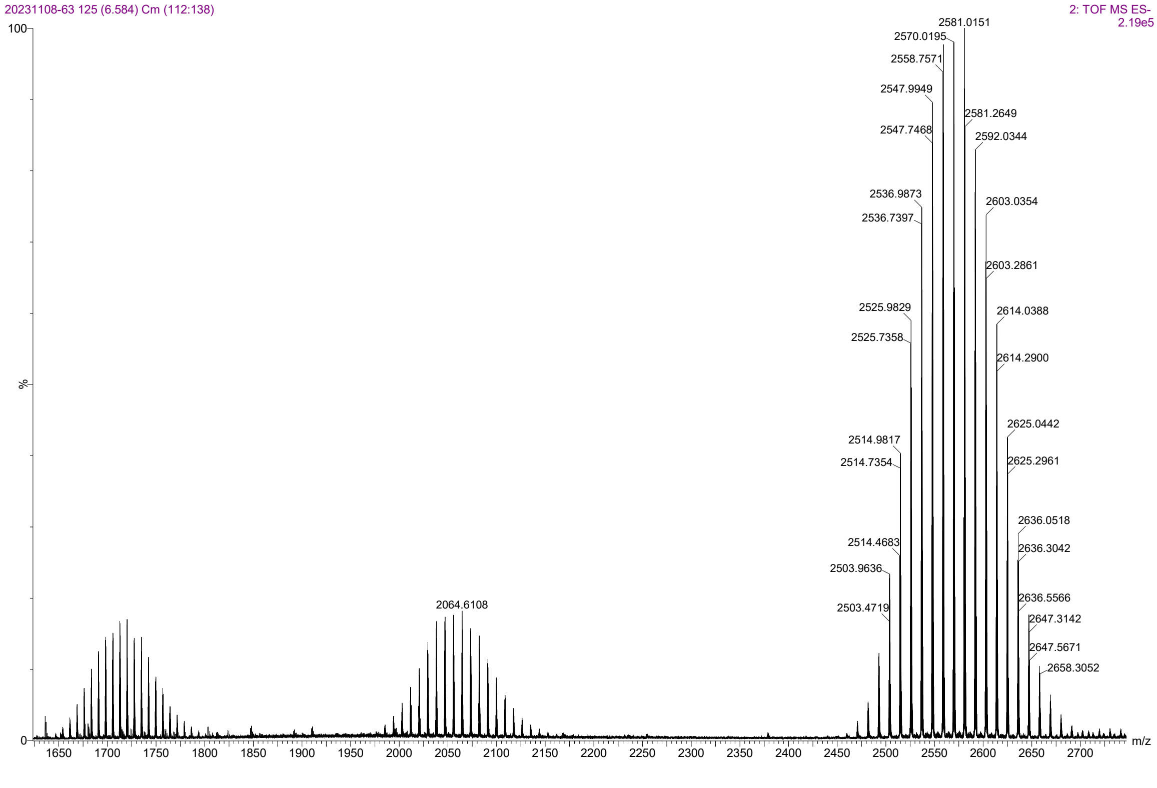

### Mono-CTP

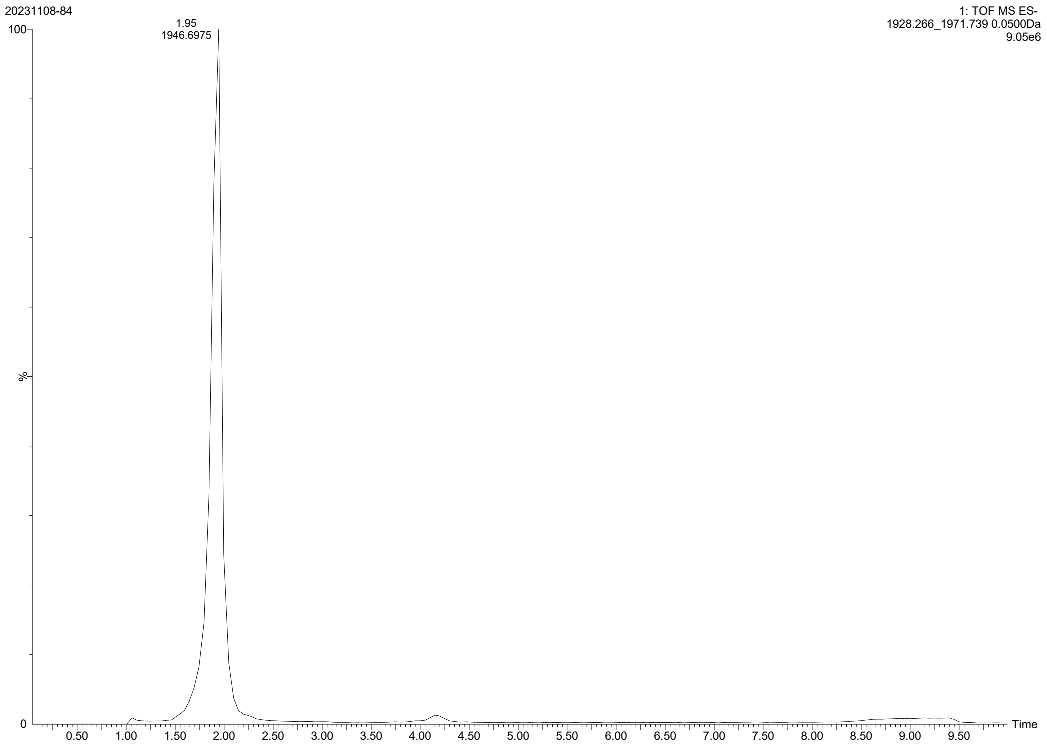
20231108-84
1: TOF MS ES-
1928.266_1971.739 0.0500Da
9.05e6
1.95
1946.6975
Time
%

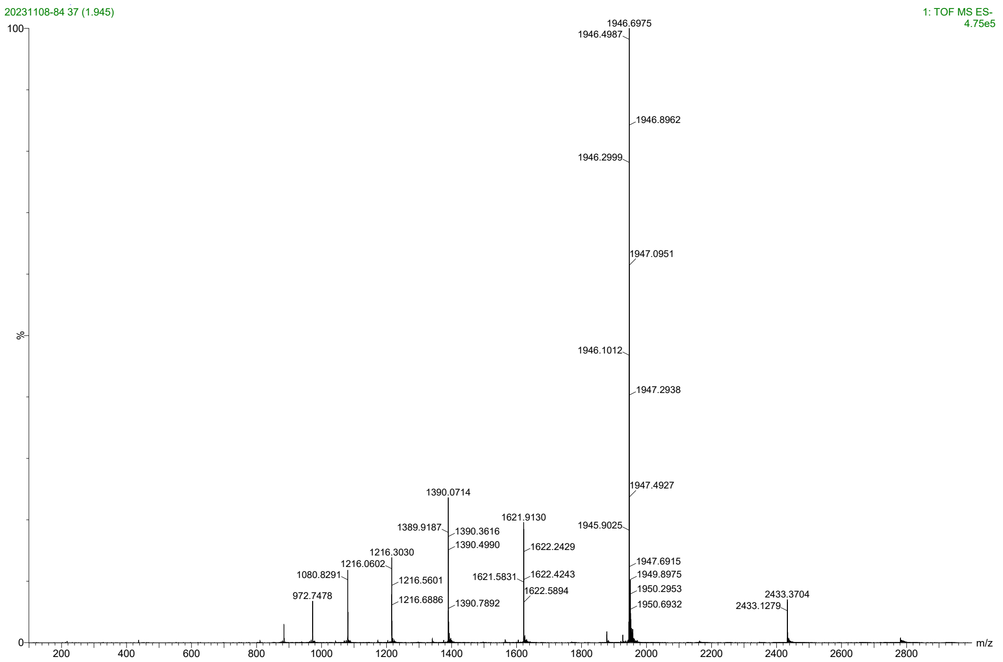
20231108-84 37 (1.945)
1: TOF MS ES-
4.75e5
1946.6975
1946.4987
1946.8962
1946.2999
1947.0951
1946.1012
1947.2938
1947.4927
1945.9025
1947.6915
1949.8975
1950.2953
1950.6932
1390.0714
1389.9187
1390.3616
1390.4990
1390.7892
1621.9130
1622.2429
1621.5831
1622.4243
1622.5894
1216.3030
1216.0602
1216.5601
1216.6886
1080.8291
972.7478
2433.3704
2433.1279
m/z
%

Bis-CTP

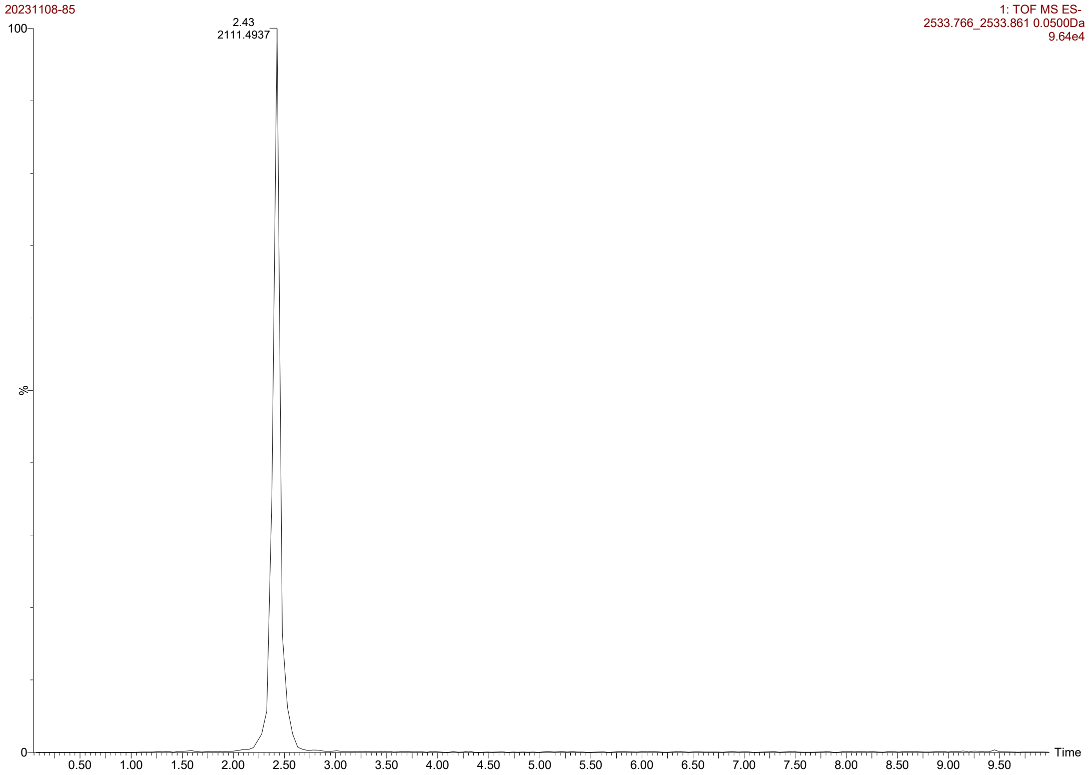
20231108-85
1: TOF MS ES-
2533.766_2533.861 0.0500Da
9.64e4
2.43
2111.4937
100
%
0
Time
0.50
1.00
1.50
2.00
2.50
3.00
3.50
4.00
4.50
5.00
5.50
6.00
6.50
7.00
7.50
8.00
8.50
9.00
9.50

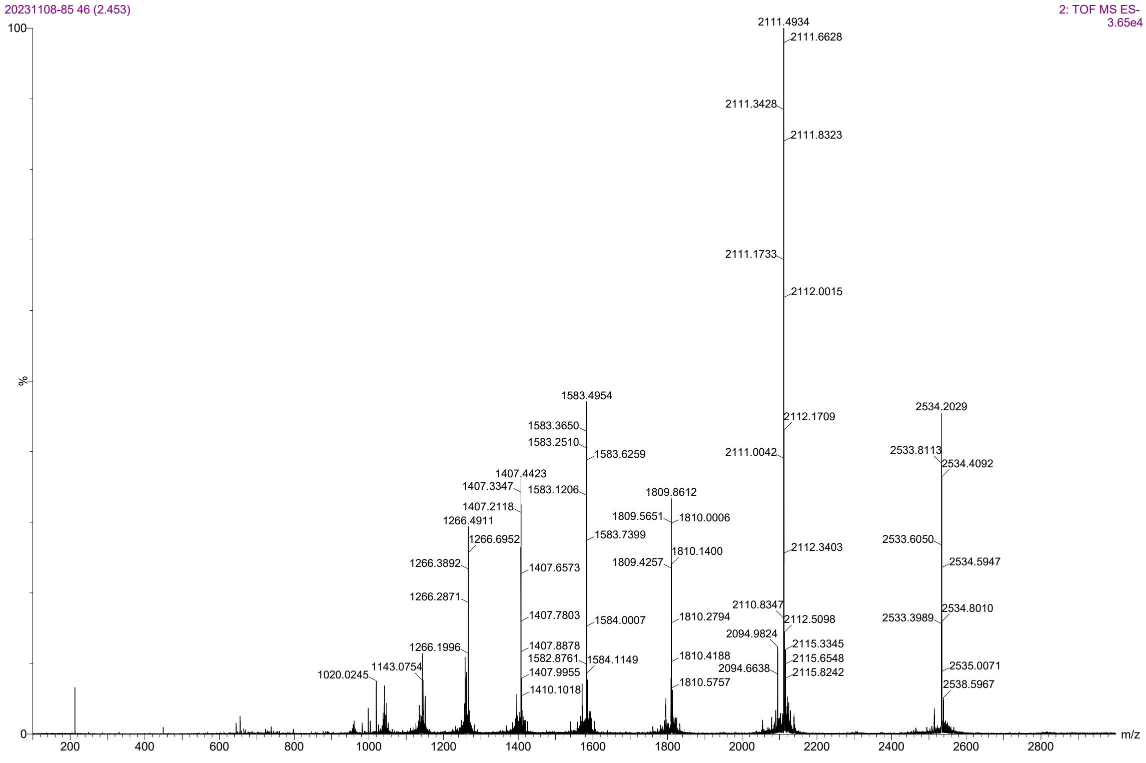
20231108-85 46 (2.453)
2: TOF MS ES-
3.65e4
2111.4934
2111.6628
2111.3428
2111.8323
2111.1733
2112.0015
1583.4954
2534.2029
2112.1709
1583.3650
1583.2510
1583.6259
2111.0042
2533.8113
2534.4092
1407.4423
1407.3347
1583.1206
1809.8612
1407.2118
1809.5651
1810.0006
1266.4911
1583.7399
1266.6952
2533.6050
1810.1400
2112.3403
1266.3892
1809.4257
1407.6573
2534.5947
1266.2871
2110.8347
2534.8010
1407.7803
1584.0007
1810.2794
2112.5098
2533.3989
2094.9824
1266.1996
1407.8878
2115.3345
1582.8761
1810.4188
1143.0754
1584.1149
2115.6548
1020.0245
1407.9955
2094.6638
2115.8242
2535.0071
1410.1018
1810.5757
2538.5967
100
%
0
m/z
200
400
600
800
1000
1200
1400
1600
1800
2000
2200
2400
2600
2800

C16-PEG4/CTP (5'/3')

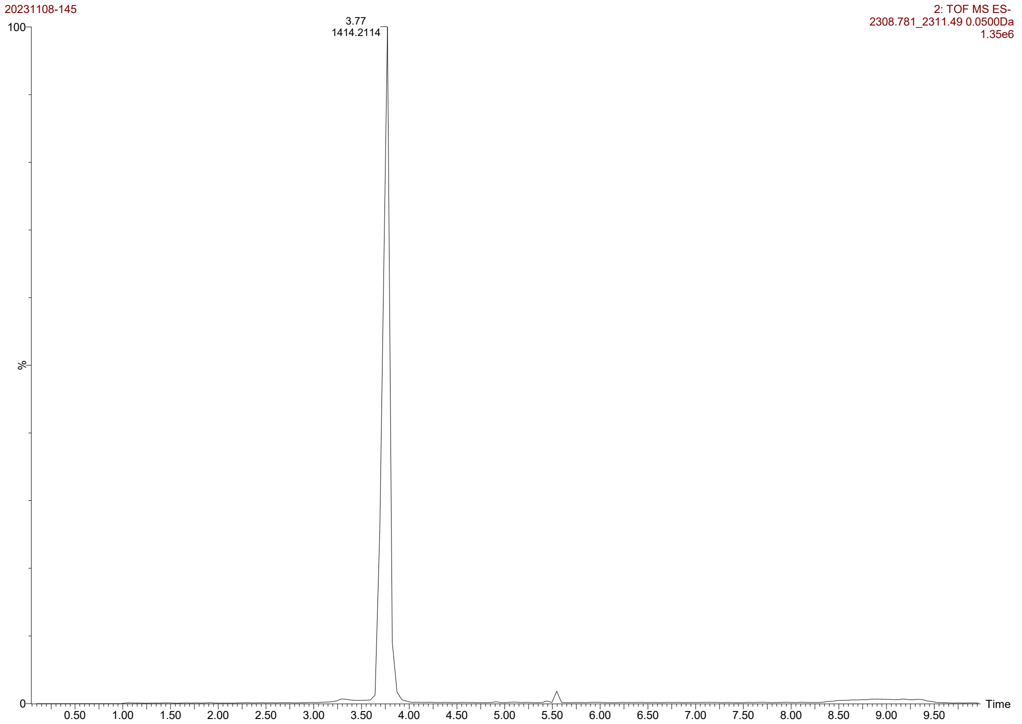


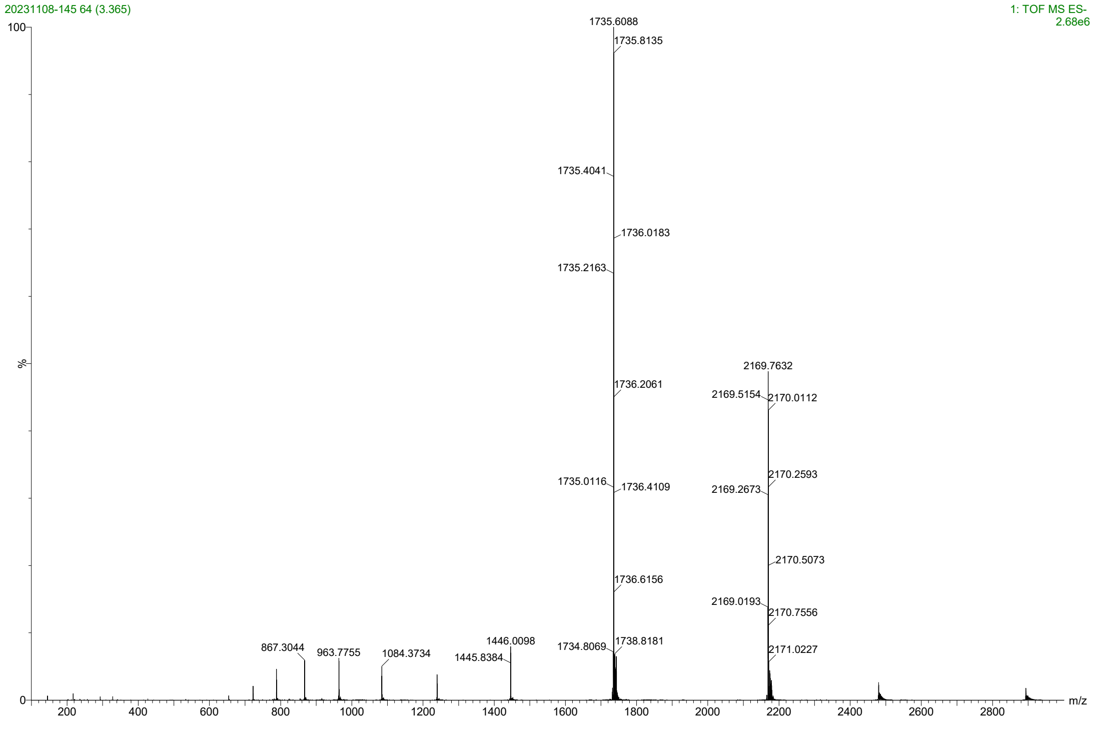

### C16-PEG4/GWWG (5'/3')

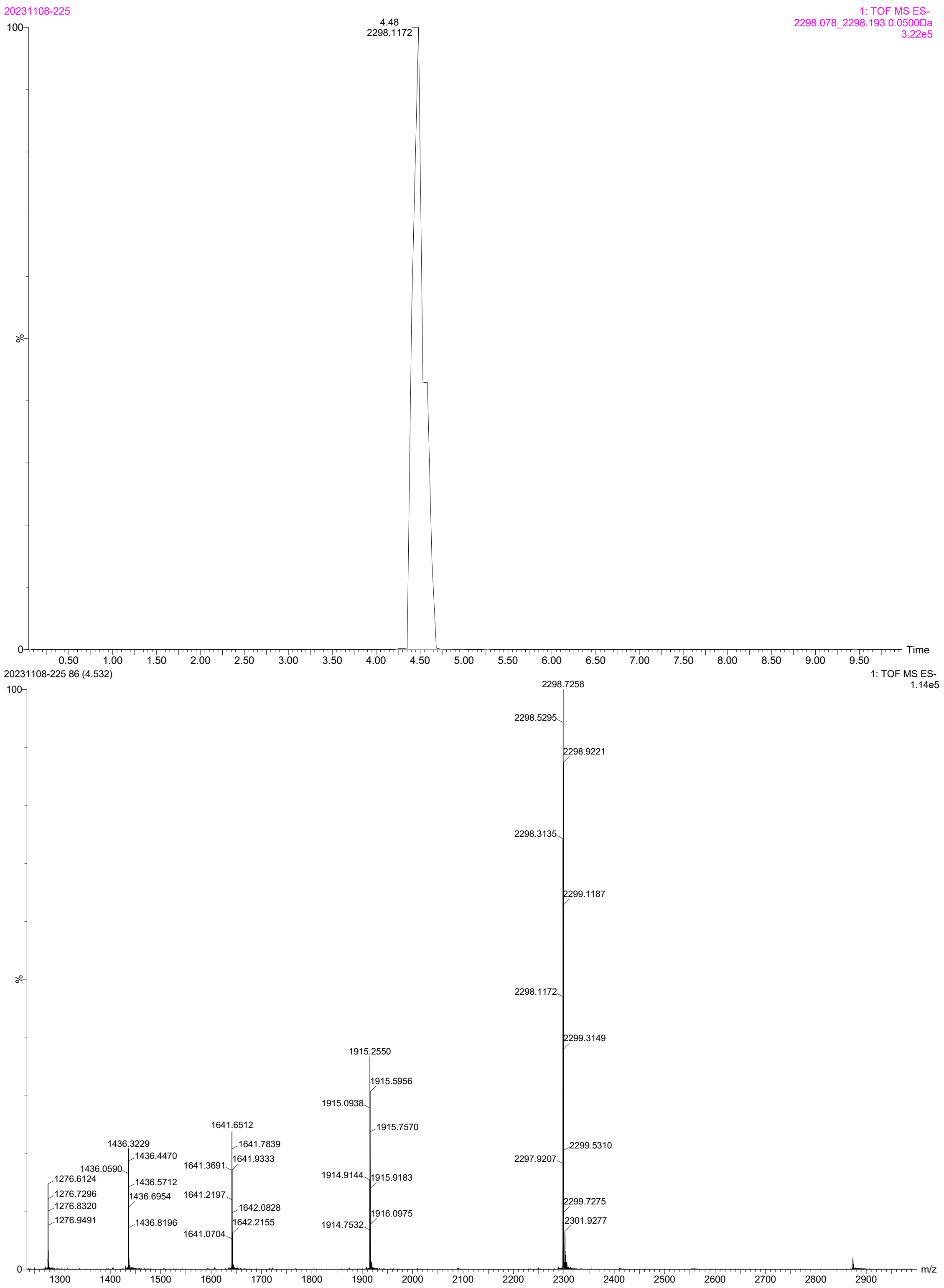

CTP/GWWG (5'/3')

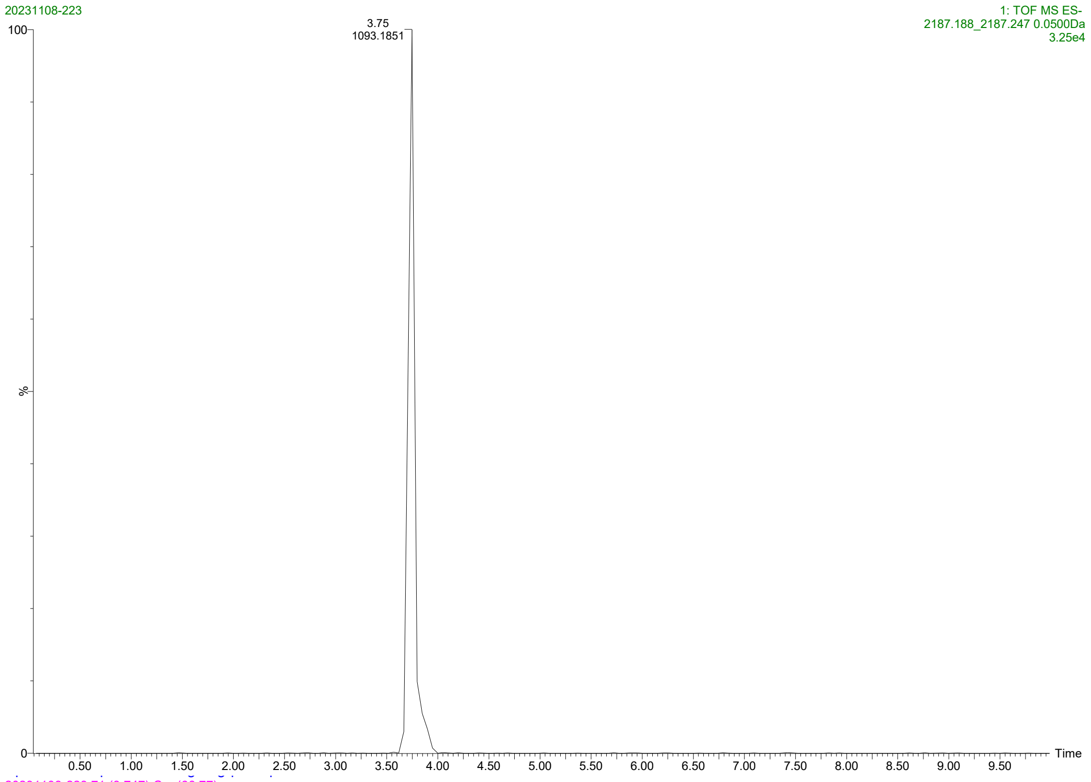
20231108-223
1: TOF MS ES-
2187.188_2187.247 0.0500Da
3.25e4
3.75
1093.1851
100
%
0
Time
0.50
1.00
1.50
2.00
2.50
3.00
3.50
4.00
4.50
5.00
5.50
6.00
6.50
7.00
7.50
8.00
8.50
9.00
9.50

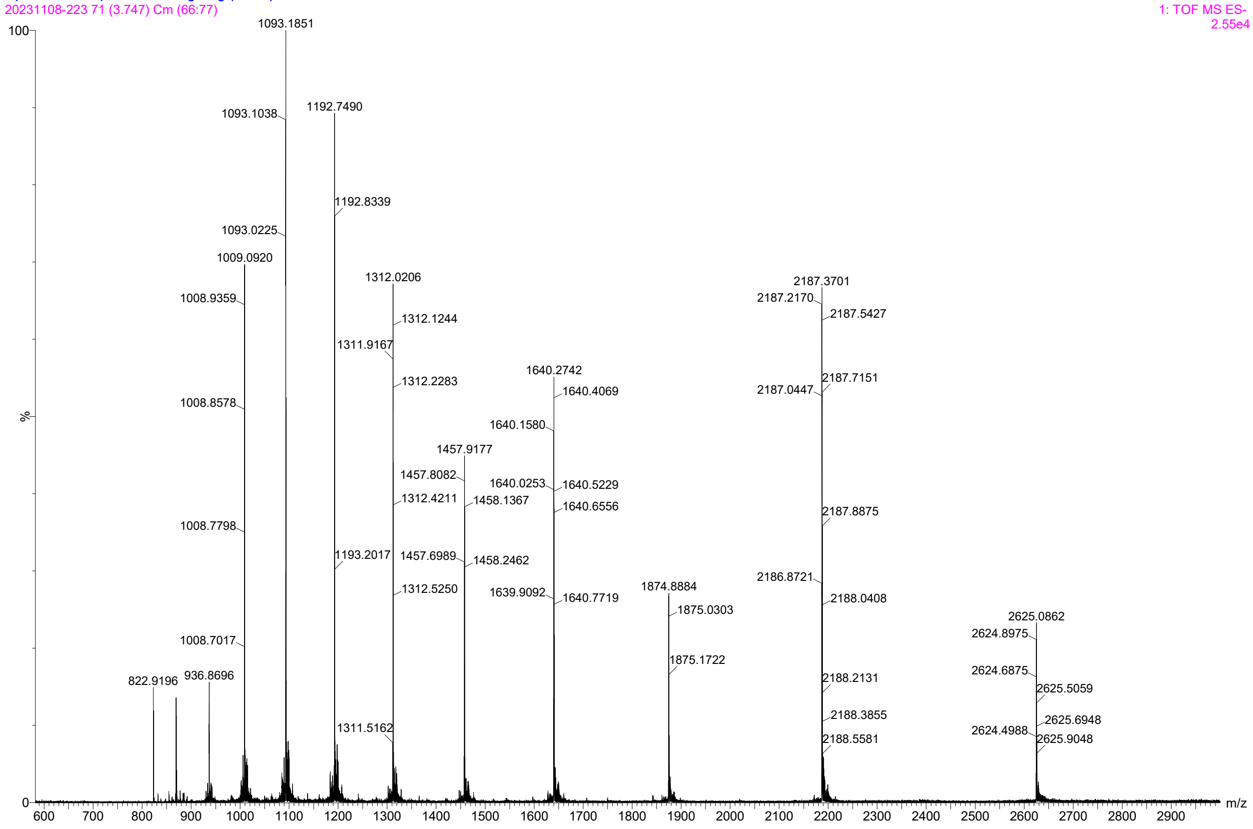
20231108-223 71 (3.747) Cm (66:77)
1: TOF MS ES-
2.55e4
1093.1851
1093.1038
1192.7490
1192.8339
1093.0225
1009.0920
1312.0206
2187.3701
1008.9359
2187.2170
2187.5427
1312.1244
1311.9167
1640.2742
1312.2283
1640.4069
2187.0447
2187.7151
1008.8578
1640.1580
1457.9177
1457.8082
1640.0253
1640.5229
1312.4211
1458.1367
1640.6556
1008.7798
2187.8875
1193.2017
1457.6989
1458.2462
2186.8721
1312.5250
1874.8884
1639.9092
1640.7719
2188.0408
1875.0303
2625.0862
2624.8975
1008.7017
1875.1722
2624.6875
822.9196
936.8696
2188.2131
2625.5059
2188.3855
2625.6948
1311.5162
2624.4988
2188.5581
2625.9048
100
%
0
m/z
600
700
800
900
1000
1100
1200
1300
1400
1500
1600
1700
1800
1900
2000
2100
2200
2300
2400
2500
2600
2700
2800
2900

<u>ASOs</u>

Naked

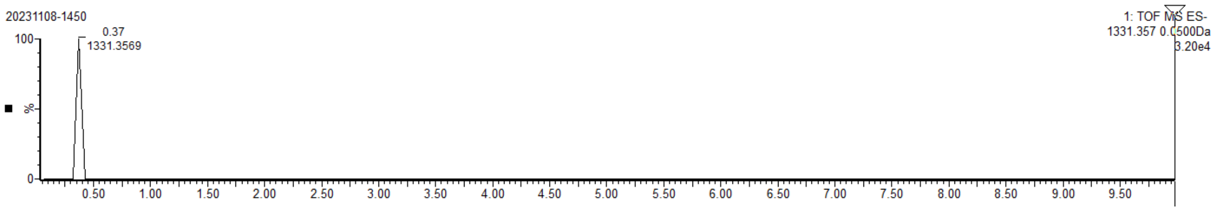


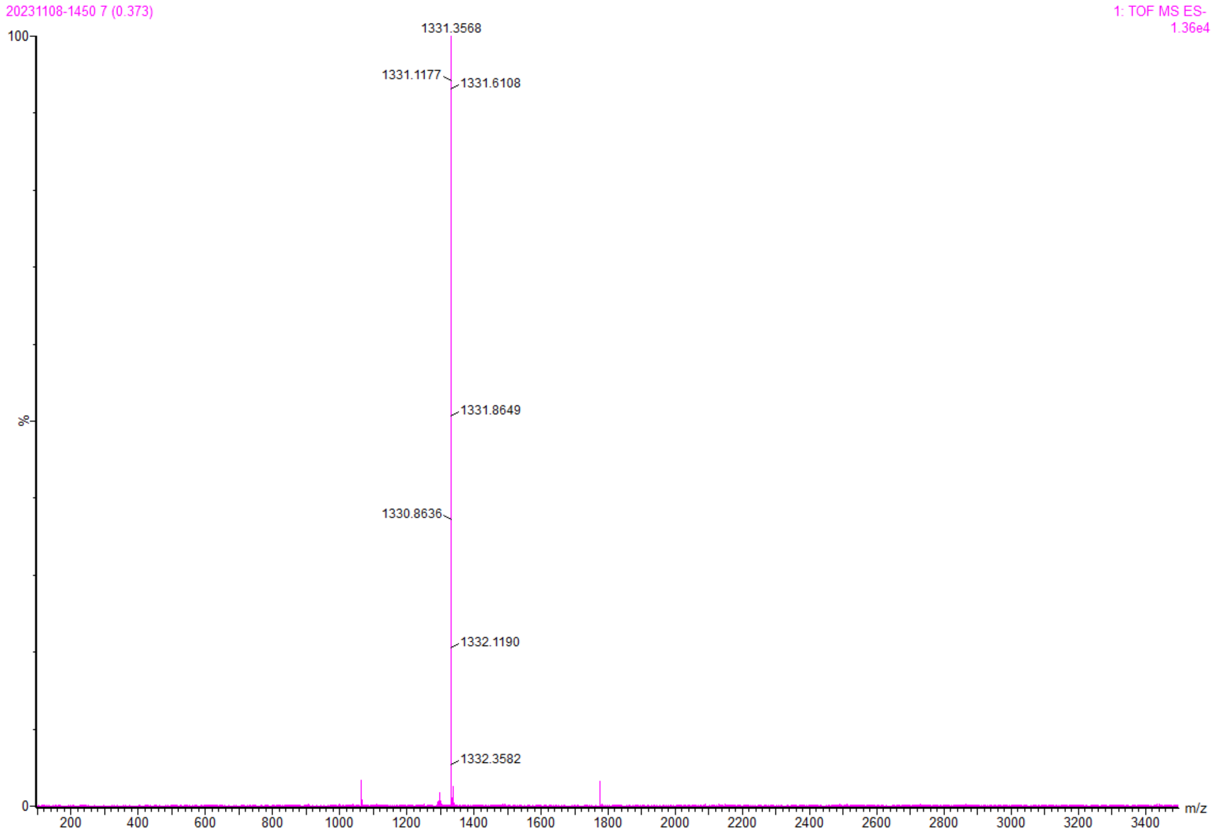

Mono-C16

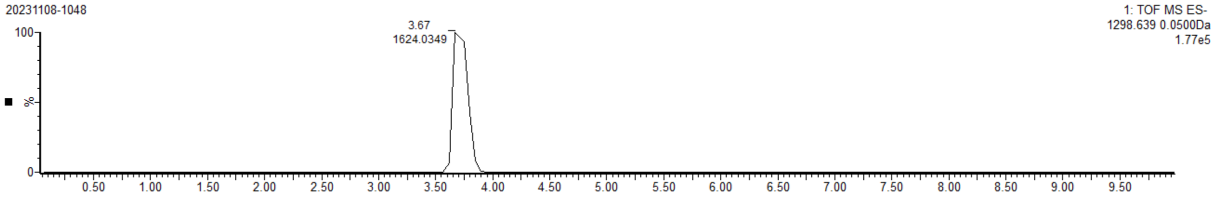

20231108-1048
1: TOF MS ES-
1298.639 0.0500Da
1.77e5
3.67
1624.0349
%


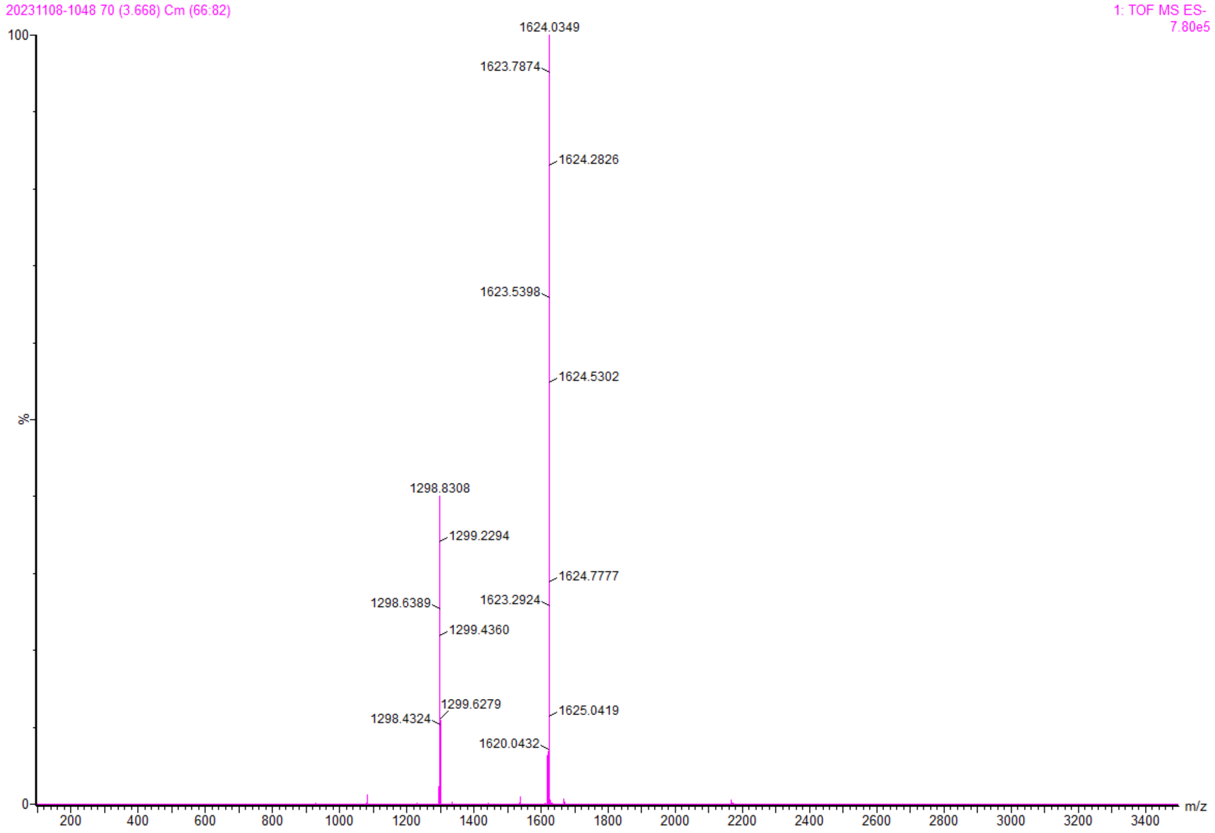

20231108-1048 70 (3.668) Cm (66:82)
1: TOF MS ES-
7.80e5
1624.0349
1623.7874
1624.2826
1623.5398
1624.5302
1298.8308
1299.2294
1624.7777
1298.6389
1623.2924
1299.4360
1299.6279
1625.0419
1298.4324
1620.0432
%
m/z

Mono-C22

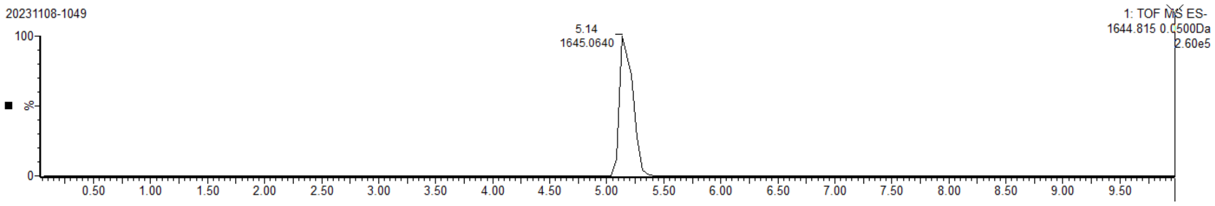
20231108-1049
1: TOF MS ES-
1644.815 0.0500Da
2.60e5
5.14
1645.0640
100
%
0
0.50
1.00
1.50
2.00
2.50
3.00
3.50
4.00
4.50
5.00
5.50
6.00
6.50
7.00
7.50
8.00
8.50
9.00
9.50

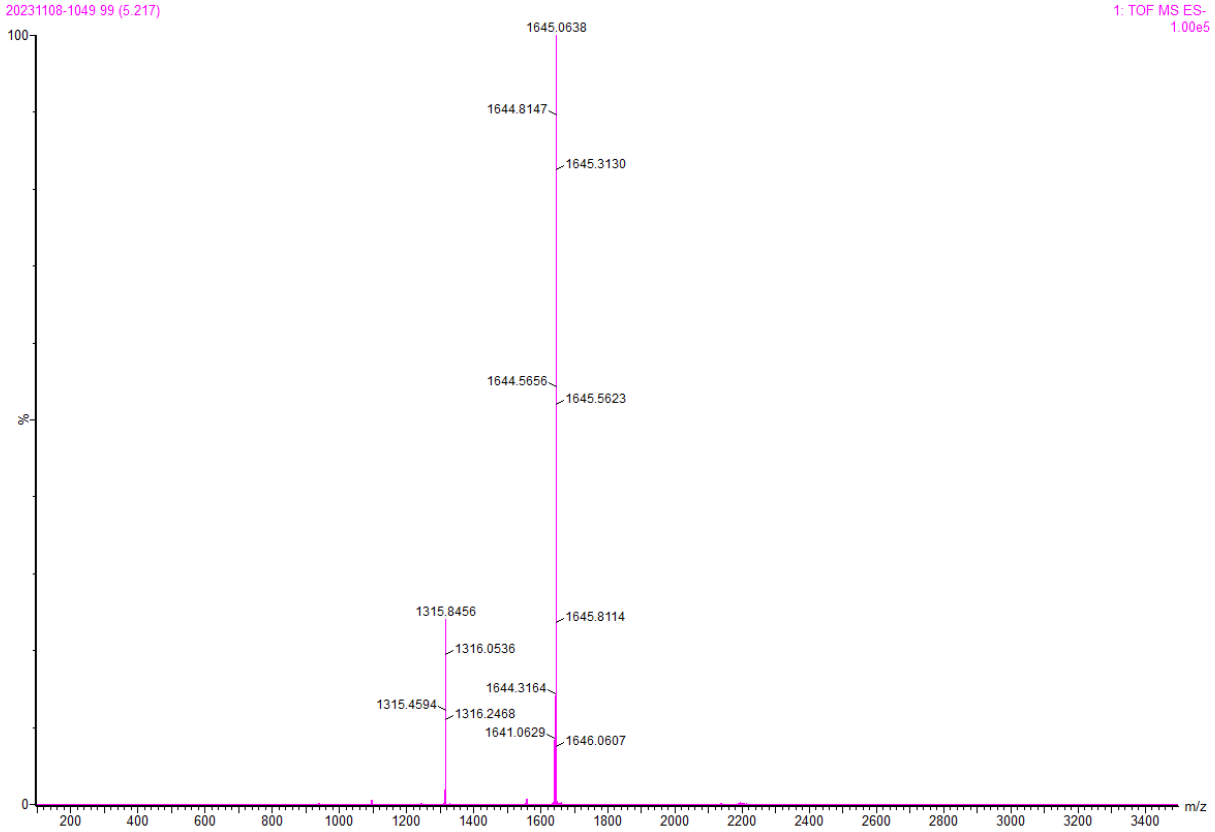
20231108-1049 99 (5.217)
1: TOF MS ES-
1.00e5
1645.0638
1644.8147
1645.3130
1644.5656
1645.5623
1315.8456
1645.8114
1316.0536
1644.3164
1315.4594
1316.2468
1641.0629
1646.0607
100
%
0
200
400
600
800
1000
1200
1400
1600
1800
2000
2200
2400
2600
2800
3000
3200
3400
m/z

Bis-C16

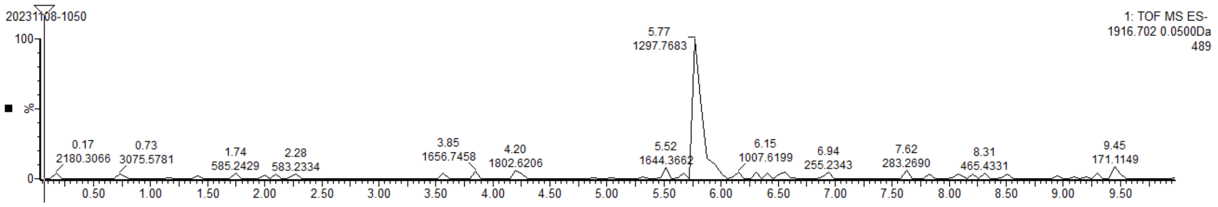
20231108-1050
1: TOF MS ES-
1916.702 0.0500Da
489
100
%
0
5.77
1297.7683
0.17
2180.3066
0.73
3075.5781
1.74
585.2429
2.28
583.2334
3.85
1656.7458
4.20
1802.6206
5.52
1644.3662
6.15
1007.6199
6.94
255.2343
7.62
283.2690
8.31
465.4331
9.45
171.1149
0.50 1.00 1.50 2.00 2.50 3.00 3.50 4.00 4.50 5.00 5.50 6.00 6.50 7.00 7.50 8.00 8.50 9.00 9.50

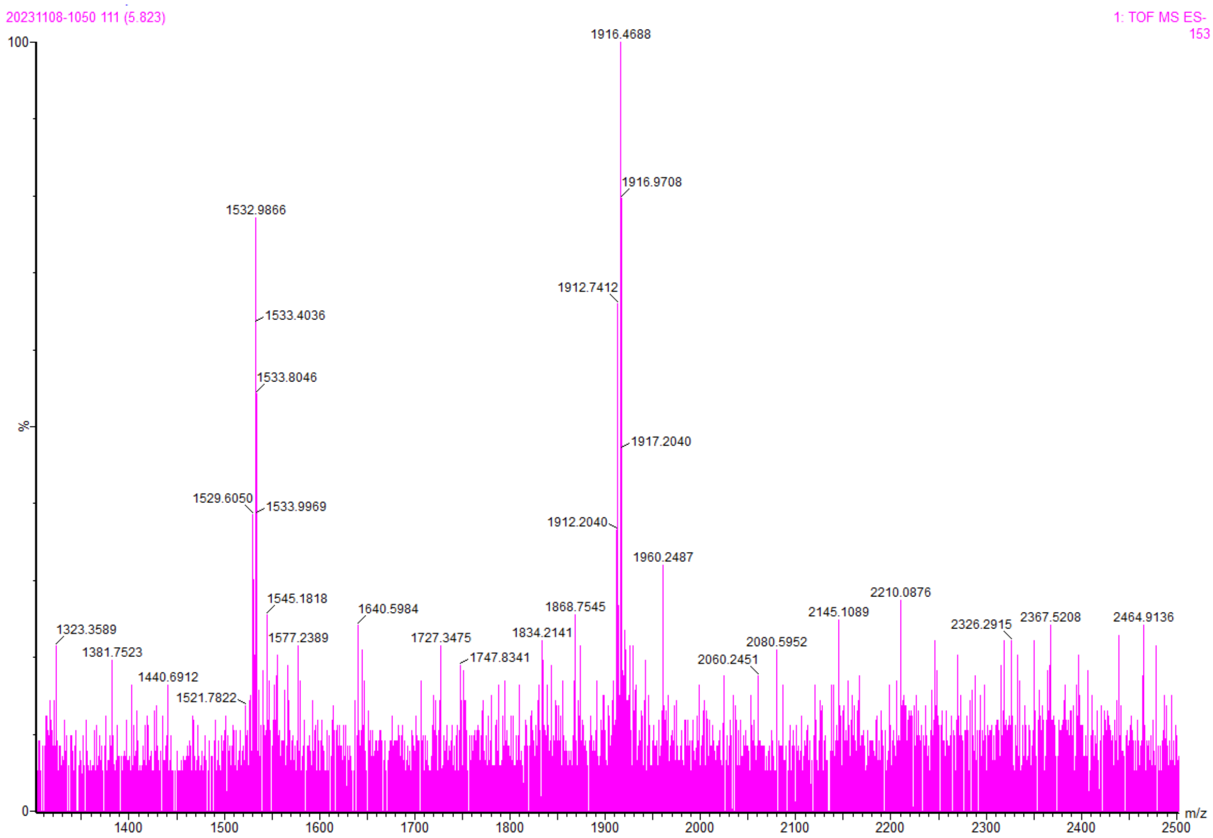
20231108-1050 111 (5.823)
1: TOF MS ES-
153
100
%
0
1916.4688
1916.9708
1532.9866
1912.7412
1533.4036
1533.8046
1917.2040
1529.6050
1533.9969
1912.2040
1960.2487
2210.0876
1545.1818
1640.5984
1868.7545
2145.1089
1323.3589
2326.2915
2367.5208
2464.9136
1577.2389
1727.3475
1834.2141
2080.5952
1381.7523
1747.8341
2060.2451
1440.6912
1521.7822
1400 1500 1600 1700 1800 1900 2000 2100 2200 2300 2400 2500
m/z
